# A switch in jaw form-function coupling during the evolution of mammals

**DOI:** 10.1101/2022.10.06.511001

**Authors:** Z. Jack Tseng, Sergio Garcia-Lara, John J. Flynn, Emily Holmes, Timothy B. Rowe, Blake V. Dickson

## Abstract

The evolutionary shift from a single-element ear, multi-element jaw to a multi-element ear, single-element jaw during the transition to crown mammals marks one of the most dramatic structural transformations in vertebrates. Research on this transformation has focused on mammalian middle-ear evolution, but a mandible comprised of only the dentary is equally emblematic of this evolutionary radiation. Here we show that the remarkably diverse jaw shapes of crown mammals are coupled with surprisingly stereotyped jaw stiffness. This strength-based morphofunctional regime has a genetic basis and allowed mammalian jaws to effectively resist deformation as they radiated into highly disparate forms with markedly distinct diets. The main functional consequences for the mandible of decoupling hearing and mastication were a trade-off between higher jaw stiffness versus decreased mechanical efficiency and speed compared to non-mammals. This fundamental and consequential shift in jaw form-function underpins the ecological and taxonomic diversification of crown mammals.

## Introduction

The radiation of crown group mammals into adaptive zones vacated by the extinction of non-avian dinosaurs led to the Cenozoic Era (66 million years ago to the present) being regarded as the “Age of Mammals”. Among the most dramatic macroevolutionary transformations leading up to the Cenozoic diversification of the ca. 6,500 living species of crown mammals plus their extinct relatives are two novel osteological features: 1. the addition of two new middle ear ossicles, elements from postdentary bones homologous to the articular and quadrate in non-mammal vertebrates, to the one element common to all tetrapods (stapes) and 2. the detachment of postdentary bones from the lower jaw and the related evolution of the temporomandibular joint (TMJ, or the squamosal-dentary joint) (*1, 2*). This second feature is an unreversed mammalian synapomorphy and accomplished the decoupling of hearing acuity on the one hand, and a single-element jaw bone with more precise dental occlusion on the other (*3*–*8*). Study of such morphological transformation and repurposing has fueled paleontological research for nearly two centuries and these shifts are understood to be crucial in enabling mammals to take fuller advantage of a broad range of food sources as active heterotrophs in broad-ranging terrestrial, marine, and aerial environments (*9, 10*).

Although past research focused heavily on the implications of this transition for the mammalian ear, this evolutionary transformation also affected jaw mechanics. Historically, we understand vertebrate feeding systems by characterizing the morphological components as lever arms and force transmission linkages (class III levers and four-bar linkage systems, for example). In this framework, the fulcrum of the jaw mechanical system (jaw joint) underwent three major transformations during the ∼440-million-year history of jawed vertebrates. First, the evolution of an upper-lower jaw articulation in the earliest gnathostomes (*11*) enabled the feeding apparatus to diversify force and speed parameters in a mechanical lever system (*12*). The second reorganization occurred in stem Osteichthyes (*13*), from a quadrate-infragnathal joint to a quadrate-articular joint (*14*); 90% of living vertebrates (∼58,500 species) still possess this class III lever anatomical configuration, attesting to its evolutionary viability and longevity. The third major transformation, from the quadrate-articular joint to a temporomandibular joint in mammals (*15*), further reduced the number of lower jaw elements and intramandibular joints present in this force transmission system (*2*). In recent decades, research on evolutionary transformations of the mammalian jaw has centered on exceptionally preserved fossils documenting the decoupling of hearing and mastication (*16*–*20*); illuminating the biomechanical and kinematic consequences of this important transformation in the loading of the jaw joint and the movements of the now single-boned lower jaw in mammals are major initiatives (*4, 21, 22*). The functional and biomechanical implications of this evolutionary transformation have largely been hypothesized as one of increased mechanical efficiency and strength with the redistribution of joint loading forces and more advantageous in-out lever configurations for jaw adductor muscles. No broad quantitative test of this scenario surrounding the origin of crown Mammalia has yet been performed, however.

Here we demonstrate that shape evolution of the mammalian mandible is characterized by a concomitant increase in morphological disparity but decrease in functional disparity, representing a form-function decoupling. Furthermore, the mammalian jaw transformation represents a trade-off between strength and efficiency; the radiation of mammalian mandibular shape diversity is coupled with higher jaw stiffness but lower jaw speed and mechanical advantage compared to non-mammals. We leverage an integrative array of analyses using elliptic Fourier transforms, experimental mechanical testing, finite element modeling, gene-phenotype-biomechanics association analyses, adaptive landscapes, and phylogenetic comparative methods on a jaw shape dataset of 1,000+ vertebrates (479 mammaliaform genera and 563 non-mammal vertebrate genera; fig. 1, fig. S1, Data S1-2). The mammal dataset includes representatives of all major crown and key stem mammal clades; the non-mammal dataset includes a sampling of fossil and living non-tetrapod vertebrates, amphibians, squamates, archosaurs, turtles, and non-mammalian synapsids. We constructed an ‘all-mandible’ morphospace using the largest 2-D profile shape dataset of jawed vertebrates to date and used it to test a ‘key innovation’ hypothesis of a major mammalian functional morphological evolutionary rate and disparity shift coinciding with a single-element lower jaw. The form-function analyses are grounded in the first principles of jaw lever mechanics (*23*) and the fundamental property of force-speed trade-off in quasi-static lever systems (*24*).

**Figure 1.**
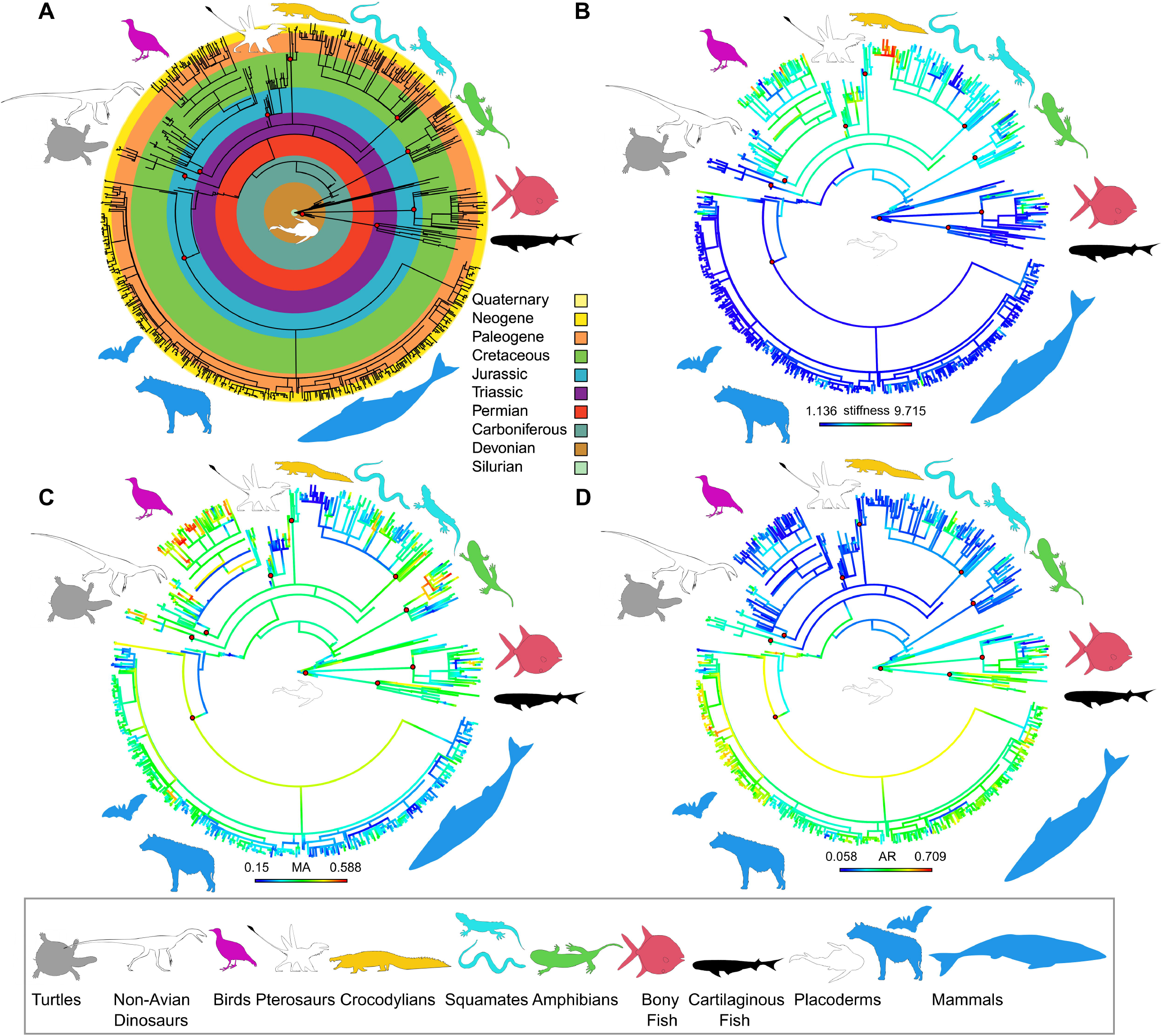
Vertebrate lower jaw shape dataset mapped over a composite phylogeny. (A) Temporal and taxonomic distribution of taxa sampled. (B) Stiffness (maximum deflection in cantilever bending experiment) with ancestral state reconstruction. Lower deflection values imply higher stiffness. (C) Mechanical advantage (MA) heatmap with ancestral state reconstructions. Higher MA values imply more mechanical efficient jaws. (D) Aspect ratio (AR) heatmap with ancestral state reconstructions. Lower AR values imply more streamlined and higher velocity transmission jaws shapes. Silhouettes from phylopic.org (See Data S21 for attributions).

## Material and Methods

### Mandible Shape Analysis

We used 2-D outline shape analysis to quantify vertebrate jaw shape (*12, 25*–*30*)(fig. 2). To capture a range of jaw shapes across vertebrates, we compiled mandible specimen image files (see supplementary materials for detailed sources) and converted them into binary images. The image dataset was imported into the R programming environment for elliptical Fourier analysis using the *Momocs* R package (*31*). A series of smoothing and standardization operations ensured the shapes represented outlines of comparable semi-landmark density (see supplementary materials for additional details). A full generalized Procrustes alignment was performed. The aligned shape data was then subjected to elliptic Fourier transformation with calibrated harmonics search to obtain 99% harmonic power. A principal components analysis (PCA) was then conducted on the transformed data and the first two PC axes used to create a two-dimensional jaw shape morphospace. Within the PC1-PC2 jaw shape morphospace we generated 100 warped theoretical jaw shapes characterizing different regions of the morphospace. Warped jaw shapes were set at regular intervals to cover the X (10 intervals) and Y (10 intervals) axes using the plotting function in R. The resulting warped shapes (hereon called ‘theoretical warp models’) were exported as vector graphics files for both linear biomechanical trait measurements and for 3D bending experiments and finite element simulations (see below).

**Figure 2.**
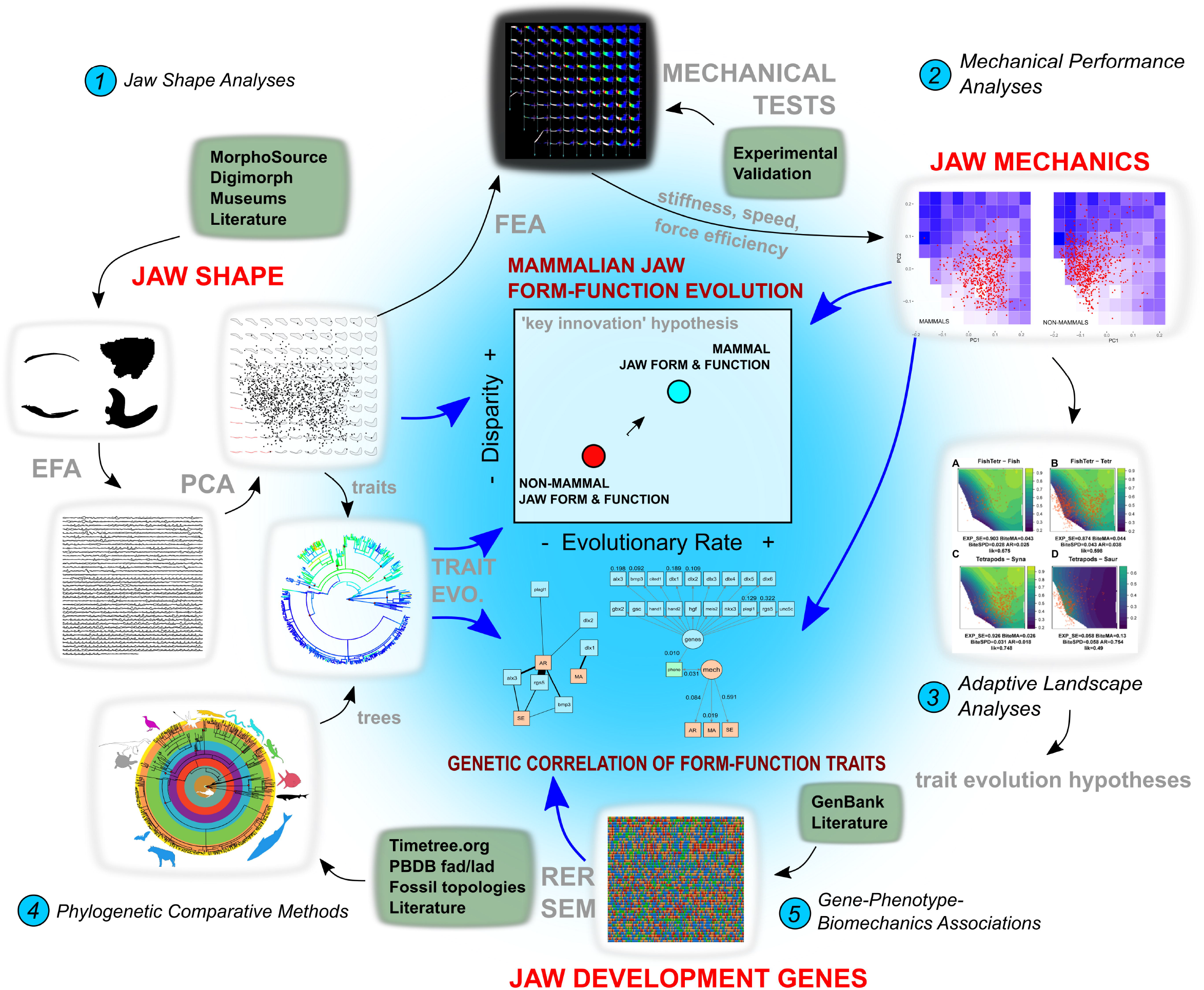
The integrated methodological framework used in this study. The major components of the analyses include 1. Jaw shape analyses using elliptical Fourier analysis (EFA), principal components analysis (PCA), and morphospace visualization. 2. Mechanical performance analyses using finite element analysis (FEA) and cantilever bending tests on theoretical warp models, interpolation of biomechanical traits in individual genera. 3. Adaptive landscape analyses that optimize the relative fitness coefficient among biomechanical traits. 4. Phylogenetic comparative methods that test fit to trait evolution models using interpolate biomechanical traits and a new composite phylogeny for both extant and fossil vertebrate taxa. 5. Relative evolutionary rates (RER) and path analyses using structural equation modeling (SEM) of genotype-phenotype-biomechanics correlations in rates of evolution and in overall similarity.

### Biomechanical Trait Estimation and Validation

Two linear biomechanical traits based on beam theory and lever mechanics principles were measured for each of the 100 theoretical warp models from the morphospace: aspect ratio and mechanical advantage. Mechanical lever ratios have been shown to overestimate force transmission and underestimate velocity transmission (*32*). Therefore, we measured both mechanical advantage and aspect ratio as separate measures of force and velocity, respectively. Jaw aspect ratio is defined as the longest line drawn across the dorsal-ventral axis of each jaw shape outline divided by the longest line drawn across the anterior-posterior axis of the jaw shape. A lower aspect ratio is interpreted as representative of higher jaw displacement or compliance, and higher aspect ratio is an indication of low jaw displacement or compliance. Jaw velocity is estimated as 1/AR; jaws that are elongate will have higher velocity than jaws that are less elongate. Mechanical advantage is defined here as the length of the line between the edge of the outline at the jaw joint to dorsal most point of the coronoid/adductor process divided by the length of the line between the jaw joint and the anterior most point of the jaw shape outline. This ratio measures the relative in-lever arm length of the generalized adductor muscle insertion site at the coronoid process to the out-lever arm length of a hypothetical bite point at the front of the jaw (equivalent to an incisor tooth position in mammals). Both aspect ratio and mechanical advantage values are unit-less and size-free. Measurements used in calculating these ratios were taken on the exported vector graphics file of the 100 theoretical warp models using the linear measurement tool in FIJI (*33*).

We estimated additional biomechanical traits using finite element simulation and validated our simulation dataset using experimental mechanical tests. The 100 theoretical warp models were converted from vector graphics files into extruded (with a thickness of 10 mm) models that were then 3D printed for experimental bending test and digitally solid-meshed for finite element simulations, respectively. For experimental bending tests, hypothetical warp models were printed and mounted on to a custom-fitted vice grip attachment to a Mark-10 ESM1500 electromechanical testing frame (Mark-10, New York, USA). Cantilever bending tests were performed on each plastic jaw model by fixing the jaw joint/grip cube and applying a ventrally directed force at the anterior tip of the model. Two mechanical property values were collected from each 3D printed model experiment: total deflection (up to 10 mm) and strain energy. Deflection is a measure of the stiffness of a particular jaw shape and is measured as the vertical distance travelled by the testing frame during a given experiment. Strain energy is another measurement of stiffness or resistance to deformation, defined as the work done in deforming a material under a particular load, and was estimated as the area under the force-deflection curve by calculating the integral of the curve (in N-mm).

For simulation-based bending tests using the finite element (FE) method, the extruded jaw shape models (in STL format) were imported into Strand7 finite element software (version 2.4.6, Strand7 Pty Ltd, Sydney, Australia) and set to have identical dimensions as the physical models tested using bending experiments. All jaw models were assigned a homogeneous set of material properties suitable for linear static analysis; the elastic (Young’s) modulus was set to 20 GPa, and Poisson’s ratio to 0.3, as in previous FE studies of jaw structures (*12, 34*). Nodal constraints preventing free body movement were applied in each jaw model at both the posterior-dorsal and the posterior-ventral extremes of the jaw shape. A nodal force of 100 N was applied to the anterior most node on the edge of each jaw shape model to simulate a ventrally directed load as in a cantilever bending experiment. The finite element models were solved with a linear static solver using the direct sparse scheme option. After solution, total deflection (in mm) was extracted from each jaw model at the point of the nodal force. Total stored strain energy (in Joules) was obtained from the solution file using the strain energy visualization option.

### Biomechanical Trait Interpolation

We estimated taxon-specific values of aspect ratio (AR), mechanical advantage (MA), total deflection (“stiffness”), and strain energy (another measure of stiffness) using double linear interpolation, implemented in the *interp* R package. Double linear (or bilinear) interpolation sequentially applies linear interpolation to two variables (x, y) in predicting a third (i.e., z). The biomechanical values obtained from linear measurement, experimental, and finite element analyses were set as the z values for the theoretical warp models and used to interpolate z values for per-species biomechanical trait in the taxon-specific shape dataset. The resulting mean interpolated values for each genus was used in all subsequent analyses.

### Jaw Shape and Biomechanical Trait Distribution

To evaluate the assumption that the simplified lateral/2-D profile jaw outlines used in this study retain significant functional morphological signal relative to a null model of randomly distributed shape and biomechanical traits, we conducted a series of non-parametric trait distribution analyses using a combination of bootstrap resampling and Wilcoxon rank sum tests. We first generated 1,000 datasets of 1,000 combinations of PC1 and PC2 values within the jaw shape morphospace as simulated taxa. The biomechanical traits (MA, AR, total deflection, strain energy) were then estimated for simulated taxa using the same interpolation formulae as those used to estimate actual taxa values. Lastly, Wilcoxon rank sum tests of similarity between the distributions of both shape and biomechanical trait values in the simulated shape dataset versus values interpolated for actual taxon shapes were conducted. The hypothesis tested here is that both the PC1-PC2 and biomechanical trait values in the taxonomic dataset collected were significantly different from randomly sampled distributions of those values in the same morphospace, and thus provide support for the 2-D approach.

### Trait Disparity Analyses

To examine potential differences in jaw shape and biomechanics trait disparity between mammals and nonmammals predicted by a ‘key innovation’ hypothesis (fig. 2), we calculated trait disparity using the *dispRity* R package (*35*). In addition, to account for potential biases in disparity metrics introduced by uneven taxonomic sampling and sample sizes, each of the disparity comparisons were both bootstrapped and rarefied. Disparity values were compared between mammals and nonmammals for five morphological and biomechanical traits: (1) PC1 and PC2 values only (reflecting the morphological data used to build the jaw shape morphospace), (2) All PC values from the PCA, (3) Taxon-specific interpolated MA values, (4) Taxon-specific interpolated AR values, and (5) Taxon-specific interpolated, experimentally derived maximum deflection values.

### Body Mass Allometry of Biomechanical and Morphological Traits

To assess the potential correlation between the morphological and biomechanical traits analyzed and body mass, we collected both body mass and jaw adductor muscle mass data from a range of mammals and other vertebrates. Average adult body mass for 411 mammal genera were extracted from the PanTHERIA database (*36*) (Data S19). In addition, total adductor muscle mass (wet weight) were compiled for select vertebrates for which dissection and comparative muscle anatomical data were available in the literature (*37*–*43*) (Data S20). All mass variables were log10 transformed prior to further analysis. The relationship between each of five functional morphological traits (MA, AR, maximum deflection, PC1 shape score, PC2 shape score) and log10 body mass (in grams) were examined using scatterplots. The relationship between log10 body mass and total jaw adductor was plotted and then linearly regressed using the “power trend line” function in Microsoft Excel.

### Adaptive Landscape Analysis

We contextualized the evolutionary and biomechanical optimality of vertebrate mandibles using adaptive landscape analysis (*44, 45*). Briefly, we generated performance surfaces over the main jaw shape morphospace for each biomechanical trait by fitting a second-order polynomial surface using least squares. A combined adaptive landscape was then computed as the summation of individual performance surfaces, each weighed by their relative importance or contribution to overall fitness. In equation form:

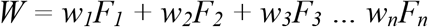

Where *W* is the combined adaptive landscape, *w*_*n*_ is the per-performance surface relative fitness, and *F*_*n*_ is the individual performance surface (*46*). For each examined clade (see below), the value *W* is optimized as the likelihood of combinations of performance surface and relative fitness, under the definition that the total fitness sums to 1 and the variance of all surfaces is set to be equal. Using experimental strain energy (SE), relative bite force (MA), bite speed (1/AR), and jaw displacement (AR) traits as performance variables, adaptive landscapes were optimized for all gnathostomes at five taxonomic levels: non-tetrapods vs tetrapods; between synapsids and sauropsids; and within synapsids, sauropsids, and archosaurs, respectively. The optimized adaptive landscapes were then used as hypotheses for testing models of trait evolution.

### Testing Models of Biomechanical Trait Evolution

To examine evolutionary rates and modes of biomechanical traits (MA, AR, stiffness as measured by strain energy and maximum deflection), we constructed a composite phylogeny including extant and fossil taxa using existing literature (see supplementary materials for additional details). We also estimated branch lengths and their uncertainty over the phylogeny by generating 100 time-tree samples using the Paleobiology Database and the *paleotree* R Package (*47*). Using these 100 time-tree samples, we evaluated the fit of the trees to different models of trait evolution for mechanical advantage, aspect ratio, and stiffness. We evaluated models of trait evolution using the *motmot* R Package to test the results from the adaptive landscape analyses: these models are (1) Brownian motion (BM; a random walk model in which trait variance increases with time since divergence at a constant rate), (2) Ornstein-Uhlenbeck (OU; a BM model with presence of a single optimal trait value or central tendency), and (3) ACDC (“early burst”; model of exponential increase or decrease in evolutionary rate through time).

Furthermore, to assess whether there is a significant change in the rate of biomechanical evolution from non-mammal vertebrates to mammaliaforms during the transition to the mammalian mandible, we modeled biomechanical evolutionary rate heterogeneity using the *traitMEDUSA* method implemented in *motmot*. This rate shift detection method first fits a BM model to the data, then systematically fits two-rate models to each node in the phylogeny and then identifies the best fitting two-rate model using Akaike Information Criterion (AIC). We used this approach to test the hypothesis of a rate shift occurring near the non-mammal to mammal transition but without specifying an exact node, by allowing the fit to be searched over all possible nodes. Because of model misspecification and other biases associated with using individual multivariate shape trait variables such as individual PC axis scores for univariate comparative analysis (*48, 49*), we conducted a separate evolutionary rate analysis for jaw shape using a distance-based method previously proposed for high-dimensional shape and other biological data (*50*). Pairwise between-species differences are used in this approach to estimate multivariate evolutionary rates using a BM model (*50*). We used permutation to obtain statistical significance values in the comparison between mammal and nonmammal shape evolution rates.

To select the best performing biomechanical trait evolution model, we used weights from AICc (AIC corrected for small sample sizes) scores of the three different trait models to identify the preferred model for each of the three biomechanical traits analyzed. To identify locations of potential evolutionary rate change in biomechanical traits on the composite phylogeny, we summarized the results of the two-rate model fitting analyses as a percentage of the 100 tree samples in which a rate increase or decrease is found at a particular node. The same 100 tree samples were used to estimate magnitude and significance of multivariate evolutionary rate differences between mammal and nonmammal jaw shapes.

### Relative Evolutionary Rates of Genes Implicated in Mandible Development

We conducted a relative evolutionary rate analysis using GenBank sequences of genes that have been shown to be involved in normal mandible development in vertebrates. 19 genes were identified from the literature as definitely expressed in the lower jaw. Reference sequences of these genes were obtained using *Homo sapiens* as the query sequence in the BLAST search engine. A summary of the gene families and groups involved in jaw development are included in the supplementary information.

We first assessed the extent to which the candidate genes to be used in relative evolutionary rate analyses display high degrees of covariation with each other. We utilized a phenetic approach (neighbor-joining) to build individual gene trees from FASTA sequences. All possible pairwise comparisons of the 19 jaw development candidate genes are then conducted by pruning each pair of gene trees to the same ingroup and outgroup taxa. The trees then are compared using the entanglement algorithm implemented in the *dendextend* R package (*51*). We then used the algorithms implemented in the *RERconverge* R package for relative evolutionary rate analysis (*52*). Three continuous trait variables from the functional morphological analyses (deflection/strain energy, MA, and AR) for matching taxa in the morphometric dataset were imported and correlated evolutionary rates between the jaw development genes and each of the continuous trait variables were calculated, followed by statistical testing to identify significant gene-morphology correlations.

### Structure Equation Modeling of Gene-Phenotype-Biomechanics Correlations

To examine the correlation between genetic and jaw shape distances among jawed vertebrates, we used structure equation modeling (also known as path analysis or latent variable analysis) to quantify gene sequence covariation with multivariate jaw shape covariation and biomechanical traits. We treated the combined effects of the 19 identified jaw development genes as a genetic latent variable on which the jaw shape, represented by the pairwise Euclidean distance of all principal component shape scores among all taxa for which genetic information is available, depends. We created a second latent variable representing jaw biomechanical performance as measured by aspect ratio, mechanical advantage, and strain energy; this latent variable is correlated to jaw shape in our model, representing structure-function linkage. We fitted this structural equation model to the genetic, shape, and biomechanical data using maximum likelihood in R (*lavaan* package) to estimate path coefficients, which provide measures of the amount of variance explained by individual traits contributing to genetic and biomechanical latent variables, respectively.

## Results

### Mandible Shape Analysis Results

The bivariate plot defined by the first two PC axes (out of a total of 40 PC axes) together represents 59.1% of the total variance in the jaw shape dataset (Fig. 3A). The x-axis (PC1) represents a gradient of shallow (dorsoventrally narrow) jaw shapes with low adductor processes on the negative axis to dorsoventrally deep jaws with high adductor (coronoid) processes on the positive axis. The y-axis (PC2) mainly represents a gradient of jaw shapes with shallow dorsoventral depth at the adductor process on the negative axis to those with deep dorsoventral depth at the adductor process on the positive axis. The lower left-hand corner of the morphospace (negative PC1 and negative PC2 values) contains a region represented by biologically impossible shapes (those with dorsal and ventral outlines contacting or crossing each other); several taxa possess jaw shapes that border those impossible regions, but none fall within it. Such intersecting outlines are a common occurrence in elliptic Fourier analyses and are treated herein as in other studies as geometrically and biologically impossible configurations and removed from further analysis (*12, 53*). Data points in the morphospace are clustered around the origin with a larger spread in the x-axis compared to the y-axis. Generally, the major gnathostome clades overlap each other, and most clades sampled have representative taxa near the origin. Clades such as pterosaurs and squamates contain clusters of taxa that fall near the edge, representing jaw shapes with high velocity (fig. S2).

**Figure 3.**
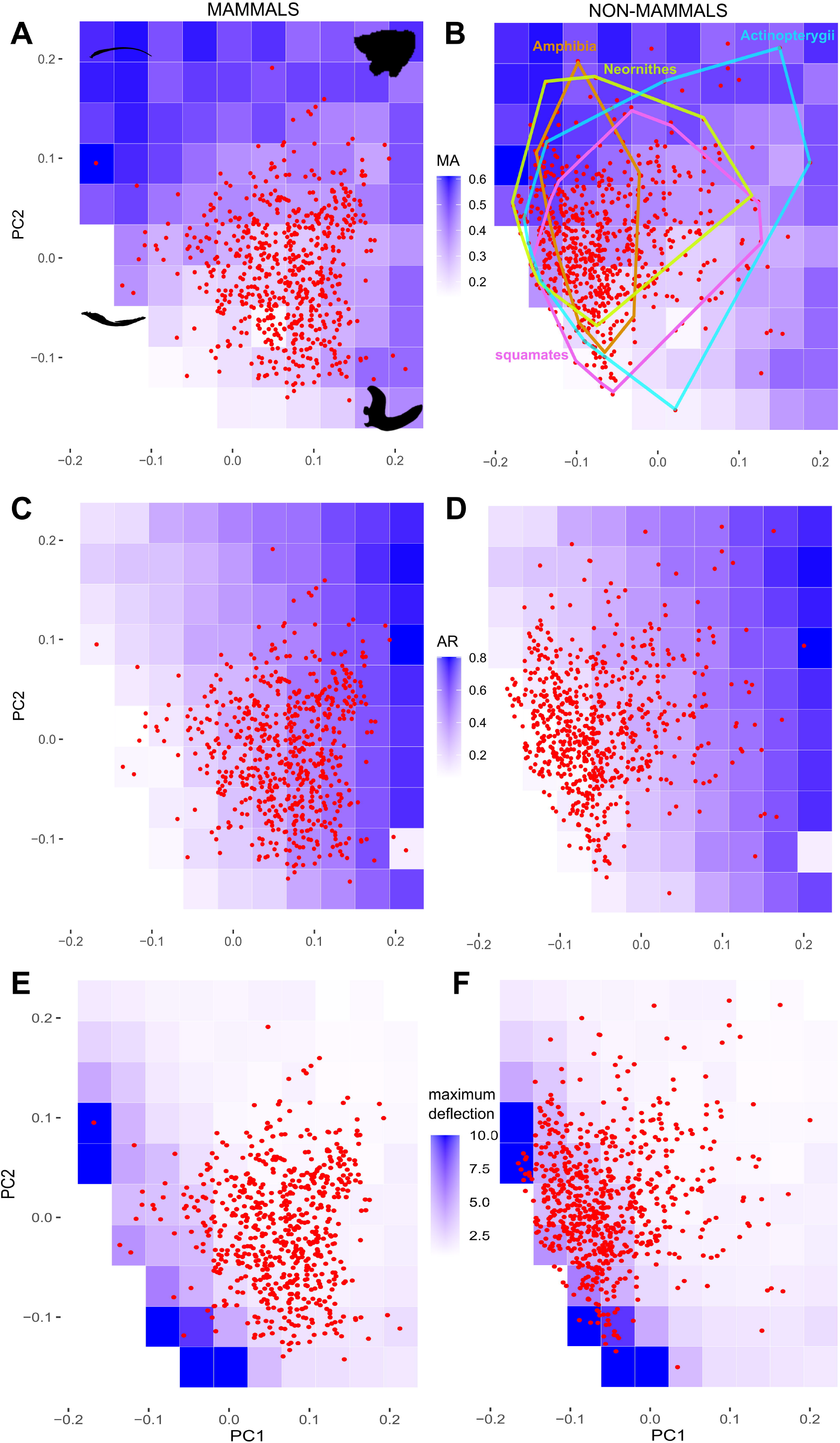
All-mandible morphospace and biomechanical traits. (A) Distribution of mammals in the morphospace (PC1 38.9%, PC2 20.2% variation) of mechanical advantage (MA) represented by background heat maps of values calculated from theoretical warp models. Some representative jaw shapes are shown at the corners of the morphospace. (B) Distribution of non-mammals in the MA morphospace; higher MA implies more efficient jaw force generation. Representative extant clades are indicated by convex hulls to show the extensive overlap of clade jaw shapes in the morphospace. (C) Mammal distribution in jaw aspect ratio morphospace; lower AR implies more streamlined and higher velocity jaw shapes. (D) Non-mammal distribution in jaw AR morphospace. (E) Mammal distribution in jaw stiffness morphospace measured by maximum deflection in cantilever bending experiments; stiffer jaws have lower deflection values. (D) Non-mammal distribution in jaw stiffness morphospace. The theoretical model at (0.2, -0.1) returned unrealistic values in some simulations, so all interpolated taxonomic values were calculated after excluding that particular theoretical model from consideration.

### Biomechanical Trait Estimation and Validation Results

MA and AR values obtained from the linear measurements are included in Data S6. Experimentally measured total deflection and strain energy values were collected for 93 out of 100 warped theoretical jaw shape models; 7 of the theoretical models were biologically impossible and were not tested. Both deflection and strain energy values show a similar pattern across the morphospace: the jaw shapes with the lowest deflection (i.e., stiffest) and strain energy values (i.e., most work efficient) are in the upper right corner of the morphospace (positive PC1 and PC2 values). Displacement and strain energy values both increased with decreasing PC1 and PC2 values. Models bordering biologically impossible forms were flexible and did not fail within the parameters of the cantilever bending experiments, and thus did not have displacement or strain energy values. Given the strong correlation between deflection and strain energy values, we use the experimentally calculated strain energy values and maximum deflection values interchangeably to represent overall structural stiffness in adaptive landscape, trait evolution, relative evolutionary rate, and latent variable path analyses.

Finite element simulations of cantilever bending across the 93 theoretical warp models that represent shapes at regular intervals across the all-mandible morphospace returned highly correlated maximum vertical deflection values between models with intramandibular joints and those without. Outputs of these computational models also correlated strongly to experimentally derived measurements (Movie S1). The lower left-hand edge of the morphospace revealed large discrepancies between simulated and experimental data (fig. S5). Such discrepancies stemmed from jaw shapes with very low aspect ratio (= high jaw closing velocity). Jaw deflection values in experimental data increased exponentially relative to change in aspect ratio towards thinner jaws (Aspect Ratio = 0.5987e^-0.284*Displacement^, *R*^2^ = 0.4728), reflecting the rapid drop in stiffness of the jaw structure as aspect ratio shifts towards higher jaw closing velocity (fig. S4).

### Jaw Shape and Biomechanical Trait Distribution Results

Wilcoxon ranked sum tests between bootstrapped null distributions and actual measured (PC1, PC2) or interpolated (MA, AR, stiffness) taxon-specific trait values showed significant differences. All of the taxon-specific attributes are significantly different from a model of random distribution. There are more occurrences of lower MA and lower AR values in sampled taxa than expected from a random distribution of jaw shapes in the morphospace. There are more higher strain energy and deflection (both indicating lower stiffness) values in sampled taxa compared to randomized jaw shape samples (Fig. S6). PC1 scores (jaw dorsoventral depth and degree of adductor process development) show a distinct bimodal distribution on either side of the origin (zero value) compared to a random distribution model. On the other hand, PC2 scores exhibit a unimodal distribution around the origin rather than a random distribution. Pairwise Wilcoxon tests for between-clade different in biomechanical trait values were conducted (Data S10); however, we caution against overinterpreting comparisons between clades that have very small sample sizes (e.g., turtles, ornithischian dinosaurs). Instead, we focus our discussion on testing mammal versus non-mammal differences.

### Trait Disparity Results

Mammals exhibit significantly lower disparity in combined PC values compared to nonmammals but have significantly higher disparity specifically in PC1 and PC2 values. In addition, mammals have significantly lower disparity in all three biomechanical traits (MA, AR, total deflection/stiffness) from PC1 and PC2 theoretical warp models compared to non-mammals (Fig. S14-15). Mammals exhibit significantly higher aspect ratios (deeper jaws) and higher stiffness (lower maximum deflection values) than non-mammals, but lower mechanical advantage compared to non-mammals (Table 1).

**Table 1.** Summary of biomechanical trait value differences between mammals and non-mammals. MA, mechanical advantage; AR, aspect ratio (higher AR implies lower jaw velocity), ANOVA, one-way analysis of variance. Lower max deflection values indicate higher stiffness.

|  | MA | AR | Stiffness (Max. deflection) |
| --- | --- | --- | --- |
| Mammal Median | 0.28 | 0.41 | 1.52 |
| Nonmammal Median | 0.34 | 0.18 | 3.40 |
| % (Mammal:Nonmammal) | 83.61% | 228.44% | 44.71% |
| p value, Wilcoxon Test | <0.0001 | <0.0001 | <0.0001 |
| Mammal Mean | 0.29 | 0.41 | 1.77 |
| Nonmammal Mean | 0.34 | 0.22 | 3.41 |
| % (Mammal:Nonmammal) | 83.38% | 188.34% | 51.91% |
| p value, ANOVA | <0.0001 | <0.0001 | <0.0001 |

### Body Mass Allometry Results

None of the scatterplots indicated a correlation between body mass and biomechanical and morphological traits; in all analyses the R^2^ was smaller than 0.01. Furthermore, linear regression analysis between body mass and total adductor mass suggests that there is no differential allometry for adductor mass to body mass scaling in the mammals (57 spp), birds (48 spp), or squamates (6 spp) sampled. The squamate slope is lower (i.e., jaw adductor mass increases slower with body mass) than in either birds or mammals, but the small sample size from the squamate dataset precludes a proper statistical comparison (Data S19-S20).

### Adaptive Landscape Analysis Results

Although each taxonomic group returned unique landscapes, pairwise comparisons found significant differences only between synapsids (mammals and their immediate predecessors) and sauropsids (all amniotes more closely related to extant reptiles than to mammals; fig. S27, Table S7), and between turtles and all other sauropsids. Our analyses indicated no differences between non-tetrapods and tetrapods, among archosaurs, among non-testudine sauropsids, or along the grade of synapsids leading to mammals.

Two primary adaptive regimes were responsible for these differences: a strength regime which is primarily adapted for reducing strain energy (i.e., increasing stiffness) and only small contributions from bite force, bite speed and jaw compliance (low AR); and a displacement regime primarily adapted for jaw compliance with moderate contribution from bite force, and small contributions from bite speed and strain energy. The strength regime provides generally high adaptive performance for a majority of morphospace, to which morphologically diverse groups are well adapted. Non-tetrapod vertebrates, stem tetrapodomorphs, turtles, and synapsids are generally broadly distributed through morphospace and are best adapted to the strength regime, while lepidosaurs and archosaurs are generally more restricted in morphospace and are best adapted to the jaw compliance regime (fig. S27). Based on these findings, we hypothesize that an evolutionary rate shift in biomechanical traits occurred in lepidosaurs and archosaurs relative to other vertebrates.

The landscape of stem tetrapodomorphs does not provide any clarity due to limited sample size; although they are “best” adapted for the strength regime, they perform poorly on it (z = 0.51), and nearly as well on a compliance regime. All synapsids were best adapted to a strength regime, and although sample sizes for non-mammalian synapsids were limited, we did find that pelycosaurs performed poorly, with a large increase in cynodonts and then again in mammals. Given the majority of the vertebrates exhibit a strength regime, we hypothesize that strain energy / stiffness evolved via an OU-like process of optimization whereas MA and AR evolution is better explained by a BM process as they were apparently not optimized in most vertebrate groups examined.

### Trait Evolution Mode and Rate Model Fitting Results

In contrast to the hypothesized evolutionary modes based on adaptive landscape analyses, the best fitted trait evolution models for all biomechanical traits (MA, AR, stiffness) across vertebrates are OU models. An OU model is also a better fit than either BM or ACDC when mammal and non-mammal datasets are fitted separately. There is no consistent evidence for evolutionary rate change in biomechanical traits in mammals versus non-mammals. In 68% of trees, laurasiatheres (hedgehogs, bats, hoofed mammals, carnivores) or all crown Mammalia are reconstructed with substantially lower rates of evolution in MA compared to other jawed vertebrates. No clade exhibits substantial increase in MA under a 50% majority rule criterion. In 74% of trees new world monkeys/primates exhibit a two-order increase in aspect ratio (AR) rate evolution. In 69% of trees Toxicofera (snakes, anguimorph lizards, iguanas) or Serpentes exhibit >80% reduction in AR evolutionary rate. In jaw stiffness, 93% of trees show a substantial slow down (<10% of average rate) in Euarchontoglires (lagomorphs, rodents, tree shrews, primates) and a substantial speed up (5-10 times of average rate) for ornithodirans (dinosaurs and pterosaurs).

### Relative Evolutionary Rate and Path Analysis Results

The calculated entanglement values suggest that the genes *CITED1, GBX2, DLX4, MEIS2*, and *RGS5* exhibit the largest differences from other jaw development genes sampled (Data S18). By contrast, *DLX2* shared similar relative taxonomic differences as most other genes sampled. The default set of relative evolutionary rate analyses used no transformations, weighting, or scaling. The results show that evolutionary rate in four genes (*ALX3, PLAGL1, BMP3, DLX5*) are significantly (*p adjusted* < 0.05) correlated to jaw AR evolution. Only *DLX1* is significantly correlated to MA evolutionary rate. Evolutionary rates of three of the analyzed genes (*ALX3, RGS5*, and *BMP3*) are significantly correlated to jaw stiffness evolutionary rate (fig. 4A). A series of sensitivity analyses were conducted using other combinations of transforms, weights, and scales; these analyses provided differential support for the significant correlations identified above (see Table S6 and figures S17-S21).

**Figure 4.**
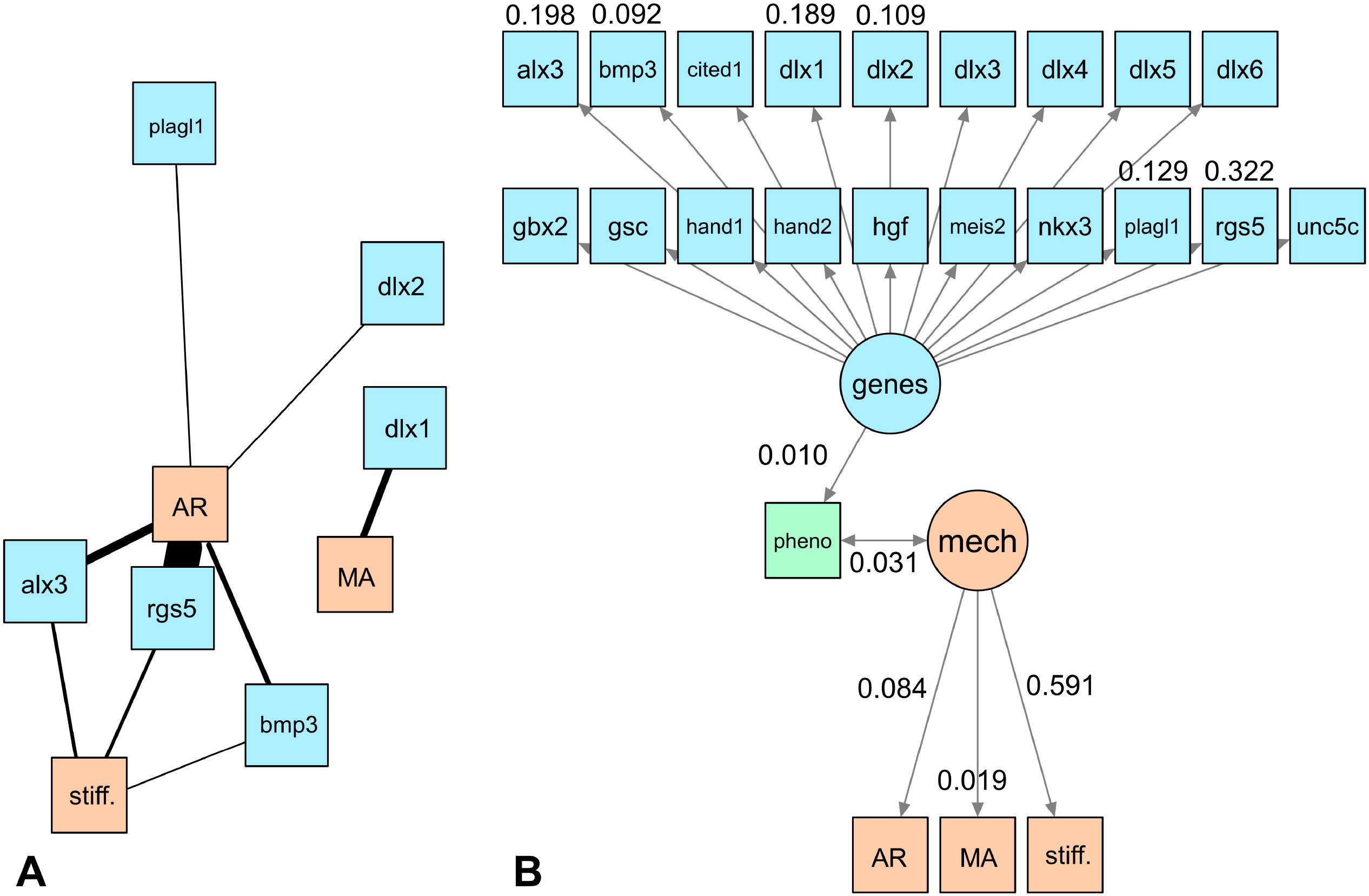
Gene-phenotype-biomechanics association networks. (A) Correlated evolutionary rates between biomechanical (AR, MA, stiffness [measured by maximum deflection in cantilever experiments]) and genes involved in mammalian mandible development portrayed as a network graph; thicker lines indicate higher quantity of significant correlations across sensitivity analyses. (B) Path diagram showing all jaw development genes analyzed as components of a latent variable representing genetic effects on phenotype, and the interaction between phenotype and a latent variable representing biomechanics as measured by AR, MA, and stiffness. Phenotype is measured by multivariate jaw shape data represented by all 40 principal component axes from the elliptic Fourier analysis. Only path coefficients for the genes found to be significantly correlated with biomechanical evolutionary traits are shown. Gene-phenotype-biomechanics associations are weak but statistically significant. Stiffness explains the largest component of variation in phenotype-biomechanics covariation relative to AR and MA.

Relative evolutionary rates of the analyzed genes were plotted over the composite phylogeny (fig. S16) and laurasiatheres are reconstructed as having ancestrally accelerated evolutionary rate for *ALX3*, whereas euarchontoglires exhibit an ancestrally accelerated *PLAGL1* evolutionary rate. Both clades are also reconstructed to have relative low rates of *BMP3* evolutionary rate and no ancestral change in *RGS5* evolutionary rate. No other clade exhibited statistically significant deviations from the overall evolutionary rates. The genes found to be significantly correlated with biomechanical trait evolutionary rates show the same relative contribution to genotype-phenotype correlation in the path analysis (fig. 4B). There is a significant but weak correlation between phenotype and biomechanics, with stiffness (maximum deflection) contributing the most to this association (path coefficient = 0.59), followed by AR (0.08) and MA (0.02).

## Discussion

Functional morphological analyses of the vertebrate lower jaw indicate the presence of biomechanical decoupling and trade-off associated with the morphological diversification of the mammalian single-element mandible. Mammalian and non-mammalian jaws are distinguished by a switch in morphological and biomechanical trait disparity relationships. Bootstrapped and rarefied disparity metrics show statistically significant increases in mandible shape disparity in mammals (fig. S14-S15; Data S10). On the other hand, mammalian jaw MA, AR, and stiffness were significantly lower in disparity, even after accounting for uneven taxon sampling. Furthermore, mammals exhibit significantly higher jaw stiffness, but lower jaw efficiency (MA) and speed (high AR / low jaw velocity), compared to non-mammals (Table 1). This combination of high morphological shape disparity, low biomechanical trait disparity, and biomechanical trade-off characterizes many-to-one structure-function mapping, one of the principal manifestations of a redundant mapping of morphology to performance in biological systems. Other prime examples of such complex mapping have been extensively documented in actinopterygian jaw systems (*54*). Presence of this many-to-one structure-function relationship in mammals indicates that they independently capitalized on this form-function complex to diversify in jaw shape while maintaining stereotyped, stiffness-optimized jaw biomechanical characteristics.

Appearance of the single-element mandible in mammals did not correspond to a shift in the optimized adaptive landscape regime for jaw biomechanics. Jawed vertebrates collectively are optimized by a strength regime in an adaptive landscape framework; jaw stiffness is high in both non-tetrapods and tetrapods (fig. S13, S24). Mammals and non-mammalian synapsids maintain this strength regime (Table S1), but sauropsids (all tetrapods more closely related to extant reptiles than to mammals) have an optimized adaptive landscape characterized by an aspect ratio (AR) regime focused more on compliant and faster-closing jaws (fig. S24). During their evolution, crown mammals, with their single-element jaw bones, optimized on the existing strength regime within the adaptive landscape by further elevating jaw stiffness (fig. 3E-F). However, the lower biomechanical trait disparity associated with the mammalian mandible within this strength regime reinforces the presence of functional constraints despite high shape disparity and stiffness; many differently shaped mammalian jaws exhibit similar mechanical traits. Furthermore, the significant increase in jaw stiffness comes at the expense of lowered jaw efficiency and speed (Table 1). Taken together, these findings indicate that the morphological outcome of crown mammalian radiation was an evolutionary exploration of jaw shape space within prescribed bounds of the tetrapod adaptive landscape, characterized by further augmentation and stereotyping of high jaw strength. This mammalian pattern is partially convergent with the earliest gnathostomes, which evolved under a similar regime of form-function decoupling (*12*); however, the diverse dental morphologies and environments in which mammals have radiated represent an unique context for interpreting the consequences of such decoupling.

Given the findings of the adaptive landscape analyses, we hypothesized that speed (AR) and efficiency (MA) functions evolved without evolutionary rate shifts during the transition from mammals to non-mammals; we also predicted that jaw stiffness experienced an evolutionary rate shift between mammals and non-mammals. We tested these hypotheses by modeling AR, MA, and stiffness trait evolution using a variable rate shift model and found no evidence for evolutionary rate shifts in AR or MA at the transition to mammals. We also did not observe a rate shift in stiffness in the transition to mammals, although the fitted model suggests Euarchontoglires (the crown mammal clades including lagomorphs, rodents, primates, tree shrews) experienced a substantial slowdown in stiffness evolutionary rate, in part reflecting the decreased disparity in stiffness values. These findings indicate that the OU-type jaw stiffness evolution suggested by the adaptive landscape analysis and supported by trait evolution modeling did not couple with modified evolutionary rates of change in jaw stiffness in all mammals; rather, the increase in jaw shape disparity in mammals occurred without modifying the jaw stiffness evolutionary tempo or mode already observed in their non-mammalian relatives. Furthermore, although adaptive landscape analyses optimized stiffness/strength as the primary regime and stiffness data best fit an OU process of evolution, both AR and MA trait evolution fit an OU model as well; this is interpreted as evidence of a biomechanical trade-off where all three traits may be evolving toward their respective functional optima, but strength/stiffness appears to be the principal trait that mammals and non-sauropsid vertebrates in general optimized at the expense of efficiency and speed. This observation suggests stiffness could be an universally optimized functional trait across vertebrate clades and is evidenced by its importance in mechanically demanding feeding tasks as disparate as suction feeding (*55*) and bone-cracking (*56*).

Relative gene-biomechanics evolutionary rate analyses demonstrate that the narrower range of biomechanical trait values that distinguish mammalian jaws from non-mammalian jaws is genetically correlated (fig. 4; fig. S16-S21). In comparison to evolutionary rates of 19 candidate genes known to play key roles in the development of the mandible within the first pharyngeal/branchial arch (BA1), biomechanical trait evolutionary rates are statistically correlated to genetic variation across living mammal species (fig. 4A). Stiffness evolutionary rates are significantly correlated with *ALX3, RGS5*, and *BMP3*. Mechanical advantage (correlated to *DLX1*) and jaw velocity (correlated to *BMP3, PLAGL1, ALX3, RGS5, DLX2*) trait values are also found to be genetically linked. Nested expression of the *DLX* gene family by cranial neural crest cells is associated with normal mandible development and jaw adductor formation, and is also known to regulate the downstream expression of *PLAGL1, ALX3, RGS5*, and other genes during normal jaw development (*57*). The association of these genes to biting-specific biomechanical traits provide evidence of heritability in jaw force, velocity transmission, and bending stiffness variation (*58, 59*). Given this finding, we speculate that these and potentially other candidate genes involved in craniofacial development were targets of selection during the Mesozoic origin of the single-element mandible in mammals.

Path analyses show that the genes with the highest degree of correlated evolutionary rates to biomechanical variables also contribute the highest amount of variance towards the gene-jaw shape covariance, which is statistically significant but weak (fig. 4B; path coefficient = 0.01). Among the biomechanical traits, strain energy explains the most variance in the jaw shape-biomechanics correlation among the three measured traits (SE path coefficient = 0.591 compared to 0.084 for AR and 0.019 for MA). These findings suggest that the low disparity of jaw stiffness in mammals is in part related to correlated rates of biomechanical evolution with jaw development genes, but such correlation is only weakly associated with overall jaw shape. We speculate that genetically encoded structure-function linkages for jaw stiffness is localized in certain regions of the jaw rather than to the overall shape. Future comparative evolutionary developmental studies would provide insights into the expression patterns of these biomechanically relevant candidate genes.

The presence of many-to-one form-function mapping in the mammalian mandible suggests that the single-element lower jaw may be an example of a “fly in a tube” model of evolution, whereby potential for evolutionary disparity and/or rate shifts within a trait complex (in this case, jaw biomechanics) is constrained in certain regions of a morphospace and shaped by genotypic and/or phenotypic integration (*60*). The absence of a significant rate shift or trait evolution model shift in biomechanical traits along the mammalian transition to a dentary-squamosal joint is consistent with expectations of this model; the single-element mammal jaw represents integrated biomechanical trait complexes that traverse a smaller part of the functional space compared to non-mammals ((*60*); fig. 4B). Such a pattern of high shape disparity coupled with lower biomechanical disparity explains previous findings of redundant and complex form-function mapping of mammalian feeding systems at lower taxonomic rank levels (*61*) and suggests a revised conceptual framework for understanding crown mammalian jaw evolution (fig. 5). These results do not support an interpretation whereby simplified, single-element jaws substantially improved the force and velocity transmission lever system in terms of MA and AR during the radiation of crown mammals, but do support the interpretation that stiffness is the principal axis of mammalian jaw adaptation. The differences between a single-element mammalian jaw and multi-element non-mammalian jaw are thus best characterized by a switch in form-function coupling rather than a wholesale ‘key innovation’. Multi-element systems, such as in squamate skulls, can be highly integrated morphologically but are nevertheless able to respond to dietary selective pressure at when examined at lower taxonomic levels (*62*); this phenomenon is consistent with the diversity of adaptive landscape regimes across sauropsid clades (fig. S24). Specific adaptive advantages for a single-element jaw may lie instead in enabling diverse jaw shapes optimized for stiffness to support the extremely diverse dentitions which are characteristic of mammals. Stiffness-optimized mandibular shapes may also facilitate evolutionary transformations in other, potentially less stereotyped components of the feeding system such as muscle physiology, bone microstructure, or other morphological traits with higher functional or mechanical sensitivity (*63*).

**Figure 5.**
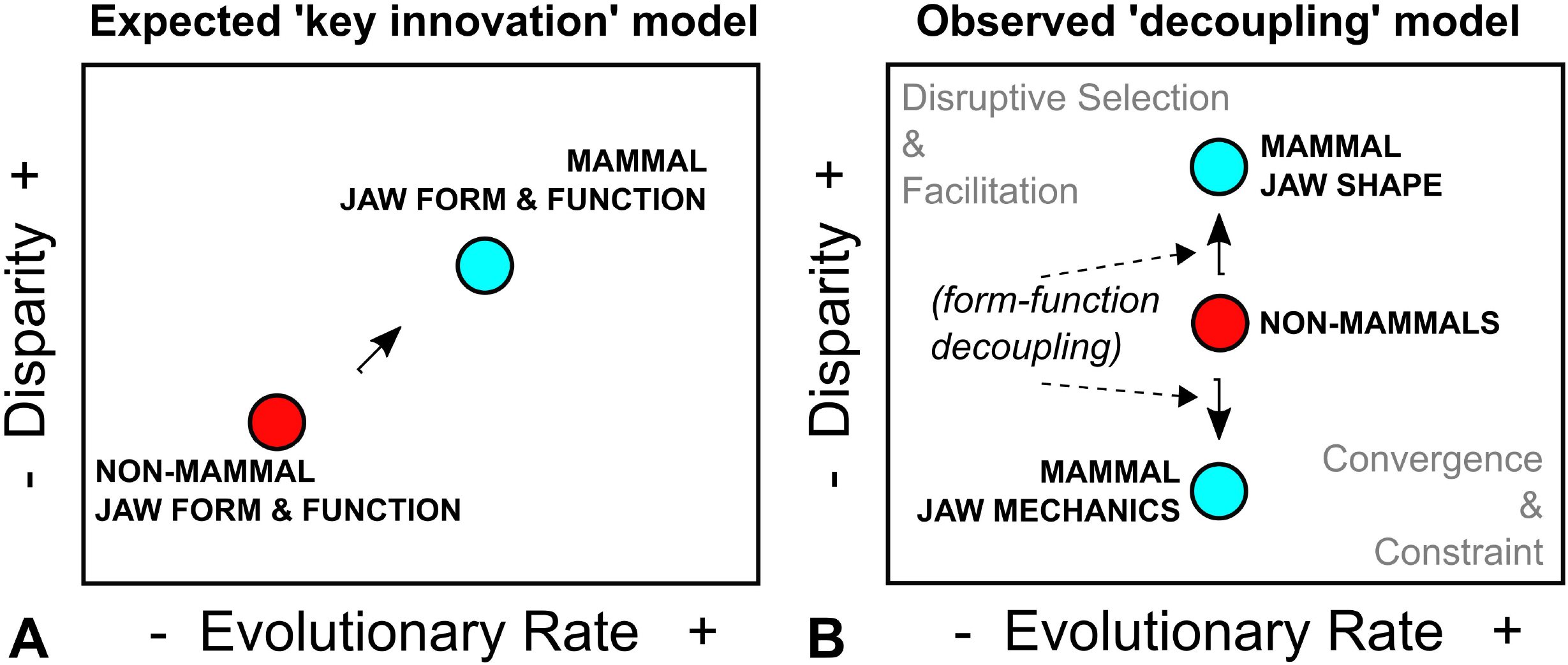
Evolutionary rate-disparity framework for mammalian jaw form-function evolution. (A) Graphical representation of a ‘key innovation’ type of model for non-mammal to mammal jaw transformation. In this model, the single-element jaw of mammals is associated with increase in both jaw shape and function disparity and evolutionary rate as a result of an adaptive radiation at the origin of mammals. (B) The observed rate-disparity shift from non-mammals to mammals is one of for-function decoupling, whereby mammal jaw shapes increased in disparity and mammal jaw biomechanical traits decreased in disparity relative to non-mammals. Modified from (*60*).

Body size is an important organismal trait underlying many biological scaling relationships (*64*); in the context of mammalian jaws, body mass is poorly correlated with morphological and biomechanical traits. This size independence applies both to shape (PC1 and PC2) and biomechanical traits (fig. S22-S23; Data S19-S20). This suggests that during their evolutionary radiation, crown mammals of different sizes comparably explored the jaw morphospace region within which they are found in terms of mandible biomechanics. This finding corroborates the recent observation that ecomorphological signals overtake those of body size in mammalian jaws (*65*) and provides new evidence that despite the extraordinarily large size range represented by living mammals, from 2g bats to almost 200,000 kg whales, the same genetically correlated, shape-induced mechanical constraints exist.

Although it is likely like functional morphological variation in 3-D bone material properties and shape are not fully captured by the 2-D mandibular profiles examined in this study, the significantly non-random distributions of both shape and biomechanical traits in our dataset suggest critical functional signals are still present in 2-D data. It is likely that MA, AR, and stiffness measures taken from 2-D jaw shapes only partially capture the biomechanical characteristics of 3-D jaw function, but any systematic errors in estimating those traits using 2-D data should not bias the interpretations made herein, unless there are significant differences in how well 2-D represent 3-D structures from clade to clade. On the other hand, the extent to which jaw shapes of different vertebrate clades overlap each other in morphospace may change if 3-D shapes are used; this may be particularly important for taxa in which there are substantial mediolateral morphological and biomechanical differences. Despite these potential shortcomings, the important role of dorsoventral kinetics and kinematics of mammalian and all gnathostome vertebrate jaws (the jaw adductor muscle vectors are principally oriented in the parasagittal plane (*66, 67*) and creates rotation mainly around the mediolateral axis and translation mainly in that parasagittal plane) indicates that 2-D projections are both informative and time-efficient sources of relevant functional morphology variation.

Other factors to be considered in future research and which may interact with the stiffness-governed form-function regime identified in this study include those associated with mammalian jaw movements about the dorsoventral axis (yaw), which increased in mechanical advantage relative to the movement about a mediolateral axis in crown mammals (*5, 68*). Research on the origin of the vertebrate dentition and its evolutionary interactions with the jaw bones housing it also suggest that mechanical environment at the tooth-bone junction is an key region of jaw system evolution (*69*); the importance of tooth-bone interactions became greatly amplified in mammals as a complex neuroanatomical system evolved along with an occlusal dentition and mastication (*70*). Furthermore, a single-bone mandible that is stiff regardless of shape may permit larger jaw closing forces via effective resistance to deformation caused by higher absolute muscle contraction magnitudes as mammals evolved increasingly large body sizes through the Cenozoic, starting during the immediate aftermath of the end-Cretaceous extinctions (*71*–*73*). Evolutionary encephalization and epigenetic effects from growing a larger brain likely exerted a cascading influence on mammalian skull configuration, especially in the auditory, olfactory, and masticatory systems (*74*–*76*). Within crown mammals, marsupial mammals appear to exhibit lower jaw disparity relative to placentals given the former’s developmental constraints (*77*); future investigations into within-mammal variations should clarify how developmental differences among mammalian clades further modulate this broad, stereotyped form-function configuration.

Although mandibular evolution was once thought principally to have enabled mammals to innovate in feeding mechanical versatility and efficiency over those of non-mammal vertebrates, the new data presented here across a wide sample of vertebrates clearly document that stereotyped stiffness in the mandible was a pre-existing condition underpinning the diverse jaw shapes expressed in crown mammals. A further optimized stiffness regime across diverse mammal jaw shapes evolved at the expense of efficiency and speed, but such a regime may have served as the hotbed for the radiation of dental morphologies and masticatory strategies which exemplify mammalian biodiversity. These new findings should encourage further investigations into how crown mammals radiated into new adaptive zones and became ecologically dominant in various ecosystems during the Cenozoic by capitalizing on a fundamentally modified and stereotyped jaw form-function configuration.

## Acknowledgements

J. Liu, C. Marshall, K. Padian, J. Hoeflich, P. Kloess, R. Dudley, R. Full, M. Koehl, C. Williams, participants of IB222 (Fall 2021), and members of the Functional Anatomy and Vertebrate Evolution Laboratory (all UCB) provided insightful discussion and comments on earlier versions of this work. A. Tseng assisted with biomechanical simulations. G. Bowers contributed jaw images. P. Sudmant (UCB) helped troubleshoot RERconverge. J. Demboski (DMNS), K. Zyskowski (YPM), M. Aja (MCZ), P. Myers (UMMZ), S. Wagner (CUMV) provided approval for use of images in their care. P.D. Polly, S. Lautenschlager, and thematic issue editor L. Fostowicz provided detailed and constructive comments that greatly improved the rigor of this study. Reviews from a declined NSF CAREER proposal (ZJT) provided motivation for this project. This project was funded in part by National Science Foundation Grants DBI-2128146 (ZJT) and DEB-1257572 (ZJT, JJF).

## Author contributions

Conceptualization: ZJT

Methodology: ZJT, SGL, EH, BVD

Data Collection: ZJT, SGL, JJF, TBR, EH

Investigation: ZJT, SGL, EH, BVD

Visualization: ZJT, SGL, EH, BVD

Funding acquisition: ZJT, JJF

Project administration: ZJT

Supervision: ZJT

Writing – original draft: ZJT, SGL, BVD

Writing – review & editing: All co-authors

## Data and materials availability

All data are available in the main text or the supplementary materials.

## Supplementary Materials

Additional Material and Methods Details

Appendix S1 to S7

Figs. S1 to S24

Tables S1 to S6

Movie S1

Data S1 to S21

## References

1. J. Wang, J. R. Wible, B. Guo, S. L. Shelley, H. Hu, S. Bi, A monotreme-like auditory apparatus in a Middle Jurassic haramiyidan. Nature. 590, 279–283 (2021).

2. H. R. Barghusen, J. A. Hopson, Dentary-Squamosal Joint and the Origin of Mammals. Science (80-.). 168, 573–575 (1970).

3. F. Mao, Y. Hu, C. Li, Y. Wang, M. Hill Chase, A. Smith, J. Meng, Integrated hearing and chewing modules decoupled in a Cretaceous stem therian mammal. Science (80-.). 367, 305–308 (2020).

4. S. Lautenschlager, P. G. Gill, Z.-X. Luo, M. J. Fagan, E. J. Rayfield, The role of miniaturization in the evolution of the mammalian jaw and middle ear. Nature. 561, 533– 537 (2018).

5. D. M. Grossnickle, The evolutionary origin of jaw yaw in mammals. Sci. Rep. 7, 45094 (2017).

6. D. A. Reed, J. Iriarte-Diaz, T. G. H. Diekwisch, A three dimensional free body analysis describing variation in the musculoskeletal configuration of the cynodont lower jaw. Evol. Dev. 18, 41–53 (2016).

7. A. W. Crompton, F. A. Jenkins, Mammals from Reptiles: A Review of Mammalian Origins. Annu. Rev. Earth Planet. Sci. 1, 131–155 (1973).

8. K. A. Kermack, F. Mussett, H. Rigney, The lower jaw of Morganucodon. Zool. J. Linn. Soc. 53, 87–175 (1973).

9. A. W. Crompton, The Evolution of the Mammalian Jaw. Evolution (N. Y). 17, 431–439 (1963).

10. D. M. Bramble, Origin of the Mammalian Feeding Complex: Models and Mechanisms. Paleobiology. 4, 271–301 (1978).

11. R. Cerny, M. Cattell, T. Sauka-Spengler, M. Bronner-Fraser, F. Yu, D. M. Medeiros, Proc. Natl. Acad. Sci., in press, doi:10.1073/pnas.1009304107.

12. W. Deakin, P. S. Anderson, W. den Boer, T. Smith, H. J. J., M. Rücklin, P. C. Donoghue, E. Rayfield, Increasing morphological disparity and decreasing optimality for jaw speed and strength during the radiation of jawed vertebrates. Sci. Adv. 8, eabl3644 (2022).

13. E. C. Olson, Jaw mechanisms: Rhipidiatians, Amphibians, Reptiles. Am. Zool. 1, 205–215 (1961).

14. M. Zhu, P. Ahlberg, Z. Pan, Y. Zhu, T. Qiao, W. Zhao, L. Jia, J. Lu, A Silurian maxillate placoderm illuminates jaw evolution. Science (80-.). 354, 334–336 (2016).

15. A. S. Romer, Cynodont Reptile with Incipient Mammalian Jaw Articulation. Science (80-.). 166, 881–882 (1969).

16. C.-F. Zhou, B.-A. Bhullar, A. Neander, T. Martin, Z.-X. Luo, New Jurassic mammaliaform sheds light on early evolution of mammal-like hyoid bones. Science (80-.). 365, 276–279 (2019).

17. Z.-X. Luo, J. A. Schultz, E. G. Ekdale, in Evolution of the Vertebrate EarJ: Evidence from the Fossil Record, J. A. Clack, R. R. Fay, A. N. Popper, Eds. (Springer International Publishing, Cham, 2016; https://doi.org/10.1007/978-3-319-46661-3_6), pp. 139–174.

18. M. Thomas, L. Zhe-Xi, Homoplasy in the Mammalian Ear. Science (80-.). 307, 861–862 (2005).

19. Z.-X. Luo, Q.-J. Meng, D. M. Grossnickle, D. Liu, A. I. Neander, Y.-G. Zhang, Q. Ji, New evidence for mammaliaform ear evolution and feeding adaptation in a Jurassic ecosystem. Nature. 548, 326–329 (2017).

20. Z.-X. Luo, S. Gatesy, N. Shubin, Mandibular and dental characteristics of Late Triassic mammaliaform Haramiyavia and their ramifications for basal mammal evolution. Proc. Natl. Acad. Sci. 112, E7101–E7109 (2015).

21. G. L. Benevento, R. B. J. Benson, M. Friedman, Patterns of mammalian jaw ecomorphological disparity during the Mesozoic/Cenozoic transition. Proc. R. Soc. B Biol. Sci. 286, 20190347 (2019).

22. N. M. Morales-García, P. G. Gill, C. M. Janis, E. J. Rayfield, Jaw shape and mechanical advantage are indicative of diet in Mesozoic mammals. Commun. Biol. 4, 242 (2021).

23. P. C. Wainwright, Ecological Explanation through Functional Morphology: The Feeding Biology of Sunfishes. Ecology. 77, 1336–1343 (1996).

24. M. J. McHenry, There is no trade-off between speed and force in a dynamic lever system. Biol. Lett. 7, 384–386 (2011).

25. J. S. Crampton, Elliptic Fourier shape analysis of fossil bivalves: some practical considerations. Lethaia. 28, 179–186 (1995).

26. J. J. Hill, M. N. Puttick, T. L. Stubbs, E. J. Rayfield, P. C. J. Donoghue, Evolution of jaw disparity in fishes. Palaeontology. 61, 847–854 (2018).

27. C. A. Navarro, E. Martin-Silverstone, T. L. Stubbs, Morphometric assessment of pterosaur jaw disparity. R. Soc. Open Sci. 5, 172130 (2022).

28. J. Schaeffer, M. J. Benton, E. J. Rayfield, T. L. Stubbs, Morphological disparity in theropod jaws: comparing discrete characters and geometric morphometrics. Palaeontology. 63, 283–299 (2020).

29. S. Ferson, F. J. Rohlf, R. K. Koehn, Measuring Shape Variation of Two-Dimensional Outlines. Syst. Zool. 34, 59–68 (1985).

30. P. E. Lestrel, Method for analyzing complex two-dimensional forms: Elliptical Fourier functions. Am. J. Hum. Biol. 1, 149–164 (1989).

31. V. Bonhomme, S. Picq, C. Gaucherel, J. Claude, Momocs: Outline Analysis Using R. J. Stat. Softw. 56, 1–24 (2014).

32. M. W. Westneat, A biomechanical model for analysis of muscle force, power output and lower jaw motion in fishes. J. Theor. Biol. 223, 269–281 (2003).

33. J. Schindelin, I. Arganda-Carreras, E. Frise, V. Kaynig, M. Longair, T. Pietzsch, S. Preibisch, C. Rueden, S. Saalfeld, B. Schmid, J.-Y. Tinevez, D. J. White, V. Hartenstein, K. Eliceiri, P. Tomancak, A. Cardona, Fiji: an open-source platform for biological-image analysis. Nat. Methods. 9, 676–682 (2012).

34. Z. J. Tseng, D. F. Su, X. Wang, S. C. White, X. Ji, Feeding capability in the extinct giant Siamogale melilutra and comparative mandibular biomechanics of living Lutrinae. Sci. Rep. 7, 15225 (2017).

35. T. Guillerme, dispRity: A modular R package for measuring disparity. Methods Ecol. Evol. 9, 1755–1763 (2018).

36. K. E. Jones, J. Bielby, M. Cardillo, S. A. Fritz, J. O’Dell, C. D. L. Orme, K. Safi, W. Sechrest, E. H. Boakes, C. Carbone, PanTHERIA: a specieslJlevel database of life history, ecology, and geography of extant and recently extinct mammals: Ecological Archives E090lJ184. Ecology. 90, 2648 (2009).

37. V. Schaerlaeken, A. Herrel, P. Aerts, C. F. Ross, The functional significance of the lower temporal bar in Sphenodon punctatus. J. Exp. Biol. 211, 3908–3914 (2008).

38. A. Herrel, D. Adriaens, W. Verraes, P. Aerts, Bite performance in clariid fishes with hypertrophied jaw adductors as deduced by bite modeling. J. Morphol. 253, 196–205 (2002).

39. M. van der Meij, R. Bout, Scaling of jaw muscle size and maximal bite force in finches. J. Exp. Biol. 207, 2745–2753 (2004).

40. A. Herrel, A. De Smet, L. F. Aguirre, P. Aerts, Morphological and mechanical determinants of bite force in bats: do muscles matter? J. Exp. Biol. 211, 86–91 (2008).

41. J. M. G. Perry, A. Hartstone-Rose, C. E. Wall, The Jaw Adductors of Strepsirrhines in Relation to Body Size, Diet, and Ingested Food Size. Anat. Rec. 294, 712–728 (2011).

42. A. Hartstone-Rose, I. Hertzig, E. Dickinson, Bite Force and Masticatory Muscle Architecture Adaptations in the Dietarily Diverse Musteloidea (Carnivora). Anat. Rec. 302, 2287–2299 (2019).

43. A. Hartstone-Rose, E. Dickinson, A. R. Deutsch, N. Worden, G. A. Hirschkorn, Masticatory muscle architectural correlates of dietary diversity in Canidae, Ursidae, and across the order Carnivora. Anat. Rec. 305, 477–497 (2022).

44. B. V Dickson, J. A. Clack, T. R. Smithson, S. E. Pierce, Functional adaptive landscapes predict terrestrial capacity at the origin of limbs. Nature. 589, 242–245 (2021).

45. B. V Dickson, S. E. Pierce, Functional performance of turtle humerus shape across an ecological adaptive landscape. Evolution (N. Y). 73, 1265–1277 (2019).

46. P. D. Polly, C. T. Stayton, E. R. Dumont, S. E. Pierce, E. J. Rayfield, K. D. Angielczyk, Combining geometric morphometrics and finite element analysis with evolutionary modeling: towards a synthesis. J. Vertebr. Paleontol. 36, e1111225 (2016).

47. D. W. Bapst, paleotree: an R package for paleontological and phylogenetic analyses of evolution. Methods Ecol. Evol. 3, 803–807 (2012).

48. J. C. Uyeda, D. S. Caetano, M. W. Pennell, Comparative Analysis of Principal Components Can be Misleading. Syst. Biol. 64, 677–689 (2015).

49. D. C. Adams, M. L. Collyer, Multivariate Phylogenetic Comparative Methods: Evaluations, Comparisons, and Recommendations. Syst. Biol. 67, 14–31 (2018).

50. D. C. Adams, Quantifying and Comparing Phylogenetic Evolutionary Rates for Shape and Other High-Dimensional Phenotypic Data. Syst. Biol. 63, 166–177 (2014).

51. T. Galili, dendextend: an R package for visualizing, adjusting and comparing trees of hierarchical clustering. Bioinformatics. 31, 3718–3720 (2015).

52. A. Kowalczyk, W. K. Meyer, R. Partha, W. Mao, N. L. Clark, M. Chikina, RERconverge: an R package for associating evolutionary rates with convergent traits. Bioinformatics. 35, 4815–4817 (2019).

53. G. Diaz, Contour recognition of complex leaf shapes. PLoS One. 12, e0189427 (2017).

54. P. C. Wainwright, M. E. Alfaro, D. I. Bolnick, C. D. Hulsey, Many-to-One Mapping of Form to Function: A General Principle in Organismal Design?1. Integr. Comp. Biol. 45, 256–262 (2005).

55. P. C. Wainwright, M. D. McGee, S. J. Longo, L. Patricia Hernandez, Origins, Innovations, and Diversification of Suction Feeding in Vertebrates. Integr. Comp. Biol. 55, 134–145 (2015).

56. Z. J. Tseng, Testing adaptive hypotheses of convergence with functional landscapes: a case study of bone-cracking hypercarnivores. PLoS One. 8, e65305 (2013).

57. J. Jeong, X. Li, R. J. McEvilly, M. G. Rosenfeld, T. Lufkin, J. L. R. Rubenstein, Dlx genes pattern mammalian jaw primordium by regulating both lower jaw-specific and upper jaw-specific genetic programs. Development. 135, 2905–2916 (2008).

58. D. M. J., L. Thomas, R. J. L. R., Specification of Jaw Subdivisions by Dlx Genes. Science (80-.). 298, 381–385 (2002).

59. É. Heude, K. Bouhali, Y. Kurihara, H. Kurihara, G. Couly, P. Janvier, G. Levi, Jaw muscularization requires Dlx expression by cranial neural crest cells. Proc. Natl. Acad. Sci. 107, 11441–11446 (2010).

60. R. N. Felice, M. Randau, A. Goswami, A fly in a tube: Macroevolutionary expectations for integrated phenotypes. Evolution (N. Y). 72, 2580–2594 (2018).

61. Z. J. Tseng, J. J. Flynn, Structure-function covariation with nonfeeding ecological variables influences evolution of feeding specialization in Carnivora. Sci. Adv. 4, eaao5441 (2018).

62. A. Watanabe, A.-C. Fabre, R. N. Felice, J. A. Maisano, J. Müller, A. Herrel, A. Goswami, Proc. Natl. Acad. Sci., in press, doi:10.1073/pnas.1820967116.

63. M. M. Muñoz, P. S. L. Anderson, S. N. Patek, Mechanical sensitivity and the dynamics of evolutionary rate shifts in biomechanical systems. Proc. R. Soc. B Biol. Sci. 284, 20162325 (2017).

64. M. LaBarbera, Analyzing Body Size as a Factor in Ecology and Evolution. Annu. Rev. Ecol. Syst. 20, 97–117 (1989).

65. D. M. Grossnickle, Feeding ecology has a stronger evolutionary influence on functional morphology than on body mass in mammals. Evolution (N. Y). 74, 610–628 (2020).

66. A. Datovo, R. P. Vari, The Jaw Adductor Muscle Complex in Teleostean Fishes: Evolution, Homologies and Revised Nomenclature (Osteichthyes: Actinopterygii). PLoS One. 8, e60846 (2013).

67. J. M. Ziermann, Diaz Raul E.,, Diogo, Rui,, Heads, jaws, and musclesJ: anatomical, functional, and developmental diversity in chordate evolution (Cham, 2019).

68. B.-A. S. Bhullar, A. R. Manafzadeh, J. A. Miyamae, E. A. Hoffman, E. L. Brainerd, C. Musinsky, A. W. Crompton, Rolling of the jaw is essential for mammalian chewing and tribosphenic molar function. Nature. 566, 528–532 (2019).

69. V. Valéria, C. Donglei, T. Paul, J. Zerina, E. Boris, B. Henning, A. P. Erik, Marginal dentition and multiple dermal jawbones as the ancestral condition of jawed vertebrates. Science (80-.). 369, 211–216 (2020).

70. T. B. Rowe, J. H. B. T.-E. N. (Second E. Kaas, Ed. (Academic Press, London, 2020), pp. 263–319.

71. J. Alroy, Cope’s Rule and the Dynamics of Body Mass Evolution in North American Fossil Mammals. Science (80-.). 280, 731–734 (1998).

72. J. Baker, A. Meade, M. Pagel, C. Venditti, Adaptive evolution toward larger size in mammals. Proc. Natl. Acad. Sci. 112, 5093–5098 (2015).

73. O. Bertrand, S. Shelley, T. Williamson, J. Wible, S. Chester, J. Flynn, L. Holbrook, T. Lyson, J. Meng, I. Miller, H. Püschel, T. Smith, M. Spaulding, Z. Tseng, S. Brusatte, Brawn before brains in placental mammals after the end-Cretaceous extinction. Science (80-.). 376, 80–85 (2022).

74. T. Rowe, Brain heterochrony and origin of the mammalian middle ear. Mem. Calif. Acad. Sci. 20, 71–95 (1996).

75. T. Rowe, Coevolution of the Mammalian Middle Ear and Neocortex. Science (80-.). 273, 651–654 (1996).

76. T. B. Rowe, T. E. Macrini, Z.-X. Luo, Fossil Evidence on Origin of the Mammalian Brain. Science (80-.). 332, 955–957 (2011).

77. A.-C. Fabre, C. Dowling, R. Portela Miguez, V. Fernandez, E. Noirault, A. Goswami, Functional constraints during development limit jaw shape evolution in marsupials. Proc. R. Soc. B Biol. Sci. 288, 20210319 (2021).

