## Supplementary Materials for "A switch in jaw form-function coupling during the evolution of mammals"

**This PDF file includes:**

Additional Material and Methods Details
Appendices S1 to S7
Figs. S1 to S24
Tables S1 to S6
Caption and DOI for Movie S1
Captions and DOI for Data S1 to S21

**Other Supplementary Materials for this manuscript include the following:**

Movie S1
Data S1 to S21

### Additional Material and Methods Details

Mandible Image Source

We compiled a dataset of vertebrate lower jaw images from several major sources. Published images were modified from (1), two specimens; (2), one specimen; (3), three specimens; (4), one specimen; (5), 7 specimens; (6), 34 specimens; (7), one specimen; (8), one specimen; (9), 23 specimens; (10), one specimen; (11), 16 specimens; (12), 1 specimen; (13), one specimen; (14), 7 specimens; (15), 138 specimens; (16), one specimen; (17), 7 specimens; (18), three specimens; (19), four specimens; (20), 7 specimens; (21), 31 specimens; (22), 9 specimens; (23), 4 specimens; (24), one specimen; (25), one specimen; (26), 2 specimens; (27), 30 specimens; (28), 46 specimens; (29), 28 specimens; (30), 4 specimens; (31), 4 specimens; (32), 13 specimens; (33), 1 specimen; (34), 1 specimen; (35), 1 specimen; (36), 2 specimens; (37), 60 specimens; (38), 14 specimens; (39), 1 specimen; (40), 10 specimens; (41), 16 specimens; (42), 1 specimen; (43), one specimen; (44), 13 specimens; (45), 7 specimens.

Additionally, we used screenshots from online collection databases (with proper permissions and/or attributions in Data S1, S2): Denver Museum of Natural Sciences (in Arctos database) with permission from J. Demboski, two specimens; Animal Diversity Web, University of Michigan Museum of Zoology, 227 specimens; Morphosource, 259 specimens; Digimorph, 140 specimens; Museum of Vertebrate Zoology, University of California, Berkeley (in Arctos database), 44 specimens; Understanding Evolution, University of California Museum of Paleontology, one specimen; Museum of Southwestern Biology (in Arctos database), one specimen; Yale Peabody Museum, 29 specimens; Museum of Comparative Zoology, Harvard University, 39 specimens; Cornell Lab of Ornithology, 1 specimen.

Lastly, we incorporated new data (for details see Data S1) from 45 specimens in the American Museum of Natural History.

Additional Shape Analysis Details

We used 2-D outline shape analysis to quantify vertebrate jaw shape (*12*, *25–30*)(fig. 2). To capture a range of jaw shapes across vertebrates, we compiled mandible specimen image files (see supplementary materials for detailed sources) and processed them in GIMP (GNU Image Manipulation Program, https://www.gimp.org/) version 2.10.24 to 2.10.30. We used the 'magic wand' function in GIMP to select and remove background pixels that are not part of the mandible. Given the lack of homologous dental positions at the level of vertebrates, we also digitally removed the dentition from all dentate taxa. Each image was then cropped to a minimal size that contained the entire mandible. Contrast of the image was then maximized, following by a binary threshold filter to render background pixels white and mandible pixels, black. Where necessary, internal regions within each mandible were filled with black pixels so that white pixels only represented background surrounding each mandible. Each processed mandible image was saved as a JPEG file.

Next, the mandible profile dataset was imported into the R programming environment for elliptical Fourier analysis using the *Momocs* R package (*31*). To ensure consistent image quality across different data sources, all images were first smoothed using the 'coo_smoothcurve' function in *Momocs* for ten iterations. The smoothed curves then were first centered on the origin, then aligned according to their long axes. Next, each curve representing a jaw shape was interpolated so that the outline is represented by 500 equidistant points. At this stage, two sensitivity analyses were conducted to evaluate the arbitrary choice of choosing 500 interpolated points to represent each jaw shape; the interpolation was repeated with 1,000 points, then with 100 points, and all downstream analyses from this step to the full shape morphospace were replicated. If the resulting morphospace showed no visible or substantial differences between the interpolation sample sizes, the original 500-point interpolation dataset was deemed adequate for portraying the jaw shapes. The elliptic Fourier transform data objects generated at different stages in this process are included in Data S3 and S4.

Prior to performing superimposition analysis and constructing the shape morphospace, we choose four fixed landmarks to limit the amount of rotation that occurs during superimposition to maintain comparable jaw shape orientations for downstream analyses. The four landmarks chosen for superimposition were (1) anterior most point of the jaw shape, (2) dorsal most point of the jaw shape (= 'coronoid process'), (3) the location of the jaw joint (temporomandibular joint in mammals and quadrate-articular joint in other vertebrates), and (4) the ventral most point on the ventral border of the mandibular body. Next, a full generalized Procrustes alignment was performed. The aligned shape data was then subjected to elliptic Fourier transformation with calibrated harmonics search to obtain 99% harmonic power. A principal components analysis (PCA) was then conducted on the transformed data and the first two PC axes used to create a two-dimensional jaw shape morphospace.

The list of specimens included in the initial morphospace analysis are shown in Table S1 and Data S1. The bivariate plot defined by the first two PC axes (out of a total of 40 PC axes) together represents 59.1% of the total variance in the jaw shape dataset (Fig. 3A). The x-axis (PC1) represents a gradient of shallow (dorsoventrally narrow) jaw shapes with low adductor processes on the negative axis to dorsoventrally deep jaws with high adductor (coronoid) processes on the positive axis. The y-axis (PC2) mainly represents a gradient of jaw shapes with shallow dorsoventral depth at the adductor process on the negative axis to those with deep dorsoventral depth at the adductor process on the positive axis. The lower left-hand corner of the morphospace (negative PC1 and negative PC2 values) contains a region represented by biologically impossible shapes (those with dorsal and ventral outlines contacting or crossing each other); several taxa possess jaw shapes that border those impossible regions, but none fall within it. Such intersecting outlines are a common occurrence in elliptic Fourier analyses and are treated herein as in other studies as geometrically and biologically impossible configurations and removed from further analysis (*12*, *62*). Data points in the morphospace are clustered around the origin with a larger spread in the x-axis compared to the y-axis. Generally, the major gnathostome clades overlap each other, and most clades sampled have representative taxa near the origin. Clades such as pterosaurs and squamates contain clusters of taxa that fall near the edge, representing jaw shapes with high velocity (fig. S3). The complete PC score matrix is included in Data S5.

Additional Biomechanical Analysis Details

We estimated additional biomechanical traits using finite element simulation and validated our simulation dataset using experimental mechanical tests. The 100 theoretical warp models were converted from vector graphics files into extruded (i.e., with thickness) models that were then 3D printed for experimental bending test and digitally solid-meshed for finite element simulations, respectively. Model extrusion was performed by converting the binary jaw shape vector graphics files into line segments in Inkscape (versions 1.0, 1.1; https://inkscape.org/) and exporting the resulting segments as DXF files. The DXF files then were imported into OpenSCAD (https://openscad.org) and converted to STL files with a standardized length of 30 mm and lateral thickness of 10 mm.

For experimental bending tests, all jaw shape STL files were printed in a polylactic acid (PLA) thermoplastic material on a Ultimaker S5 FDM (fused deposition modeling) 3D printer with an infill of 100% (solid). To permit a cantilever test to be performed with the posterior jaw (equivalent to jaw joint position) fixed, a cubic test grip (10 x 10 x 10 mm) was designed and added to the extruded jaw shape models in Geomagic Wrap 2021 (3D Systems, South Carolina, USA)(Movie S1). All models were printed with this grip and were individually mounted on to a custom-fitted vice grip attachment to a Mark-10 ESM1500 electromechanical testing frame (Mark-10, New York, USA). Cantilever bending tests were performed on each plastic jaw model by fixing the jaw joint/grip cube and applying a ventrally directed force at the anterior tip of the model. A physiologically conservative loading rate of 1 mm/s was used, and the test was conducted until the vertical deflection reached 10 mm or when the model failed, whichever condition occurred first. Force (in Newtons) and deflection (in mm) were collected at a sampling rate of 20 Hz using MESURgage Plus (Mark-10, New York, USA).

Two mechanical property values were collected from each 3D printed model experiment: total deflection (up to 10 mm) and strain energy. Deflection is a measure of the stiffness of a particular jaw shape and is measured as the vertical distance travelled by the testing frame during a given experiment. Strain energy is another measurement of stiffness or resistance to deformation, defined as the work done in deforming a material under a particular load, and in this study was estimated as the area under the force-deflection curve by calculating the integral of the curve (in N-mm).

For simulation-based bending tests using the finite element (FE) method, the extruded jaw shape models (in STL format) were imported into Strand7 finite element software (version 2.4.6, Strand7 Pty Ltd, Sydney, Australia) and set to have identical dimensions as the physical models tested using bending experiments. All jaw models were assigned a homogeneous set of material properties suitable for linear static analysis; the elastic (Young's) modulus was set to 20 GPa, and Poisson's ratio to 0.3, as in previous FE studies of jaw structures (*12*, *34*). Nodal constraints preventing free body movement were applied in each jaw model at both the posterior-dorsal and the posterior-ventral extremes of the jaw shape; the presence of two constraints were necessary to fully constrain movement during the solution of this static scenario. These constraints simulated the vice grip positions in the experimental setup. A nodal force of 100 N was applied to the anterior most node on the edge of each jaw shape model to simulate a ventrally directed load as in a cantilever bending experiment. Because we are interested in the relative performance of the theoretical warp models that represent different locations in the shape morphospace, the absolute magnitude of the load does not need to match those of the physical experiments. The finite element models were solved with a linear static solver using the direct sparse scheme option. After solution, total deflection (in mm) was extracted from each jaw model at the point of the nodal force. Total stored strain energy (in Joules) was obtained from the solution file using the strain energy visualization option.

Two sets of finite element simulations were run on the theoretical warp models to account for variation in the presence of intramandibular joints in different jawed vertebrates. Cranial kinesis is a well-established phenomenon extensively observed in non-tetrapod vertebrates (*35–41*), but other vertebrates such as snakes (*42*) and to some extent, birds, also may exhibit different degrees of kinesis in the jaw system. To examine the effects of intramandibular joint presence on the estimated biomechanical variables, a single joint running dorsoventrally across the entire mid-section on each theoretical jaw model was introduced. The joints were given a different set of mechanical properties based on (*43*). All other boundary conditions and material properties were kept constant as in the solid model simulations. The experimentally derived deflection and strain energy values are regressed against the finite element outputs of both no-joint and intra-mandibular joint model datasets to verify the validity of the finite element model results.

The experimental validation data are included in Data S9, and R script is included in Appendix S2. The relative strain magnitudes of the warped jaw FE models are shown in fig. S4. The calculated deflection and strain energy values are included in the table Data S7, S8, S10. The interpolated biomechanical traits (MA, AR, maximum deflection) are included in Data S11. See Appendix S1 for R script.

Trait Disparity Analyses Details

To examine potential differences in jaw shape and biomechanics trait disparity between mammals and nonmammals predicted by a 'key innovation' hypothesis, we calculated trait disparity using the *dispRity* R package (*44*). The metric used for calculating disparity within each categorical group is the median of distances between pairwise taxonomic values; the metric used for testing differences between the categorical groups is deviation from the centroid. The statistical test used to compare group disparity values is the Wilcoxon test. In addition, to account for potential biases in disparity metrics introduced by uneven taxonomic sampling and sample sizes, each of the disparity comparisons were both bootstrapped and rarefied using the *boot.matrix()* function. Disparity values were compared between mammals and nonmammals for five morphological and biomechanical traits: (1) PC1 and PC2 values only (reflecting the morphological data used to build the jaw shape morphospace), (2) All PC values from the PCA, (3) Taxon-specific interpolated MA values, (4) Taxon-specific interpolated AR values, and (5) Taxon-specific interpolated, experimentally derived maximum deflection values. The R script used for these analyses are included in Appendix S5. Additionally, we also tested for significant differences in mean and median values of the biomechanical traits (MA, AR, stiffness or maximum deflection) between mammals and non-mammals using parametric (ANOVA) and non-parametric (Wilcoxon rank sum) methods.

Models of Biomechanical Trait Evolution Details

To examine evolutionary rates and modes of biomechanical traits (MA, AR, stiffness as measured by strain energy and maximum deflection), we constructed a composite phylogeny including extant and fossil taxa using existing literature. Because of model misspecification and other biases associated with using individual multivariate shape trait variables such as individual PC axis scores for univariate comparative analysis (*56*, *57*), we conducted a separate evolutionary rate analysis for jaw shape using a distance-based method previously proposed for high-dimensional shape and other biological data (*58*). A composite phylogeny of extant species included in the jaw shape dataset was generated using a custom text file uploaded to timetree.org. The resulting phylogeny was downloaded as a NEWICK file including both topological and branch length information (Data S13). The lack of a comprehensive phylogeny including all living and fossil vertebrate taxa necessitated a manually constructed composite tree that incorporated fossil taxa. We first compiled information about the interrelationships of taxa within extinct clades using a literature search. We then inserted fossil taxa into the main extant vertebrate phylogeny based on proposed phylogenetic relationships from the literature.

In addition to generating a composite topology for the maximum number of analyzed taxa possible, it was also necessary to estimate branch lengths over the phylogeny to estimate evolutionary rates. We calculated branch lengths on the composite phylogeny using stratigraphic occurrence ranges (First Appearance Datum [FAD] and Last Appearance Datum [LAD]) reported in the Paleobiology Database. The topology and stratigraphic ranges were combined using the *paleotree* R Package (*59*). To account for uncertainty in the sampling of FAD and LAD with regard to the actual divergence age at a given internal node, we used the *cal3TimePaleoPhy* function in *paleotree* to scale branch lengths using sampling rate estimated from the FAD-LAD list. A sample of 100 time-trees were generated using this approach, and all subsequent phylogenetic comparative analyses were done on the whole sample of 100 trees to account for time-scale uncertainty in datasets that include fossils.

Using the 100 time-tree samples, we evaluated the fit of the trees to different models of trait evolution for mechanical advantage, aspect ratio, and stiffness. We evaluated models of trait evolution using the *motmot* R Package to test the results from the adaptive landscape analyses: these models are (1) Brownian motion (BM; a random walk model in which trait variance increases with time since divergence at a constant rate), (2) Ornstein-Uhlenbeck (OU; a BM model with presence of a single optimal trait value or central tendency), (3) ACDC ("early burst"; model of exponential increase or decrease in evolutionary rate through time. Furthermore, to assess whether there is a significant change in the rate of biomechanical evolution from non-mammal vertebrates to mammaliaforms during the transition to the mammalian mandible, we modeled biomechanical evolutionary rate heterogeneity using the *traitMEDUSA* method implemented in *motmot*. This rate shift detection method first fits a BM model to the data, then systematically fits two-rate models to each node in the phylogeny and then identifies the best fitting two-rate model using Akaike Information Criterion (AIC). We used this approach to test the hypothesis of a rate shift occurring near the non-mammal to mammal transition but without specifying an exact node, by allowing the fit to be searched over all possible nodes. For the multivariate jaw shape traits, we tested for rate differences between mammal and non-mammal PC scores using a distance based method for high-dimensional data implemented in the *geomorph* R package; pairwise between-species differences are used in this approach to estimate multivariate evolutionary rates using a BM model (*58*). We used permutation to obtain statistical significance values in the comparison between mammal and nonmammal shape evolution rates.

To select the best performing biomechanical trait evolution model, we used weights from AICc (AIC corrected for small sample sizes) scores of the three different trait models to identify the preferred model for each of the three biomechanical traits analyzed. To identify locations of potential evolutionary rate change in biomechanical traits on the composite phylogeny, we summarized the results of the two-rate model fitting analyses as a percentage of the 100 tree samples in which a rate increase or decrease is found at a particular node. The same 100 tree samples were used to estimate magnitude and significance of multivariate evolutionary rate differences between mammal and nonmammal jaw shapes.

In addition to accounting for the branch length uncertainty in the composite phylogeny, we also examined the effect of uneven or biased taxonomic sampling on the resulting interpretations of trait evolution rate and mode. To accomplish this, we resampled the jaw shape dataset using a bootstrap approach. First, we tallied the approximate number of extant species currently recognized in each of the major clades of jawed vertebrates, including Actinopterygii, Sarcopterygii, Chondrichthyes, Amphibia, Squamata, Rhynchocephalia, Testudines, Loricata, Neornithes, and Mammalia. This was done by tabulating species diversity values from several clade-specific databases (including the California Academy of Sciences Catalog of Fishes, AmphibiaWeb, American Society of Mammalogists Mammal Diversity Database, International Ornithologists' Union World Bird List, and the Reptile Database)(Data S14). Based on the relative percentage of the total number of species in these groups represented by each respective clade, we resampled (without replacement) the extant species clades to have proportionally similar diversity to their percentage of jawed vertebrate diversity. The resampled dataset was set to 95 total extant taxa based on setting the number of taxa sampled in the most undersampled clade (Actinopterygii, 40 sampled taxa or 5.25% of total dataset) to be the corresponding percentage of the new sample (47.38%) they are supposed to represent based on their representation among all living jawed vertebrate species. In this resampling scheme, all fossil taxa were included in the resampled datasets without consideration for their respective taxonomic diversity, given the difficulty of estimating fossil diversity or treating them as representative of their actual macroevolutionary diversity.

A total of 100 taxonomic resampled datasets were generated, and for each taxonomic resampled dataset a total of 100 time-tree samples were generated using the method described for the main composite phylogeny above. This sample of 10,000 trees were then subjected to the same trait evolution analyses as for the main composite phylogeny.

Sources of topological information used in constructing the composite phylogeny

Fossil taxa (Data S12): phylogenetic placement and topological hypotheses were compiled from the literature (46, 47, 56–65, 48, 66–75, 49, 76–85, 50, 86–95, 51, 96–105, 52, 106–115, 53, 116–119, 54, 55) with aid from the Paleobiology Database for target publications containing data on interrelationships. Additionally, topological relationships were compared and verified with additional data from review papers for pterosaurs (49), tetrapodomorphs (120, 121), coelacanths (122), some Chondrichthyes (123), non-avian theropod dinosaurs (117, 124, 125), early actinopterygians (126), early turtles (127), early gnathostomes (99), ichthyosaurs (128, 129), cynodonts (130), and early reptiles (131).

The extant partition of the composite phylogeny obtain from timetree.org was compared and vetted using a secondary source of topological relationships based on phylotranscriptomic analysis (132).

Trait Evolution Analyses Details

The R script for trait evolution analyses is included in Appendix S3.

Relative Evolutionary Rates of Genes in Mandible Development: Additional Details

We first assessed the extent to which the candidate genes to be used in relative evolutionary rate analyses display high degrees of covariation with each other. We utilized a phenetic approach to build individual gene trees from FASTA sequences. The sequences were aligned in ClustalX and then converted to dendrograms in R using neighbor-joining and UPGMA methods. All possible pairwise comparisons of the 19 jaw development candidate genes are then conducted by pruning each pair of gene trees to the same ingroup and outgroup taxa. The trees then are compared using the entanglement algorithm implemented in the *dendextend* R package (*60*). One example script is included in Appendix S8.

We used the algorithms implemented in the *RERconverge* R package for relative evolutionary rate analysis (*61*). We vetted candidate mandible-development genes by including only those that have sequences available for more than 100 taxa. We generated a time tree using timetree.org for the vetted taxa, downloaded the sequences for all taxa in the list with candidate genes available, and estimated individual gene-based tree branch lengths using the *estimatePhangornTree()* function. The main time tree and individual gene-calibrated branch length files were then used to calculate relative evolutionary rates for each candidate gene. Next, three continuous trait variables from the functional morphological analyses (deflection/strain energy, MA, and AR) for the matching taxa in the morphometric dataset were imported and correlated evolutionary rates between the jaw development genes and each of the continuous trait variables were calculated. Adjusted *p* values were generated using the *correlateWithContinuuousPhenotype()* function to identify significant gene-morphology correlations. The R script associated with these analyses are included in Appendix S6.

The analyses summarized above and the figures generated were done using the R script included in Appendix S6 and S7.

*DLX5, DLX6, DLX3 & DLX7*—All gnathostomes are thought to show next expression of the *DLX* gene family, with an inferred in the number of *DLX* genes increasing as jaws evolved (133). The *DLX* gene family shares similarities in regulatory elements and expression patterns and interestingly, *DLX5* and *DLX6* loss results in a homeotic transformation of the mandible (lower jaw) into a maxillary (upper) jaw (134). During normal development, the distal region of the first pharyngeal (mandibular) arch is marked by expression of not only *DLX* genes, but also *ALX*4, *dHAND*, *BMP7*, and *PITX1*. On the other end, the proximal region correlates with elevated expression of *WNT5a*, *MEIS2* and *PRX2*.

Mice express six *DLX* genes, which are organized as three linked pairs in the genome (*DLX* 1/2, *DLX* 3/4 and *DLX* 5/6). *DLX* 1/2 are expressed in both the maxillary region of branchial arch 1 (mxBA1) and the mandibular region of branchial arch 1 (mdBA1), whereas *DLX* 5/6 are expressed in mdBA1 only (135). *DLX* 3/4 expression is restricted to a narrow domain within mdBA1. The mouse model suggests that *DLX* 1 & *DLX* 2 mostly regulate upper jaw development (although they are also expressed in the lower jaw), whereas *DLX* 5 and *DLX* 6 confer the lower jaw fate in mammals.

*GBX2*—*GBX2* is a direct downstream target of the *DLX* 5/6 genes in the first branchial arch.

*PRX1, PRX2*—Through knock-out studies, it has been shown that *PRX2* loss of-function mutants do not show any abnormalities, but *PRX1/PRX2* double mutants show severe skeletal abnormalities in the skull, craniofacial region, and limbs (136) . A striking defects in the lower jaw includes hypoplasticity and presence of just incisors. Results from Berge et al. (136) demonstrate that they are required for correct outgrowth of the lower jaw. *PRX1* loss-of-function mutants have skeletal abnormalities that are most severe at the lateral aspect of the skull, but milder defects are found in jaws, axial skeleton and limbs (137).

*MEIS2*—*MEIS2* has been shown to be involved in the normal development of the upper and lower jaw, in addition to a variety of other structures (137).

<u>Gene sequence sampling approach</u>

*BLAST*—The search was carried out in the blastn program (138) with the following configurations: In the “Choose Search Set” section the database chosen was “Nucleotide collection (nr/nt)” and the search was constrained to vertebrates using the Taxid: 7742 in the “Organism” field. In the Program Selection section, the search was optimized by using “More dissimilar sequences (discontiguous megablast)” which is the algorithm intended for cross-species comparisons. In the “Algorithm” parameters section, the “Max target sequences” chosen was 5000, the default values of the other parameters were kept.

The search for similar sequences to the query sequence was made in the same database for all genes discussed in the previous section, namely: "Nucleotide collection (nt)" which is described from the BLAST results page as:
"[nt] consists of GenBank+EMBL+DDBJ+PDB+RefSeq sequences, but excludes EST, STS, GSS, WGS, TSA, patent sequences as well as phase 0, 1, and 2 HTGS sequences and sequences longer than 100Mb. The database is non-redundant. Identical sequences have been merged into one entry, while preserving the accession, GI, title and taxonomy information for each entry".

The RIDs (alphanumeric codes that identify search results in Blast web page for retrieval within 24 hours of query creation), program, database, query ID, description, molecule type, query length was recorded for each gene.

Next, from the results page the Description Table was downloaded as a CSV file, the name of the file includes for default the RID. Also, the name of the gene and the number of sequences targeted (5000) was added to file names. Even though the max target sequences were 5,000, the BLAST search did not always return 5,000 sequences (see Table S3, Data S16) for each gene. Furthermore, in all cases due to the algorithm chosen for the search, sequences with a gene name different from the name of the query sequence were recovered. Also, some records were not properly filled out as they do not feature the scientific name in the relevant data column; instead, a scientific name was pasted in the description of the gene. Therefore, we carried out the following vetting procedure to further standardize the verify the gene sequences downloaded and potentially incorporated into downstream analyses:

1. Each CSV file was imported into Excel as a table.
2. A first filter was applied by extracting the exact name of the gene in the description tab into a new sheet in the same file.
3. Then, a filter was applied to exclude instances where a sequence was labeled "[gene]-like" rather than as the gene in question. *DLX*1, *DLX* 2, *DLX* 3, *DLX* 4, *DLX* 5, *DLX* 6, *MEIS*2, *UNC*5c all included sequences with such designations. These sequences were removed from further consideration.
4. A second filter was applied in the original sheet to look for sequences with synonyms names genes, using for this purpose the information obtained from GenBank. The sequences resulting were copied and pasted into the previously created sheet.
5. The table was reduced so that only columns representing gene description, scientific name, E-value, I (accession length), and accession number were retained.
6. The queried sequences were sorted by scientific name and by sequence length.
7. Once the sequences were ordered by scientific name and accession length, the list was searched for sequences whose scientific name was missing. In some cases, the scientific name was included in the description tab instead. In order to include the maximum number of correctly-identified sequences possible, the scientific names for those missing them in the name field were extracted when available (Table S5).
8. We retained a single sequence per taxon per gene queried by retaining the query result with the longest sequence length. In the case of a sequence with presumed taxonomic identity, the presumed sequence was kept only when there was no other sequence properly filled out and it was the longest among the presumed sequences.
9. Another round of vetting was conducted to ensure that only gnathostome taxa were included in the table.

10. The vetted gene sequence query tables were saved (Data S17) and used to extract the FASTA files from GenBank.

References in Supplementary Materials


1. P. R. Bell, E. Snively, L. Shychoski, A Comparison of the Jaw Mechanics in Hadrosaurid and Ceratopsid Dinosaurs Using Finite Element Analysis. *Anat. Rec.* **292**, 1338–1351 (2009).
2. M. J. Benton, J. L. Allen, Boreopricea from the Lower Triassic of Russia, and the relationships of the prolacertiform reptiles. *Palaeontology* **40**, 931–953 (1997).
3. P. Bover, J. A. Alcover, J. J. Michaux, L. Hautier, R. Hutterer, Body Shape and Life Style of the Extinct Balearic Dormouse Hypnomys (Rodentia, Gliridae): New Evidence from the Study of Associated Skeletons. *PLoS One* **5**, e15817 (2011).
4. D. W. Dilkes, The early Triassic rhynchosaur Mesosuchus browni and the interrelationships of basal archosauromorph reptiles. *Philos. Trans. R. Soc. London. Ser. B Biol. Sci.* **353**, 501–541 (1998).
5. H. Dutel, M. Herbin, G. Clément, A. Herrel, Bite Force in the Extant Coelacanth Latimeria: The Role of the Intracranial Joint and the Basicranial Muscle. *Curr. Biol.* **25**, 1228–1233 (2015).
6. S. E. Evans, *The Skull of lizards and tuatara* (2008).
7. M. D. Ezcurra, R. J. Butler, Taxonomy of the proterosuchid archosauriforms (Diapsida: Archosauromorpha) from the earliest Triassic of South Africa, and implications for the early archosauriform radiation. *Palaeontology* **58**, 141–170 (2015).
8. D. J. Field, R. Campbell-Malone, J. A. Goldbogen, R. E. Shadwick, Quantitative Computed Tomography of Humpback Whale (Megaptera novaeangliae) Mandibles: Mechanical Implications for Rorqual Lunge-Feeding. *Anat. Rec.* **293**, 1240–1247 (2010).
9. T. M. Fletcher, C. M. Janis, E. J. Rayfield, Finite element analysis of ungulate jaws: can mode of digestive physiology be determined. *Palaeontol. Electron.* **13**, 1–15 (2010).
10. J. Flynn, S. J. Nesbitt, M. J. Parrish, L. Ranivoharimanana, A. R. Wyss, A new species of Azendohsaurus (Diapsida: Archosauromorpha) from the Triassic Isalo Group of southwestern Madagascar: cranium and mandible. *Palaeontology* **53**, 669–688 (2010).
11. C. Gans, R. Montero, “An Atlas of Amphisbaenian Skull Anatomy” in *Biology of the Reptilia, Vol. 21, Morphology I, The Skull and Appendicular Locomotor Apparatus of Lepidosauria*, C. Gans, A. Gaunt, K. Adler, Eds. (Society for the Study of Amphibians and Reptiles, 2008), pp. 621–738.
12. D. J. Gower, The cranial and mandibular osteology of a new rauisuchian archosaur from the Middle Triassic of southern Germany . *Stuttgarter Beiträge zur Naturkunde. Ser. B (Geologie und Paläontologie)* **280**, 1–49 (1999).
13. A. B. Heckert, S. G. Lucas, J. A. Spielmann, A new species of the enigmatic archosauromorph Doswellia from the Upper Triassic Bluewater Creek Formation, New Mexico, USA. *Palaeontology* **55**, 1333–1348 (2012).
14. A. Herrel, *et al.*, “Feeding in Amphibians: Evolutionary Transformations and Phenotypic Diversity as Drivers of Feeding System Diversity” in *Feeding in Vertebrates: Evolution, Morphology, Behavior, Biomechanics*, V. Bels, I. Q. Whishaw, Eds. (Springer International Publishing, 2019), pp. 431–467.
15. J. J. Hill, M. N. Puttick, T. L. Stubbs, E. J. Rayfield, P. C. J. Donoghue, Evolution of jaw disparity in fishes. *Palaeontology* **61**, 847–854 (2018).
16. J. M. Parrish, Phylogeny of the Erythrosuchidae (Reptilia: Archosauriformes). *J. Vertebr. Paleontol.* **12**, 93–102 (1992).

17. C. F. Kammerer, L. Grande, M. W. Westneat, Comparative and developmental functional morphology of the jaws of living and fossil gars (Actinopterygii: Lepisosteidae). *J. Morphol.* **267**, 1017–1031 (2006).
18. T. Kleinteich, A. Haas, A. P. Summers, Caecilian jaw-closing mechanics: integrating two muscle systems. *J. R. Soc. Interface* **5**, 1491–1504 (2008).
19. T. M. Lehman, S. L. Tomlinson, Terlinguachelys fischbecki, a new genus and species of sea turtle (Chelonioidea: Protostegidae) from the Upper Cretaceous of Texas. *J. Paleontol.* **78**, 1163–1178 (2004).
20. Z.-X. Luo, J. A. Schultz, E. G. Ekdale, "Evolution of the Middle and Inner Ears of Mammaliaforms: The Approach to Mammals" in *Evolution of the Vertebrate Ear : Evidence from the Fossil Record*, J. A. Clack, R. R. Fay, A. N. Popper, Eds. (Springer International Publishing, 2016), pp. 139–174.
21. J. Marcé-Nogué, T. A. Püschel, T. M. Kaiser, A biomechanical approach to understand the ecomorphological relationship between primate mandibles and diet. *Sci. Rep.* **7**, 8364 (2017).
22. M. Thomas, L. Zhe-Xi, Homoplasy in the Mammalian Ear. *Science (80-. ).* **307**, 861–862 (2005).
23. M. A. Kolmann, S. B. Crofts, M. N. Dean, A. P. Summers, N. R. Lovejoy, Morphology does not predict performance: jaw curvature and prey crushing in durophagous stingrays. *J. Exp. Biol.* **218**, 3941–3949 (2015).
24. F. Miedema, S. N. F. Spiekman, V. Fernandez, J. W. F. Reumer, T. M. Scheyer, Cranial morphology of the tanystropheid Macrocnemus bassanii unveiled using synchrotron microtomography. *Sci. Rep.* **10**, 12412 (2020).
25. B. A. Mikuriya, The gross anatomy and microscopic anatomy of the tongue and lower jaw of Gnathonemus petersii (Gthr. 1862) (Mormyridae, Teleostei). *Zeitschrift für Morphol. der Tiere* **73**, 195–208 (1972).
26. N. M. Morales-García, T. D. Burgess, J. J. Hill, P. G. Gill, E. J. Rayfield, The use of extruded finite-element models as a novel alternative to tomography-based models: a case study using early mammal jaws. *J. R. Soc. Interface* **16**, 20190674 (2019).
27. R. Motani, *et al.*, Lunge feeding in early marine reptiles and fast evolution of marine tetrapod feeding guilds. *Sci. Rep.* **5**, 8900 (2015).
28. C. A. Navarro, E. Martin-Silverstone, T. L. Stubbs, Morphometric assessment of pterosaur jaw disparity. *R. Soc. Open Sci.* **5**, 172130 (2022).
29. J. M. Neenan, M. Ruta, J. A. Clack, E. J. Rayfield, Feeding biomechanics in Acanthostega and across the fish–tetrapod transition. *Proc. R. Soc. B Biol. Sci.* **281**, 20132689 (2014).
30. G. Parra-Olea, *et al.*, Biology of tiny animals: three new species of minute salamanders (Plethodontidae: Thorius) from Oaxaca, Mexico. *PeerJ* **4**, e2694 (2016).
31. R. E. Blanco, W. W. Jones, G. A. Grinspan, Fossil marsupial predators of South America (Marsupialia, Borhyaenoidea): bite mechanics and palaeobiological implications. *Alcheringa An Australas. J. Palaeontol.* **35**, 377–387 (2011).
32. D. A. Reed, J. Iriarte-Diaz, T. G. H. Diekwisch, A three dimensional free body analysis describing variation in the musculoskeletal configuration of the cynodont lower jaw. *Evol. Dev.* **18**, 41–53 (2016).
33. R. R. Reisz, D. S. Berman, D. Scott, The cranial anatomy and relationships of Secodontosaurus, an unusual mammal-like reptile (Synapsida: Sphenacodontidae) from the early Permian of Texas. *Zool. J. Linn. Soc.* **104**, 127–184 (1992).

34. R. R. Reisz, Origin of dental occlusion in tetrapods: signal for terrestrial vertebrate evolution? *J. Exp. Zool. Part B Mol. Dev. Evol.* **306B**, 261–277 (2006).
35. A. S. Romer, The Chanares (Argentina) Triassic reptile fauna. XI. Two new long-snouted thecodonts, Chanaresuchus and Gualosuchus. *Breviora.* **379**, 1–22 (1971).
36. S. H. Reynolds, *The vertebrate skeleton* (University press, 1897).
37. J. Schaeffer, M. J. Benton, E. J. Rayfield, T. L. Stubbs, Morphological disparity in theropod jaws: comparing discrete characters and geometric morphometrics. *Palaeontology* **63**, 283–299 (2020).
38. S. Serrano-Fochs, S. De Esteban-Trivigno, J. Marcé-Nogué, J. Fortuny, R. A. Fariña, Finite Element Analysis of the Cingulata Jaw: An Ecomorphological Approach to Armadillo's Diets. *PLoS One* **10**, e0120653 (2015).
39. R. B. Sookias, *et al.*, The craniomandibular anatomy of the early archosauriform Euparkeria capensis and the dawn of the archosaur skull. *R. Soc. Open Sci.* **7**, 200116 (2022).
40. Z. J. Tseng, D. F. Su, X. Wang, S. C. White, X. Ji, Feeding capability in the extinct giant Siamogale melilutra and comparative mandibular biomechanics of living Lutrinae. *Sci. Rep.* **7**, 15225 (2017).
41. C. W. Walmsley, *et al.*, Why the Long Face? The Mechanics of Mandibular Symphysis Proportions in Crocodiles. *PLoS One* **8**, e53873 (2013).
42. S. P. Welles, Dilophosaurus (Reptilia, Saurischia), a new name for a dinosaur. *J. Paleontol.* **44**, 989 (1970).
43. C. D. Wilga, P. J. Motta, Feeding mechanism of the atlantic guitarfish rhinobatos lentiginosus:modulation of kinematic and motor activity. *J. Exp. Biol.* **201**, 3167–3183 (1998).
44. L. M. Witmer, K. D. Rose, Biomechanics of the jaw apparatus of the gigantic Eocene bird Diatryma: implications for diet and mode of life. *Paleobiology* **17**, 95–120 (1991).
45. C.-F. Zhou, B.-A. Bhullar, A. Neander, T. Martin, Z.-X. Luo, New Jurassic mammaliaform sheds light on early evolution of mammal-like hyoid bones. *Science (80-. ).* **365**, 276–279 (2019).
46. P. E. Ahlberg, J. A. Clack, The smallest known Devonian tetrapod shows unexpectedly derived features. *R. Soc. Open Sci.* **7**, 192117 (2022).
47. P. E. Ahlberg, Z. Johanson, Osteolepiforms and the ancestry of tetrapods. *Nature* **395**, 792–794 (1998).
48. A. H. Turner, P. J. Makovicky, M. A. Norell, A Review of Dromaeosaurid Systematics and Paravian Phylogeny. *Bull. Am. Museum Nat. Hist.*, 1–206 (2012).
49. B. Andres, J. Clark, X. Xu, The Earliest Pterodactyloid and the Origin of the Group. *Curr. Biol.* **24**, 1011–1016 (2014).
50. J. Anquetin, C. Püntener, J.-P. Billon-Bruyat, Portlandemys gracilis n. sp., a New Coastal Marine Turtle from the Late Jurassic of Porrentruy (Switzerland) and a Reconsideration of Plesiochelyid Cranial Anatomy. *PLoS One* **10**, e0129193 (2015).
51. M. Archer, *et al.*, Australia's first fossil marsupial mole (Notoryctemorphia) resolves controversies about their evolution and palaeoenvironmental origins. *Proc. Biol. Sci.* **278**, 1498–1506 (2011).
52. M. S. Arkhangelsky, N. G. Zverkov, O. S. Spasskaya, A. V Evgrafov, On the First Reliable Record of the Ichthyosaur Ophthalmosaurus icenicus Seeley in the Oxfordian–Kimmeridgian Beds of European Russia. *Paleontol. J.* **52**, 49–57 (2018).

53. S. Bi, Y. Wang, J. Guan, X. Sheng, J. Meng, Three new Jurassic euharamiyidan species reinforce early divergence of mammals. *Nature* **514**, 579–584 (2014).
54. K. S. Brink, R. R. Reisz, Hidden dental diversity in the oldest terrestrial apex predator Dimetrodon. *Nat. Commun.* **5**, 3269 (2014).
55. B. P. Polley, R. R. Reisz, A new Lower Permian trematopid (Temnospondyli: Dissorophoidea) from Richards Spur, Oklahoma. *Zool. J. Linn. Soc.* **161**, 789–815 (2011).
56. S. L. Brusatte, T. D. Carr, The phylogeny and evolutionary history of tyrannosauroid dinosaurs. *Sci. Rep.* **6**, 20252 (2016).
57. K. S. W. Campbell, R. E. Barwick, Paleozoic Dipnoan Phylogeny: Functional Complexes and Evolution Without Parsimony. *Paleobiology* **16**, 143–169 (1990).
58. C. J. Burrow, K. Trinajstic, J. Long, First acanthodian from the Upper Devonian (Frasnian) Gogo Formation, Western Australia. *Hist. Biol.* **24**, 349–357 (2012).
59. T. D. Carr, Craniofacial ontogeny in Tyrannosauridae (Dinosauria, Coelurosauria). *J. Vertebr. Paleontol.* **19**, 497–520 (1999).
60. M. Zhang, X. Yu, Reexamination of the relationship of Middle Devonian osteolepids: fossil characters and their interpretations. *Am. Museum Novit.* **3189**, 1–20 (1997).
61. J. M. Clark, J. A. Hopson, Distinctive mammal-like reptile from Mexico and its bearing on the phylogeny of the Tritylodontidae. *Nature* **315**, 398–400 (1985).
62. A. M. Clement, A new species of long-snouted lungfish from the Late Devonian of Australia, and its functional and biogeographical implications. *Palaeontology* **55**, 51–71 (2012).
63. J. O. Farlow, Scipionyx samniticus (Theropoda: Compsognathidae) from the Lower Cretaceous of Italy: osteology, ontogenetic assessment, phylogeny, soft tissue anatomy, taphonomy and palaeobiology (2012).
64. K. E. Criswell, The comparative osteology and phylogenetic relationships of African and South American lungfishes (Sarcopterygii: Dipnoi). *Zool. J. Linn. Soc.* **174**, 801–858 (2015).
65. F. Vecchia, Comments on Triassic pterosaurs with a commentary on the" ontogenetic stages" of Kellner (2015) and the validity of Bergamodactylus wildi. *Riv. Ital. di Paleontol. e Stratigr.* **124** (2018).
66. F. Vecchia, Seazzadactylus venieri gen. et sp. nov., a new pterosaur (Diapsida: Pterosauria) from the Upper Triassic (Norian) of northeastern Italy. *PeerJ* **7**, e7363 (2019).
67. H. Dutel, *et al.*, The Giant Cretaceous Coelacanth (Actinistia, Sarcopterygii) Megalocoelacanthus dobiei Schwimmer, Stewart & Williams, 1994, and Its Bearing on Latimerioidei Interrelationships. *PLoS One* **7**, e49911 (2012).
68. C. R. Eastman, Structure and relations of Mylostoma. *Bull. Museum Comp. Zool.* **50**, 1–29 (1906).
69. E. J. Swaby, D. R. Lomax, A revision of Temnodontosaurus crassimanus (Reptilia: Ichthyosauria) from the Lower Jurassic (Toarcian) of Whitby, Yorkshire, UK. *Hist. Biol.* **33**, 2715–2731 (2021).
70. F. Vecchia, Triassic pterosaurs. *Anatomy, Phylogeny Palaeobiology Early Archosaurs their Kin* **379**, 0 (2013).
71. F. A. Pretto, M. C. Langer, C. L. Schultz, A new dinosaur (Saurischia: Sauropodomorpha) from the Late Triassic of Brazil provides insights on the evolution of sauropodomorph body plan. *Zool. J. Linn. Soc.* **185**, 388–416 (2019).

72. F. Witzmann, R. R. Schoch, A megalichthyid sarcopterygian fish from the Lower Permian (Autunian) of the Saar-Nahe Basin, Germany. *Geobios* **45**, 241–248 (2012).
73. G. E. Hooks, Systematic revision of the Protostegidae, with a redescription of Calcarichelys gemma Zangerl, 1953. *J. Vertebr. Paleontol.* **18**, 85–98 (1998).
74. P. Godefroit, *et al.*, Reduced plumage and flight ability of a new Jurassic paravian theropod from China. *Nat. Commun.* **4**, 1394 (2013).
75. T. Holland, J. A. Long, On the phylogenetic position of Gogonasus andrewsae Long 1985, within the Tetrapodomorpha. *Acta Zool.* **90**, 285–296 (2009).
76. C. M. Holliday, N. M. Gardner, A New Eusuchian Crocodyliform with Novel Cranial Integument and Its Significance for the Origin and Evolution of Crocodylia. *PLoS One* **7**, e30471 (2012).
77. J. Huang, *et al.*, Repeated evolution of durophagy during ichthyosaur radiation after mass extinction indicated by hidden dentition. *Sci. Rep.* **10**, 7798 (2020).
78. J. Botha, F. Abdala, R. Smith, The oldest cynodont: new clues on the origin and early diversification of the Cynodontia. *Zool. J. Linn. Soc.* **149**, 477–492 (2007).
79. M. L. Jacobs, D. M. Martill, N. Ibrahim, N. Longrich, A new species of Coloborhynchus (Pterosauria, Ornithocheiridae) from the mid-Cretaceous of North Africa. *Cretac. Res.* **95**, 77–88 (2019).
80. J. Boyle, M. J. Ryan, New information on Titanichthys (Placodermi, Arthrodira) from the Cleveland Shale (Upper Devonian) of Ohio, USA. *J. Paleontol.* **91**, 318–336 (2017).
81. J. K. O'Connor, *et al.*, Phylogenetic support for a specialized clade of Cretaceous enantiornithine birds with information from a new species. *J. Vertebr. Paleontol.* **29**, 188–204 (2009).
82. J. Klembara, J. A. Clack, A. R. Milner, M. Ruta, Cranial anatomy, ontogeny, and relationships of the Late Carboniferous tetrapod Gephyrostegus bohemicus Jaekel, 1902. *J. Vertebr. Paleontol.* **34**, 774–792 (2014).
83. J. Lu, H. Pu, L. Xu, Y. Wu, X. Wei, Largest Toothed Pterosaur Skull from the Early Cretaceous Yixian Formation of Western Liaoning, China, with Comments on the Family Boreopteridae. *Acta Geol. Sin.* **86**, 287–293 (2012).
84. A. W. A. Kellner, L. C. Weinschütz, B. Holgado, R. A. M. Bantim, J. M. Sayao, A new toothless pterosaur (Pterodactyloidea) from Southern Brazil with insights into the paleoecology of a Cretaceous desert. *An. Acad. Bras. Cienc.* **91** (2019).
85. K. Chiba, M. J. Ryan, F. Fanti, M. A. Loewen, D. C. Evans, New material and systematic re-evaluation of Medusaceratops lokii (Dinosauria, Ceratopsidae) from the Judith River Formation (Campanian, Montana). *J. Paleontol.* **92**, 272–288 (2018).
86. A. Kriloff, *et al.*, Evolution of bone microanatomy of the tetrapod tibia and its use in palaeobiological inference. *J. Evol. Biol.* **21**, 807–826 (2008).
87. M. C. Lamanna, H.-D. Sues, E. R. Schachner, T. R. Lyson, A New Large-Bodied Oviraptorosaurian Theropod Dinosaur from the Latest Cretaceous of Western North America. *PLoS One* **9**, e92022 (2014).
88. L. Li, J. Wang, X. Zhang, S. Hou, A new enantiornithine bird from the lower cretaceous Jiufotang formation in Jinzhou area, Western Liaoning Province, China. *Acta Geol. Sin. Ed.* **86**, 1039–1044 (2012).
89. M. A. Loewen, R. B. Irmis, J. J. W. Sertich, P. J. Currie, S. D. Sampson, Tyrant Dinosaur Evolution Tracks the Rise and Fall of Late Cretaceous Oceans. *PLoS One* **8**, e79420 (2013).

90. J. Lü, *et al.*, A New Oviraptorid Dinosaur (Dinosauria: Oviraptorosauria) from the Late Cretaceous of Southern China and Its Paleobiogeographical Implications. *Sci. Rep.* **5**, 11490 (2015).
91. J. Lü, S. L. Brusatte, A large, short-armed, winged dromaeosaurid (Dinosauria: Theropoda) from the Early Cretaceous of China and its implications for feather evolution. *Sci. Rep.* **5**, 11775 (2015).
92. Z.-X. Luo, *et al.*, New evidence for mammaliaform ear evolution and feeding adaptation in a Jurassic ecosystem. *Nature* **548**, 326–329 (2017).
93. Z.-X. Luo, S. Gatesy, N. Shubin, Mandibular and dental characteristics of Late Triassic mammaliaform Haramiyavia and their ramifications for basal mammal evolution. *Proc. Natl. Acad. Sci.* **112**, E7101–E7109 (2015).
94. M. D. Ercoli, F. J. Prevosti, Estimación de masa de las especies de Sparassodonta (Mammalia, Metatheria) de edad Santacrucense (Mioceno temprano) a partir del tamaño del centroide de los elementos apendiculares: inferencias paleoecológicas. *Ameghiniana* **48**, 462–479 (2011).
95. J. E. Martin, K. Lauprasert, H. Tong, V. Suteethorn, E. Buffetaut, An Eocene tomistomine from peninsular Thailand. *Ann. Paléontologie* **105**, 245–253 (2019).
96. A. G. Martinelli, M. B. Soares, T. V. De Oliveira, P. G. Rodrigues, C. L. Schultz, The Triassic eucynodont Candelariodon barberenai revisited and the early diversity of stem prozostrodontians. *Acta Palaeontol. Pol.* **62**, 527–542 (2017).
97. M. Danowitz, S. Hou, M. Mihlbachler, V. Hastings, N. Solounias, A combined-mesowear analysis of late Miocene giraffids from North Chinese and Greek localities of the Pikermian Biome. *Palaeogeogr. Palaeoclimatol. Palaeoecol.* **449**, 194–204 (2016).
98. H. Pu, *et al.*, An Unusual Basal Therizinosaur Dinosaur with an Ornithischian Dental Arrangement from Northeastern China. *PLoS One* **8**, e63423 (2013).
99. T. Qiao, B. King, J. A. Long, P. E. Ahlberg, M. Zhu, Early Gnathostome Phylogeny Revisited: Multiple Method Consensus. *PLoS One* **11**, e0163157 (2016).
100. R. Pêgas, B. Holgado, M. Leal, On Targaryendraco wiedenrothi gen. nov. (Pterodactyloidea, Pteranodontoidea, Lanceodontia) and recognition of a new cosmopolitan lineage of Cretaceous toothed pterodactyloids. *Hist. Biol.* **33**, 1266–1280 (2021).
101. M. Rucklin, First selenosteid placoderms from the eastern Anti-Atlas of Morocco; osteology, phylogeny and palaeogeographical implications. *Palaeontology* **54**, 25–62 (2011).
102. R. Takasaki, K. Chiba, Y. Kobayashi, P. J. Currie, A. R. Fiorillo, Reanalysis of the phylogenetic status of Nipponosaurus sachalinensis (Ornithopoda: Dinosauria) from the Late Cretaceous of Southern Sakhalin. *Hist. Biol.* **30**, 694–711 (2018).
103. H.-P. Schultze, Scales, Enamel, Cosmine, Ganoine, and Early Osteichthyans. *Comptes Rendus Palevol* **15**, 83–102 (2016).
104. S.-X. Jiang, X.-L. Wang, X. Meng, X. Cheng, A new boreopterid pterosaur from the Lower Cretaceous of western Liaoning, China, with a reassessment of the phylogenetic relationships of the Boreopteridae. *J. Paleontol.* **88**, 823–828 (2014).
105. S. H. Hwang, M. A. Norell, J. Qiang, G. Keqin, A large Compsognathid from the early cretaceous Yixian formation of China. *J. Syst. Palaeontol.* **2**, 13–30 (2004).
106. B. Swartz, A Marine Stem-Tetrapod from the Devonian of Western North America. *PLoS One* **7**, e33683 (2012).

107. T. S. Myers, First North American occurrence of the toothed pteranodontoid pterosaur Cimoliopterus. *J. Vertebr. Paleontol.* **35**, e1014904 (2015).
108. V. M. Arbour, P. J. Currie, Systematics, phylogeny and palaeobiogeography of the ankylosaurid dinosaurs. *J. Syst. Palaeontol.* **14**, 385–444 (2016).
109. X. Wang, *et al.*, New evidence from China for the nature of the pterosaur evolutionary transition. *Sci. Rep.* **7**, 42763 (2017).
110. X. Wang, A. W. A. Kellner, Z. Zhou, D. de A. Campos, Pterosaur diversity and faunal turnover in Cretaceous terrestrial ecosystems in China. *Nature* **437**, 875–879 (2005).
111. X. Cheng, X. Wang, S. Jiang, A. W. A. Kellner, A new scaphognathid pterosaur from western Liaoning, China. *Hist. Biol.* **24**, 101–111 (2012).
112. X. Xu, *et al.*, A new dromaeosaurid (Dinosauria: Theropoda) from the Upper Cretaceous Wulansuhai formation of inner Mongolia, China. *Zootaxa* **2403**, 1–9 (2010).
113. L. Xu, *et al.*, A new ornithomimid dinosaur with North American affinities from the Late Cretaceous Qiupa Formation in Henan Province of China. *Cretac. Res.* **32**, 213–222 (2011).
114. X. Xu, Q. Tan, J. Wang, X. Zhao, L. Tan, A gigantic bird-like dinosaur from the Late Cretaceous of China. *Nature* **447**, 844–847 (2007).
115. Y.-A. Zhu, M. Zhu, A redescription of Kiangyousteus yohii (Arthrodira: Eubrachythoraci) from the Middle Devonian of China, with remarks on the systematics of the Eubrachythoraci. *Zool. J. Linn. Soc.* **169**, 798–819 (2013).
116. G. C. Young, J. A. Long, Crossopterygian fishes from the Devonian of Antarctica: systematics, relationships and biogeographic significance. *Rec. Aust. Museum, Suppl.* **14**, 1–77 (1992).
117. X. Xu, *et al.*, A bizarre Jurassic maniraptoran theropod with preserved evidence of membranous wings. *Nature* **521**, 70–73 (2015).
118. J. A. Long, New ischnacanthid acanthodians from the Early Devonian of Australia, with comments on acanthodian interrelationships. *Zool. J. Linn. Soc.* **87**, 321–339 (1986).
119. Y.-A. Zhu, M. Zhu, J.-Q. Wang, Redescription of Yinostius major (Arthrodira: Heterostiidae) from the Lower Devonian of China, and the interrelationships of Brachythoraci. *Zool. J. Linn. Soc.* **176**, 806–834 (2016).
120. F. A. Bockmann, M. R. De Carvalho, M. De Carvalho, The salmon, the lungfish (or the coelacanth) and the cow: a revival? *Zootaxa* **3750**, 265–276 (2013).
121. D. Marjanović, M. Laurin, Phylogeny of Paleozoic limbed vertebrates reassessed through revision and expansion of the largest published relevant data matrix. *PeerJ* **6**, e5565 (2019).
122. D. Casane, P. Laurenti, Why coelacanths are not 'living fossils.' *BioEssays* **35**, 332–338 (2013).
123. J.-P. M. Hodnett, D. K. Elliott, T. J. Olson, J. H. Wittke, Ctenacanthiform sharks from the Permian Kaibab Formation, northern Arizona. *Hist. Biol.* **24**, 381–395 (2012).
124. C. Hendrickx, S. A. Hartman, O. Mateus, An Overview of Non-Avian Theropod Discoveries and Classification. *PalArch's J. Vertebr. Palaeontol.* **12**, 1–73 (2015).
125. C. Hendrickx, O. Mateus, E. Buffetaut, Morphofunctional Analysis of the Quadrate of Spinosauridae (Dinosauria: Theropoda) and the Presence of Spinosaurus and a Second Spinosaurine Taxon in the Cenomanian of North Africa. *PLoS One* **11**, e0144695 (2016).
126. J. Lu, S. Giles, M. Friedman, J. L. den Blaauwen, M. Zhu, The Oldest Actinopterygian Highlights the Cryptic Early History of the Hyperdiverse Ray-Finned Fishes. *Curr. Biol.*

**26**, 1602–1608 (2016).
127. G. S. Bever, T. R. Lyson, D. J. Field, B.-A. S. Bhullar, Evolutionary origin of the turtle skull. *Nature* **525**, 239–242 (2015).
128. P. Thorne, R. Marcello, M. Benton, Resetting the evolution of marine reptiles at the Triassic-Jurassic boundary. *Proc. Natl. Acad. Sci.* **108**, 8339–8344 (2011).
129. J. Huang, *et al.*, The new ichthyosauriform Chaohusaurus brevifemoralis (Reptilia, Ichthyosauromorpha) from Majiashan, Chaohu, Anhui Province, China. *PeerJ* **7**, e7561 (2019).
130. M. Ruta, J. Botha-Brink, S. A. Mitchell, M. J. Benton, The radiation of cynodonts and the ground plan of mammalian morphological diversity. *Proc. R. Soc. B Biol. Sci.* **280**, 20131865 (2013).
131. D. P. Ford, R. B. J. Benson, The phylogeny of early amniotes and the affinities of Parareptilia and Varanopidae. *Nat. Ecol. Evol.* **4**, 57–65 (2020).
132. I. Irisarri, *et al.*, Phylotranscriptomic consolidation of the jawed vertebrate timetree. *Nat. Ecol. Evol.* **1**, 1370–1378 (2017).
133. A. Graham, Jaw Development: Chinless Wonders. *Curr. Biol.* **12**, R810–R812 (2002).
134. M. Depew, T. Lufkin, J. L. Rubenstein, Specification of Jaw Subdivisions by Dlx Genes. *Science (80-. ).* **298**, 381–385 (2002).
135. J. Jeong, *et al.*, Dlx genes pattern mammalian jaw primordium by regulating both lower jaw-specific and upper jaw-specific genetic programs. *Development* **135**, 2905–2916 (2008).
136. D. ten Berge, *et al.*, Prx1 and Prx2 are upstream regulators of sonic hedgehog and control cell proliferation during mandibular arch morphogenesis. *Development* **128**, 2929–2938 (2001).
137. O. Machon, J. Masek, O. Machonova, S. Krauss, Z. Kozmik, Meis2 is essential for cranial and cardiac neural crest development. *BMC Dev. Biol.* **15**, 40 (2015).
138. Z. Zhang, S. Schwartz, L. Wagner, W. Miller, A Greedy Algorithm for Aligning DNA Sequences. *J. Comput. Biol.* **7**, 203–214 (2000).
139. T. Square, *et al.*, A gene expression map of the larval Xenopus laevis head reveals developmental changes underlying the evolution of new skeletal elements. *Dev. Biol.* **397**, 293–304 (2015).

**Appendix S1. Functional morphological data interpolation and Wilcoxon tests**

```
#R script to take theoretical model simulation results from FEA and use double linear
interpolation formula to extract FEA values for taxon-specific xy values from PC1-PC2 plot

#libraries used
library(interp)
library(dplyr)
library(autoimage)

#import data
setwd(choose.dir())
theo_data <- read.csv(choose.files(),header=T)
taxon_data <- read.csv(choose.files(),header=T)

#map theoretical data and taxon data onto same grid framework
linMap <- function(x, from, to)
  (x - min(x)) / max(x - min(x)) * (to - from) + from
x <- linMap(taxon_data[,3], 0, 9)
y <- linMap(taxon_data[,4], 0, 9)
theox <- linMap(theo_data[,2], 0, 9)
theoy <- linMap(theo_data[,3], 0, 9)

#calculation of interpolated values in MA, AR
interp_MA <- interp(theox,theoy,theo_data[,8],
                    x,y,input="points",
                    output="points")
interp_AR <- interp(theox,theoy,theo_data[,9],
                    x,y,input="points",
                    output="points")
#(repeat for other biomechanical traits…)

#combining all interpolated data and write to file
taxon_interp_all <- cbind(taxon_data,interp_maxDisp$z,
                          interp_adjSE$z,
                          interp_maxDisp_imj$z,
                          interp_adjSE_imj$z,
                          interp_MA$z,
                          interp_AR$z)
write.csv(taxon_interp_all, "n1325_interp_5-18-2021.csv")

#bootstrap resample from within morphospace limits to generate null distribution, then calculate
Wilcoxon test (non-parametric equivalent of chi square test) against observed taxon values
## then summarize % of 1000 bootstraps with similar distributions as observed

#library for chi squared/Wilcoxon analyses and plotting
```

```
library(MASS)
library(ggplot2)
library(ggpubr)

#set up bootstrap resampling objects
bootstrap_sample <- matrix(nrow=nrow(taxon_interp_all),ncol=2)
bootstrap_stat_pc1 <- vector(length=1000)
bootstrap_stat_pc2 <- vector(length=1000)
bootstrap_stat_ma <- vector(length=1000)
bootstrap_stat_ar <- vector(length=1000)
pc1_counts <- taxon_interp_all$PC1
pc2_counts <- taxon_interp_all$PC2
ma_counts <- taxon_interp_all$interp_MA.z
ar_counts <- taxon_interp_all$interp_AR.z

pc1_boot_interp <- vector()
pc2_boot_interp <- vector()
ma_boot_interp <- vector()
ar_boot_interp <- vector()

#MA bootstrap
for (i in 1:1000) {
 bootstrap_sample[,1] <- runif(nrow(taxon_interp_all),min(x),max(x))
 bootstrap_sample[,2] <- runif(nrow(taxon_interp_all),min(y),max(y))
 interp_resample <- interp(theox,theoy,theo_data[,8],
                    bootstrap_sample[,1],bootstrap_sample[,2],input="points",
                              output="points")
 random_counts <- interp_resample$z
 ma_boot_interp <- append(ma_boot_interp,random_counts)
 wilcox_sample <- wilcox.test(ma_counts,random_counts )
 bootstrap_stat_ma[i] <- wilcox_sample$p.value
}

#AR bootstrap
for (i in 1:1000) {
 bootstrap_sample[,1] <- runif(nrow(taxon_interp_all),min(x),max(x))
 bootstrap_sample[,2] <- runif(nrow(taxon_interp_all),min(y),max(y))
 interp_resample <- interp(theox,theoy,theo_data[,9],
                    bootstrap_sample[,1],bootstrap_sample[,2],input="points",
                              output="points")
 random_counts <- interp_resample$z
 ar_boot_interp <- append(ar_boot_interp,random_counts)
 wilcox_sample <- wilcox.test(ar_counts,random_counts )
 bootstrap_stat_ar[i] <- wilcox_sample$p.value
}
```

```
#PC1 bootstrap
for (i in 1:1000) {
 bootstrap_sample[,1] <- runif(nrow(taxon_interp_all),min(x),max(x))
 bootstrap_sample[,2] <- runif(nrow(taxon_interp_all),min(y),max(y))
 interp_resample <- interp(theox,theoy,theo_data[,2],
                    bootstrap_sample[,1],bootstrap_sample[,2],input="points",
                                   output="points")
 random_counts <- interp_resample$z
 pc1_boot_interp <- append(pc1_boot_interp,random_counts)
 wilcox_sample <- wilcox.test(pc1_counts,random_counts )
 bootstrap_stat_pc1[i] <- wilcox_sample$p.value
}

#PC2 bootstrap
for (i in 1:1000) {
 bootstrap_sample[,1] <- runif(nrow(taxon_interp_all),min(x),max(x))
 bootstrap_sample[,2] <- runif(nrow(taxon_interp_all),min(y),max(y))
 interp_resample <- interp(theox,theoy,theo_data[,3],
                    bootstrap_sample[,1],bootstrap_sample[,2],input="points",
                                   output="points")
 random_counts <- interp_resample$z
 pc2_boot_interp <- append(pc2_boot_interp,random_counts)
 wilcox_sample <- wilcox.test(pc2_counts,random_counts )
 bootstrap_stat_pc2[i] <- wilcox_sample$p.value
}

#summarizing bootstrapped sample p values from 100 replicates
hist(bootstrap_stat_ma)
sum(bootstrap_stat_ma > 0.05)
hist(bootstrap_stat_ar)
sum(bootstrap_stat_ar > 0.05)
hist(bootstrap_stat_pc1)
sum(bootstrap_stat_pc1 > 0.05)
hist(bootstrap_stat_pc2)
sum(bootstrap_stat_pc2 > 0.05)

#plotting bootstrapped random sample versus observed sample - MA
observed <- data.frame(ma=ma_counts)
expected <- data.frame(ma=ma_boot_interp)
observed$cat <- 'observed'
expected$cat <- 'expected'
samp_histo_ma <- rbind(observed, expected)
#ggplot both null and observed distributions
ma_nonrandom <- ggplot(samp_histo_ma, aes(ma, fill = cat)) +
  geom_histogram(alpha = 0.5, aes(y = ..density..), position = 'identity', show.legend = FALSE)
  + labs(y= "Density", x = "Mechanical advantage")
```

```
#plotting bootstrapped random sample versus observed sample - AR
observed <- data.frame(ar=ar_counts)
expected <- data.frame(ar=ar_boot_interp)
observed$cat <- 'observed'
expected$cat <- 'expected'
samp_histo_ar <- rbind(observed, expected)
#ggplot both distributions
windows()
ar_nonrandom <- ggplot(samp_histo_ar, aes(ar, fill = cat)) +
  geom_histogram(alpha = 0.5, aes(y = ..density..), position = 'identity', show.legend = FALSE)
  +labs(y= "Density", x = "Aspect ratio")

#plotting bootstrapped random sample versus observed sample – PC1
observed <- data.frame(pc1=pc1_counts)
expected <- data.frame(pc1=pc1_boot_interp)
observed$cat <- 'observed'
expected$cat <- 'expected'
samp_histo_pc1 <- rbind(observed, expected)
#ggplot two distributions
windows()
pc1_nonrandom <- ggplot(samp_histo_pc1, aes(pc1, fill = cat)) +
  geom_histogram(alpha = 0.5, aes(y = ..density..), position = 'identity', show.legend = FALSE)
  +labs(y= "Density", x = "PC1 score")

#plotting bootstrapped random sample versus observed sample – PC2
observed <- data.frame(pc2=pc2_counts)
expected <- data.frame(pc2=pc2_boot_interp)
observed$cat <- 'observed'
expected$cat <- 'expected'
samp_histo_pc2 <- rbind(observed, expected)
#ggplot two distributions
windows()
pc2_nonrandom <- ggplot(samp_histo_pc2, aes(pc2, fill = cat)) +
  geom_histogram(alpha = 0.5, aes(y = ..density..), position = 'identity', show.legend = FALSE)
  +labs(y= "Density", x = "PC2 score")

#2x2 plot of results with panel labels
ggarrange(ma_nonrandom, ar_nonrandom, pc1_nonrandom, pc2_nonrandom,
        labels = c("A", "B", "C", "D"),
        ncol = 2, nrow = 2,
          common.legend = TRUE, legend = "bottom")
```

**Appendix S2. Maximum displacement and strain energy experimental validation**

```
#libraries for plotting
library(ggplot2)
library(ggpubr)
library(hrbrthemes)
library(phytools)
library(plotrix)

#read in dataset containing displacement and strain energy values from experimental analyses of
the 93 theoretical jaw models
validation_all_data <- read.csv("bending_test_validation_data_9-1-2021.csv",header=T)

#use linear model to model relationship between validation and simulation data
adjSE_validation <- lm(adj_SE ~ EXP_SE, data = validation_all_data)
summary(adjSE_validation)
plot(adjSE_validation)

#removing outliers (those at the edge of impossibility in morphospace) for adj_SE data
Q <- quantile(validation_all_data$adj_SE, probs=c(.25, .75), na.rm = FALSE)
iqr <- IQR(validation_all_data$adj_SE)
up <-  Q[2]+1.5*iqr # Upper Range
low<- Q[1]-1.5*iqr # Lower Range?
adjSE_no_outliers <- subset(validation_all_data, validation_all_data$adj_SE > (Q[1] - 1.5*iqr)
& validation_all_data$adj_SE < (Q[2]+1.5*iqr))
adjSE_validation_nooutliers <- lm(adj_SE ~ EXP_SE, data = adjSE_no_outliers)
summary(adjSE_validation_nooutliers)
plot(adjSE_no_outliers$adj_SE~adjSE_no_outliers$EXP_SE)
abline(adjSE_validation_nooutliers)

#linear model on displacement data
disp_validation <- lm(max_DY ~ EXP_MAX_DISP, data = validation_all_data)
summary(disp_validation)
windows()
plot(validation_all_data$max_DY~validation_all_data$EXP_MAX_DISP)
abline(disp_validation)

#linear model on displacement data using IMJ simulation results
disp_imj_validation <- lm(max_DY_IMJ ~ EXP_MAX_DISP, data = validation_all_data)
summary(disp_imj_validation)
windows()
plot(validation_all_data$max_DY_IMJ~validation_all_data$EXP_MAX_DISP)
abline(disp_imj_validation)

#linear model on strain energy data using IMJ simulation results
se_imj_validation <- lm(adj_SE_IMJ ~ EXP_SE, data = validation_all_data)
```

```
summary(se_imj_validation)
windows()
plot(validation_all_data$adj_SE_IMJ~validation_all_data$EXP_SE)
abline(se_imj_validation)

#removing outliers for adj_SE data from IMJ simulations
Qimj <- quantile(validation_all_data$adj_SE_IMJ, probs=c(.25, .75), na.rm = FALSE)
iqrimj <- IQR(validation_all_data$adj_SE_IMJ)
up <-  Qimj[2]+1.5*iqrimj # Upper Range
low<- Qimj[1]-1.5*iqrimj # Lower Range?
adjSEimj_no_outliers <- subset(validation_all_data, validation_all_data$adj_SE_IMJ > (Qimj[1]
- 1.5*iqrimj) & validation_all_data$adj_SE_IMJ < (Qimj[2]+1.5*iqrimj))
adjSEimj_validation_nooutliers <- lm(adj_SE_IMJ ~ EXP_SE, data = adjSEimj_no_outliers)
summary(adjSEimj_validation_nooutliers)
plot(adjSEimj_no_outliers$adj_SE_IMJ~adjSEimj_no_outliers$EXP_SE)
abline(adjSEimj_validation_nooutliers)

#conclusions from these validation regressions: FEA data are generally consistent with
experimental data in the center of the morphospace, but FEA data do not accurately portray the
values of jaw displacement and strain energy at the lower left edge of the morphospace.

#in conjunction with additional regression analyses conducted in excel, it is clear that DISP and
SE are linearly correlated when not at the lower left edge of the morphospace
#whereas disp was measured at a max of 10 mm, SE can't be easily/consistently calculated for
flexible jaw models. Therefore, displacement is treated as interchangeable with SE in
downstream analyses

#plot validate theomorph values as heatmap

# Give extreme colors to set gradient
#heatmap only
windows()
ggplot(validation_all_data, aes(PC1, PC2, fill= EXP_MAX_DISP)) +
  geom_tile() +
  scale_fill_gradient(low="white", high="blue") +
  theme_ipsum() + ggplot(n1318_pc_values, aes(x=PC1, y=PC2) )

#heatmap plus data
windows()
ggplot(validation_all_data, aes(PC1, PC2 )) +
  geom_tile(color='white',aes(fill= EXP_MAX_DISP)) +
  scale_fill_gradient(low='white', high='blue') +
  geom_point(data=n1318_pc_values, aes(x=PC1, y=PC2,z=NULL), col='red') +
  labs(title = "all gnathostomes")
```

```
#plot actual taxa distributions on this heatmap

#hex bin, no heatmap added
# Bin size control + color palette
all_data <- ggplot(n1318_pc_values, aes(x=PC1, y=PC2) ) +
  geom_hex(bins = 20) +
  scale_fill_continuous(type = "viridis") +
  theme_bw() +
  xlim(-0.16726,0.213705) +
  ylim(-0.15065,0.217361)
print (all_data + ggtitle("all gnathostome data"))

#hex bin no heat map, individual clades
gnathostome_clades <- levels(as.factor(n1318_pc_values$Factor))

gg_data <- vector(mode = "list", length = length(gnathostome_clades))

for (i in 1:length(gnathostome_clades)) {
gg_data[[i]] <- ggplot(n1318_pc_values[n1318_pc_values$Factor==gnathostome_clades[i],],
aes(x=PC1, y=PC2) ) +
  geom_hex(bins = 20) +
  scale_fill_continuous(type = "viridis") +
  theme_bw() +
  xlim(-0.16726,0.213705) +
  ylim(-0.15065,0.217361)
}

#scatter plot with heatmap
all_pts_heat <- ggplot(validation_all_data, aes(PC1, PC2 )) +
  geom_tile(color='white',aes(fill= 1/AR)) + #change the fill variable to make different plots
  scale_fill_gradient(low='white', high='blue') +
  geom_point(data=n1318_pc_values[!n1318_pc_values$Factor=="Mammaliaform",],
aes(x=PC1, y=PC2,z=NULL), col='red') +
  labs(title = "non mammaliaforms")
windows()
all_pts_heat

#plotting individual vertebrate clades and compare distributions on a panel figure
gg_data <- vector(mode = "list", length = length(gnathostome_clades))

for (i in 1:length(gnathostome_clades)) {
gg_data[[i]] <- ggplot(validation_all_data, aes(x=PC1, y=PC2) ) +
  geom_tile(color='white',aes(fill= 1/AR)) + #change the fill variable to make different plots
  scale_fill_gradient(low='white', high='blue') +
  geom_point(data=n1318_pc_values[n1318_pc_values$Factor==gnathostome_clades[i],],
aes(x=PC1, y=PC2,z=NULL), col='red') +
```

```
  labs(title = gnathostome_clades[i])
}

figure <-
ggpubr::ggarrange(all_pts_heat,gg_data[[1]],gg_data[[2]],gg_data[[3]],gg_data[[4]],gg_data[[5]]
,
                                    gg_data[[6]],gg_data[[7]],gg_data[[8]],gg_data[[9]],gg_data[[10]],
                                    gg_data[[11]],gg_data[[12]],gg_data[[13]],gg_data[[14]],

          gg_data[[16]],gg_data[[17]],gg_data[[18]],gg_data[[19]],gg_data[[20]],
                                    labels =
c("A","B","C","D","E","F","G","H","I","J","K","L","M","N","O","P","Q","R","S","T"),
                                    ncol = 4, nrow = 5,
                                    #legend = "none"
                                    common.legend = TRUE, legend = "bottom")
figure

ggexport(figure, filename = "figure_gnathostome_MA_heatmap_distributation.pdf")
```

**Appendix S3. R script for generating tree samples using composite topology and stratigraphic ranges.**

```
#script to generate time tree sample using
#1. tree topology for ~890 genera merged from timetree.org extant phylogeny and authors’
vetted fossil phylogeny based on literature review.
#2. FAD and LAD data compiled from Paleobiology Database query via R

#set working directory
setwd()

#libraries for generating time trees
library(phytools)
library(caper)
library(paleobioDB) #to query first and last appearance data from PBDB
library(paleotree)
library(RRphylo)
library(geiger)
#libraries for trait evolution analyses
library(phytools)
library(dplyr)
library(caper)
library(plotrix)
library(pmc) #for bootstrapping of trait evolution parameters
library(geiger) #for treedata function and others
library(motmot)
library(qpcR)
#library for parallel computing
library(parallel)
library(snow)

#master topology with uniform branch lengths and no nodelabels
tree <- read.tree()

#read in merged FADLAD data and trait data
traits <- read.csv("Merged_FADLAD_trait_data.csv",header=T)

#match dataset to tips present in the tree
genus_dataset_and_tree <- comparative.data(tree, traits,
        Genus, warn.dropped=T)

#extract tree vs data
vetted_tree <- genus_dataset_and_tree$phy
vetted_data <- genus_dataset_and_tree$data

#extract genus ranges
```

```
genus_ranges <- as.matrix(vetted_data[,c("fea","lla")])
class(genus_ranges) <- "numeric"
row.names(genus_ranges) <- rownames(vetted_data)

#create time bins in 1 my intervals
genus_range_list <- binTimeData(genus_ranges, int.length = 1) #change int.length to modify bin
width
str(genus_range_list)
taxicDivDisc(genus_range_list)

#890 tips to create time tree from
#Basic Time-Scaling
tree1 <- bin_timePaleoPhy(tree=vetted_tree,timeList=genus_range_list,type="basic",
   ntrees=1,plot=F)
plotTree(tree1, fsize=0.5)

#code below based on rate calibrated time-scaling method tutorial by David Bapst (Bapst, 2014)
#estimate sampling rate
likFun <- make_durationFreqDisc(genus_range_list)
srRes <- optim(parInit(likFun),
likFun,
lower = parLower(likFun),
upper = parUpper(likFun),
method = "L-BFGS-B",
control = list(maxit = 1000000)
)
sRate <- srRes[[1]][2]

# assume extRate = brRate (Foote et al., 1999)
divRate <- srRes[[1]][1]

#generate 100 timetrees
ttrees <- bin_cal3TimePaleoPhy(
vetted_tree,
genus_range_list,
brRate = divRate,
extRate = divRate,
sampRate = sRate,
ntrees = 100,
plot = F)
multiDiv(ttrees)

#save 100 time trees
save(ttrees, file="ttrees_100cal3")
load("ttrees_100cal3")
```

**Appendix S4. R script for plotting traits over phylogenies**

```
#read in PC1, PC2, and EXP_MAX_DISP data
all_genera_3var <- read.csv("n1318_interp_9-8-2021.csv",header=T)

all_genera_3var <- all_genera_3var[,-1]

#take mean of all interpolated biomechanical and PC values by genus
unique_genera_traits <- aggregate(.~Genus+Factor, all_genera_3var, mean)

load("vetted_combined_tree")
vetted_tree

#match dataset to tips present in the tree for EXP_MAX_DISP dataset
genus_dataset_and_tree <- comparative.data(vetted_tree, unique_genera_traits,
        Genus, warn.dropped=F)

#extract tree vs data
vetted_tree <- genus_dataset_and_tree$phy
vetted_data <- genus_dataset_and_tree$data

#extract genus ranges
FADLAD <- read.csv("Merged_FADLAD_trait_data_genus_8-5-2021.csv",header=T)
genus_ranges <- as.matrix(FADLAD[,c("fea","lla")])
class(genus_ranges) <- "numeric"
row.names(genus_ranges) <- FADLAD$Genus

#create time bins in 1 my intervals
genus_range_list <- binTimeData(genus_ranges, int.length = 1) #change int.length to modify bin
width
str(genus_range_list)
taxicDivDisc(genus_range_list)

#Basic Time-Scaling
tree1 <- bin_timePaleoPhy(tree=vetted_tree,timeList=genus_range_list,type="basic",
   ntrees=1,plot=F)
plotTree(tree1, fsize=0.5)

#extract displacement data with rownames
interp_disp <- vetted_data$interp_maxDisp.z
names(interp_disp) <- rownames(genus_dataset_and_tree$data)
interp_ar <- vetted_data$interp_AR.z
names(interp_ar) <- rownames(genus_dataset_and_tree$data)
interp_ma <- vetted_data$interp_MA.z
names(interp_ma) <- rownames(genus_dataset_and_tree$data)
```

```
# Graphing time tree with displacement trait ##
#plot experimental displacement trait over tree
#graphing a fan cladogram with EXP_MAX_DISP mapped as heatmap and major clades labeled
windows()
nodes<-c(1749,1738,1694,1647,1523,1357,1341,912,1483,1514)
labels<-c("placoderms","sharks","bony
fish","amphibia","squamates","dinosaurs","turtles","mammals","pterosaurs","crocs")
obj<-contMap(tree1,interp_disp,plot=FALSE)
obj<-setMap(obj,invert=TRUE) ## invert color map
plot(obj,ftype="off",type="fan",outline=FALSE,
   fsize=c(0,0.8),lwd=2,xlim=c(-700,700),leg.txt="disp")
for(i in 1:length(nodes))
   arc.cladelabels(text=labels[i],node=nodes[i], orientation="horizontal", ln.offset=100)

#same for AR
windows()
nodes<-c(1749,1738,1694,1647,1523,1357,1341,912,1483,1514)
labels<-c("placoderms","sharks","bony
fish","amphibia","squamates","dinosaurs","turtles","mammals","pterosaurs","crocs")
obj<-contMap(tree1,1/interp_ar,plot=FALSE)
obj<-setMap(obj,invert=TRUE) ## invert color map
plot(obj,ftype="off",type="fan",outline=FALSE,
   fsize=c(0,0.8),lwd=2,xlim=c(-700,700),leg.txt="1/AR")
for(i in 1:length(nodes))
   arc.cladelabels(text=labels[i],node=nodes[i], orientation="horizontal", ln.offset=100)

#same for MA
windows()
nodes<-c(1749,1738,1694,1647,1523,1357,1341,912,1483,1514)
labels<-c("placoderms","sharks","bony
fish","amphibia","squamates","dinosaurs","turtles","mammals","pterosaurs","crocs")
obj<-contMap(tree1,interp_ma,plot=FALSE)
obj<-setMap(obj,invert=TRUE) ## invert color map
plot(obj,ftype="off",type="fan",outline=FALSE,
   fsize=c(0,0.8),lwd=2,xlim=c(-700,700),leg.txt="MA")
for(i in 1:length(nodes))
   arc.cladelabels(text=labels[i],node=nodes[i], orientation="horizontal", ln.offset=100)

#adding geologic time scale to cladogram with clade labels, no heatmap#
max(nodeHeights(tree1))
objb<-geo.legend()
r<-max(objb$leg[,1])-objb$leg[,2]
plotTree(tree1,type="fan",fsize=0.7,lwd=1,ftype="off",xlim=c(-700,700))

for(i in 1:nrow(objb$leg)){
   color<-paste(strsplit(objb$colors[i],"")[[1]][1:7],collapse="")
```

```
   draw.circle(0,0,radius=r[i],col=color,border="transparent")
}
par(fg="transparent")
plotTree(tree1,type="fan",add=TRUE,fsize=0.7,lwd=1,ftype="off",xlim=c(-700,700))
par(fg="black")

add.simmap.legend(colors=sapply(objb$colors[rownames(objb$leg)],
   function(x) paste(strsplit(x,"")[[1]][1:7],collapse="")),
   prompt=FALSE,x=0.95*par()$usr[1],y=-0.95*par()$usr[3])
for(i in 1:length(nodes))
   arc.cladelabels(text=labels[i],node=nodes[i], orientation="horizontal", ln.offset=100)
```

**Appendix S5. R script for shape and biomechanical trait disparity analyses in mammals versus nonmammals.**

```
#set working directory
setwd()

#libraries
library(dispRity)

## First part is an analysis of shape disparity in PC values of Elliptic Fourier harmonics of gnathostome jaw shapes, comparing mammals vs nonmammals

#load PC values for n1325 2nd morphospace analysis dataset
data <- read.csv(file="0utput_n1325_PCA_trimmed_PCscores_labeled_disparity.csv",header=T)

n1325_matrix <- data[,3:42]
two_groups <- list("mammals" = which(data[,1]=="Mammaliaform"), "nonmammals" =
which(data[,1]=="Nonmammal"))

#
disparity_data <- dispRity.per.group(n1325_matrix,
                              group = two_groups,
                              metric = c(sum, variances))

summary(disparity_data)

plot(disparity_data)

## Testing for a difference between the groups
test.dispRity(disparity_data, test = wilcox.test, details = TRUE)
#mammals exhibit significantly lower disparity compared to non-mammals

## Bootstrapping 1000 times with full rarefaction: ALL PCs
mamm_nonmamm <- custom.subsets(n1325_matrix,
        group= list("mammals" = which(data[,1]=="Mammaliaform"), "nonmammals" =
which(data[,1]=="Nonmammal")))
boot_data <- boot.matrix(mamm_nonmamm, bootstraps = 1000,
        rarefaction = TRUE)
#disparity calculation using median distances between observations with centroids
disparity_boot_data <- dispRity(boot_data, metric = c(median, centroids))
summary(disparity_boot_data)
plot(disparity_boot_data)
## Running a wilcox test
test.dispRity(disparity_boot_data, test = wilcox.test, details = TRUE)
#mammals have significantly lower disparity across all PCs
```

```
#below: also test Pc1, pc2, ma, ar, and displacement disparity

## Bootstrapping 1000 times with full rarefaction: PCs 1 & 2
mamm_nonmamm_pc12 <- custom.subsets(n1325_matrix[,1:2],
        group= list("mammals" = which(data[,1]=="Mammaliaform"), "nonmammals" =
which(data[,1]=="Nonmammal")))
boot_data_pc12 <- boot.matrix(mamm_nonmamm_pc12, bootstraps = 1000,
        rarefaction = TRUE)
#disparity calculation using median distances between observations with centroids
disparity_boot_data_pc12 <- dispRity(boot_data_pc12, metric = c(median, centroids))
summary(disparity_boot_data_pc12)
plot(disparity_boot_data_pc12)
## Running a wilcox test
test.dispRity(disparity_boot_data_pc12, test = wilcox.test, details = TRUE)
#mammals have significantly higher disparity in PC1 and PC2

# Second part is using interpolated data for functional trait disparity analyses ##
#load PC values for n1325 2nd morphospace analysis dataset
data_for_genus <- read.csv(file="n1318_interp_9-8-2021_mamm_vs_nonmamm.csv",header=T)
data_for_genus <- data_for_genus[,-c(1,3)]

#take mean of all interpolated biomechanical and PC values by genus
unique_genera_traits <- aggregate(.~Genus+Factor2, data_for_genus, mean)

## Bootstrapping 1000 times with full rarefaction: MA
mamm_nonmamm_ma <- custom.subsets(as.matrix(unique_genera_traits[,7]),
        group= list("mammals" = which(unique_genera_traits[,2]=="Mammal"),
                "nonmammals" = which(unique_genera_traits[,2]=="Nonmammal")))
boot_data_ma <- boot.matrix(mamm_nonmamm_ma, bootstraps = 1000,
        rarefaction = TRUE)
#disparity calculation using median distances between observations with centroids
disparity_boot_data_ma <- dispRity(boot_data_ma, metric = c(median, centroids))
summary(disparity_boot_data_ma)
plot(disparity_boot_data_ma)
## Running a wilcox test
test.dispRity(disparity_boot_data_ma, test = wilcox.test, details = TRUE)
#mammals have significantly lower disparity in MA

## Bootstrapping 1000 times with full rarefaction: AR
mamm_nonmamm_ar <- custom.subsets(as.matrix(unique_genera_traits[,6]),
        group= list("mammals" = which(unique_genera_traits[,2]=="Mammal"),
                "nonmammals" = which(unique_genera_traits[,2]=="Nonmammal")))
boot_data_ar <- boot.matrix(mamm_nonmamm_ar, bootstraps = 1000,
        rarefaction = TRUE)
#disparity calculation using median distances between observations with centroids
disparity_boot_data_ar <- dispRity(boot_data_ar, metric = c(median, centroids))
```

```
summary(disparity_boot_data_ar)
plot(disparity_boot_data_ar)
## Running a wilcox test
test.dispRity(disparity_boot_data_ar, test = wilcox.test, details = TRUE)
#mammals have significantly lower disparity in AR

## Bootstrapping 1000 times with full rarefaction: stiffness
mamm_nonmamm_disp <- custom.subsets(as.matrix(unique_genera_traits[,5]),
        group= list("mammals" = which(unique_genera_traits[,2]=="Mammal"),
                "nonmammals" = which(unique_genera_traits[,2]=="Nonmammal")))
boot_data_disp <- boot.matrix(mamm_nonmamm_disp, bootstraps = 1000,
        rarefaction = TRUE)
#disparity calculation using median distances between observations with centroids
disparity_boot_data_disp <- dispRity(boot_data_disp, metric = c(median, centroids))
summary(disparity_boot_data_disp)
plot(disparity_boot_data_disp)
## Running a wilcox test
test.dispRity(disparity_boot_data_disp, test = wilcox.test, details = TRUE)
#mammals have significantly lower disparity in displacement

## Bootstrapping 1000 times with full rarefaction: PC1&PC2
mamm_nonmamm_pc12 <- custom.subsets(as.matrix(unique_genera_traits[,3:4]),
        group= list("mammals" = which(unique_genera_traits[,2]=="Mammal"),
                "nonmammals" = which(unique_genera_traits[,2]=="Nonmammal")))
boot_data_pc12 <- boot.matrix(mamm_nonmamm_pc12, bootstraps = 1000,
        rarefaction = TRUE)
#disparity calculation using median distances between observations with centroids
disparity_boot_data_pc12 <- dispRity(boot_data_pc12, metric = c(median, centroids))
summary(disparity_boot_data_pc12)
plot(disparity_boot_data_pc12)
## Running a wilcox test
test.dispRity(disparity_boot_data_pc12, test = wilcox.test, details = TRUE)

### make violin and box plots to visualize data distributional differences between mammals and
nonmammals

#libraries
library(ggplot2)
library(ggpubr)

#boxplots to visualize mammal vs nonmammal differences
boxplot(PC1~Factor, data=data, main="PC1 scores", xlab="Clade", ylab="Score")

#x labels
group_lab <- c("Mammal","Nonmammal")
```

```
#violin plot
p <- ggplot(data, aes(x=Factor, y=PC1, fill=Factor)) +
  geom_violin(show.legend = FALSE) + labs(y= "PC1 score", x = NULL) +
scale_x_discrete(labels= group_lab)
pj <- p + geom_jitter(shape=16, position=position_jitter(0.4), show.legend = FALSE)+
scale_fill_brewer(palette="Dark2")

p2 <- ggplot(data, aes(x=Factor, y=PC2, fill=Factor)) +
  geom_violin(show.legend = FALSE) + labs(y= "PC2 score", x = NULL) +
scale_x_discrete(labels= group_lab)
p2j <- p2 + geom_jitter(shape=16, position=position_jitter(0.4), show.legend = FALSE)+
scale_fill_brewer(palette="Dark2")

## interpolated data for functional trait disparity analyses ##
#load PC values for n1325 2nd morphospace analysis dataset
data_for_genus <- read.csv(file="n1318_interp_9-8-2021_mamm_vs_nonmamm.csv",header=T)
data_for_genus <- data_for_genus[,-c(1,3)]

#take mean of all interpolated biomechanical and PC values by genus
unique_genera_traits <- aggregate(.~Genus+Factor2, data_for_genus, mean)

p3 <- ggplot(unique_genera_traits, aes(x=Factor2, y=interp_MA.z, fill=Factor2)) +
  geom_violin(show.legend = FALSE) + labs(y= "Mechanical Advantage", x = NULL) +
scale_x_discrete(labels= group_lab)
p3j <- p3 + geom_jitter(shape=16, position=position_jitter(0.4), show.legend = FALSE)+
scale_fill_brewer(palette="Dark2")

p4 <- ggplot(unique_genera_traits, aes(x=Factor2, y=interp_AR.z, fill=Factor2)) +
  geom_violin(show.legend = FALSE) + labs(y= "Aspect Ratio", x = NULL) +
scale_x_discrete(labels= group_lab)
p4j <- p4 + geom_jitter(shape=16, position=position_jitter(0.4), show.legend = FALSE)+
scale_fill_brewer(palette="Dark2")

#clean violin plots 2 x 2
ggarrange(p, p2, p3, p4,
        labels = c("A", "B", "C", "D"),
        ncol = 2, nrow = 2,
          common.legend = TRUE, legend = "bottom")

#jitteredviolin plots 2 x 2
ggarrange(pj, p2j, p3j, p4j,
        labels = c("A", "B", "C", "D"),
        ncol = 2, nrow = 2,
          common.legend = TRUE, legend = "bottom")
```

**Appendix S6. Gene sequence alignment, tree generation, and relative evolutionary rate analyses**

```
#Install required packages
if (!requireNamespace("BiocManager", quietly = TRUE))
   install.packages("BiocManager")

BiocManager::install("ShortRead")
BiocManager::install("DECIPHER")

library(ShortRead) #to read fasta files
library(DECIPHER) #to conduct clustering of sequences

install.packages("ape")
install.packages("phangorn")
install.packages("seqinr")
library(ape)
library(phangorn)
library(seqinr)

#set working directory
setwd(choose.dir())

#script below is based on tutorial from https://www.molecularecologist.com/2016/02/26/quick-and-dirty-tree-building-in-r/

#read in aligned sequences (Alignment conducted in clustalX with .fasta file outputs)
alx3 <- read.dna("Alx3_sub_mesquite_taxa.fasta", format="fasta")
bmp3 <- read.dna("Bmp3_sub_mesquite_taxa.fasta", format="fasta")
cited1 <- read.dna("Cited1_sub_mesquite_taxa.fasta", format="fasta")
dlx1 <- read.dna("Dlx1_sub_mesquite_taxa.fasta", format="fasta")
dlx2 <- read.dna("Dlx2_sub_mesquite_taxa.fasta", format="fasta")
dlx3 <- read.dna("Dlx3_sub_mesquite_taxa.fasta", format="fasta")
dlx4 <- read.dna("Dlx4_sub_mesquite_taxa.fasta", format="fasta")
dlx5 <- read.dna("Dlx5_sub_mesquite_taxa.fasta", format="fasta")
dlx6 <- read.dna("Dlx6_sub_mesquite_taxa.fasta", format="fasta")
gbx2 <- read.dna("Gbx2_sub_mesquite_taxa.fasta", format="fasta")
gsc <- read.dna("Gsc_sub_mesquite_taxa.fasta", format="fasta")
hand1 <- read.dna("Hand1_sub_mesquite_taxa.fasta", format="fasta")
hand2 <- read.dna("Hand2_sub_mesquite_taxa.fasta", format="fasta")
hgf <- read.dna("Hgf_sub_mesquite_taxa.fasta", format="fasta")
meis2 <- read.dna("Meis2_sub_mesquite_taxa.fasta", format="fasta")
nkx3 <- read.dna("Nkx3_2_sub_mesquite_taxa.fasta", format="fasta")
plagl1 <- read.dna("Plagl1_sub_mesquite_taxa.fasta", format="fasta")
rgs5 <- read.dna("Rgs5_sub_mesquite_taxa.fasta", format="fasta")
unc5c <- read.dna("Unc5c_mesquite_taxa.fasta", format="fasta")
```

```
#convert format to DNA
alx3_phyDat <- phyDat(alx3, type = "DNA", levels = NULL)
bmp3_phyDat <- phyDat(bmp3, type = "DNA", levels = NULL)
cited1_phyDat <- phyDat(cited1, type = "DNA", levels = NULL)
dlx1_phyDat <- phyDat(dlx1, type = "DNA", levels = NULL)
dlx2_phyDat <- phyDat(dlx2, type = "DNA", levels = NULL)
dlx3_phyDat <- phyDat(dlx3, type = "DNA", levels = NULL)
dlx4_phyDat <- phyDat(dlx4, type = "DNA", levels = NULL)
dlx5_phyDat <- phyDat(dlx5, type = "DNA", levels = NULL)
dlx6_phyDat <- phyDat(dlx6, type = "DNA", levels = NULL)
gbx2_phyDat <- phyDat(gbx2, type = "DNA", levels = NULL)
gsc_phyDat <- phyDat(gsc, type = "DNA", levels = NULL)
hand1_phyDat <- phyDat(hand1, type = "DNA", levels = NULL)
hand2_phyDat <- phyDat(hand2, type = "DNA", levels = NULL)
hgf_phyDat <- phyDat(hgf, type = "DNA", levels = NULL)
meis2_phyDat <- phyDat(meis2, type = "DNA", levels = NULL)
nkx3_phyDat <- phyDat(nkx3, type = "DNA", levels = NULL)
plagl1_phyDat <- phyDat(plagl1, type = "DNA", levels = NULL)
rgs5_phyDat <- phyDat(rgs5, type = "DNA", levels = NULL)
unc5c_phyDat <- phyDat(unc5c, type = "DNA", levels = NULL)

#build distance matrices using maximum likelihood
dist_alx3 <- dist.ml(alx3_phyDat)
dist_bmp3 <- dist.ml(bmp3_phyDat)
dist_cited1 <- dist.ml(cited1_phyDat)
dist_dlx1 <- dist.ml(dlx1_phyDat)
dist_dlx2 <- dist.ml(dlx2_phyDat)
dist_dlx3 <- dist.ml(dlx3_phyDat)
dist_dlx4 <- dist.ml(dlx4_phyDat)
dist_dlx5 <- dist.ml(dlx5_phyDat)
dist_dlx6 <- dist.ml(dlx6_phyDat)
dist_gbx2 <- dist.ml(gbx2_phyDat)
dist_gsc <- dist.ml(gsc_phyDat)
dist_hand1 <- dist.ml(hand1_phyDat)
dist_hand2 <- dist.ml(hand2_phyDat)
dist_hgf <- dist.ml(hgf_phyDat)
dist_meis2 <- dist.ml(meis2_phyDat)
dist_nkx3 <- dist.ml(nkx3_phyDat)
dist_plagl1 <- dist.ml(plagl1_phyDat)
dist_rgs5 <- dist.ml(rgs5_phyDat)
dist_unc5c <- dist.ml(unc5c_phyDat)

#build UPGMA phenograms based on distance matrices
alx3_UPGMA <- upgma(dist_alx3)
plot(alx3_UPGMA)
```

```
write.tree(alx3_UPGMA,file="alx3_upgma_tree.nwk")

alx3_UPGMA <- upgma(dist_alx3)
bmp3_UPGMA <- upgma(dist_bmp3)
cited1_UPGMA <- upgma(dist_cited1)
dlx1_UPGMA <- upgma(dist_dlx1)
dlx2_UPGMA <- upgma(dist_dlx2)
dlx3_UPGMA <- upgma(dist_dlx3)
dlx4_UPGMA <- upgma(dist_dlx4)
dlx5_UPGMA <- upgma(dist_dlx5)
dlx6_UPGMA <- upgma(dist_dlx6)
gbx2_UPGMA <- upgma(dist_gbx2)
gsc_UPGMA <- upgma(dist_gsc)
hand1_UPGMA <- upgma(dist_hand1)
hand2_UPGMA <- upgma(dist_hand2)
hgf_UPGMA <- upgma(dist_hgf)
meis2_UPGMA <- upgma(dist_meis2)
nkx3_UPGMA <- upgma(dist_nkx3)
plagl1_UPGMA <- upgma(dist_plagl1)
rgs5_UPGMA <- upgma(dist_rgs5)
unc5c_UPGMA <- upgma(dist_unc5c)

UPGMA_trees <-
c(alx3_UPGMA,bmp3_UPGMA,cited1_UPGMA,dlx1_UPGMA,dlx2_UPGMA,dlx3_UPGMA,
dlx4_UPGMA,
        dlx5_UPGMA,dlx6_UPGMA,gbx2_UPGMA,gsc_UPGMA,hand1_UPGMA,hand2_UP
GMA,hgf_UPGMA,
        meis2_UPGMA,nkx3_UPGMA,plagl1_UPGMA,rgs5_UPGMA,unc5c_UPGMA)

write.tree(UPGMA_trees,file="19genes_upgma_gene_trees.nwk")

#save compiled trees
tree_tips_compiled <- matrix(nrow=105, ncol=19)
for (i in 1: length(UPGMA_trees)) {
        for (j in 1:105) {
        tree_tips_compiled[j,i]<-UPGMA_trees[[i]]$tip.label[j]
        }
}

write.csv(tree_tips_compiled, file="19_Gene_tree_tip_labels.csv")


#####
RERCONVERGE SECTION
#####
#these packages were installed ad hoc as errors popped up during devtools install
```

```
install.packages("testthat")
install.packages("processx")
install.packages("gplots")
install.packages("fansi")
install.packages("backports")
install.packages("remotes")
#then install devtools
install.packages("devtools")
#7-15-2021 additional package required by RERconverge installer
#install from  here: https://cran.r-project.org/bin/windows/Rtools/
writeLines('PATH="${RTOOLS40_HOME}\\usr\\bin;${PATH}"', con = "~/.Renviron")
#then verify that make can be found, using:
Sys.which("make")

#these libraries installed manually in order for devtools to load
library(testthat)
library(processx)
library(gplots)
library(fansi)
library(backports)
library(devtools)

#this library turns off warning during github install of version differences
library(remotes)
#Turn off warning-error-conversion, because the tiniest warning stops installation
Sys.setenv("R_REMOTES_NO_ERRORS_FROM_WARNINGS" = "true")

#installation of RERconverge from github, with dependencies
install_github("nclark-lab/RERconverge", dependencies = T)
#load
library(RERconverge)

#read master tree
master_tree <- read.newick("19genes_tip_labels_105taxa_master_tree.nwk")
gene_trees <- read.tree(file="19genes_upgma_gene_trees_named.nwk")
#comapre gene tree to master tree to verify tip labels
comparePhylo(master_tree,gene_trees[[1]])

#read gene trees
gene_trees_n19 <- readTrees(file="19genes_upgma_gene_trees_named.nwk",
        minTreesAll=15,minSpecs=100)

RERw = getAllResiduals(gene_trees_n19, useSpecies=gene_trees_n19$masterTree$tip.label,
        transform ="none", weighted = T, scale = T)

#make average and gene tree plots
```

```
#plot average tree
avgtree=plotTreeHighlightBranches(gene_trees_n19$masterTree,
hlspecies=c("Canis_lupus","Cebus_capucinus"),
        hlcols=c("blue","red"), main="Average tree")

#plot individual gene tree
bmp3tree=plotTreeHighlightBranches(gene_trees_n19$trees$BMP3,
hlspecies=c("Canis_lupus","Cebus_capucinus"),
        hlcols=c("blue","red"), main="BMP3 tree")

# according to https://github.com/nclark-lab/RERconverge/issues/58
#all gene trees need to have the same topology, so they provided the function below to take in
#one master topology and generate gene tree branch length differences based on that topology.

estimatePhangornTree = function(alnfile, treefile, submodel="LG", type = "AA",
                        format = "fasta", k=4, ...) {
  #Generates distance-based tree using submodel for substitutions
  #Read in pruned alignment and pruned master tree
  #If species lists are different, prune both to be the same.
  #read in the alignment
  #zjt commented out#alnPhyDat = read.phyDat(alnfile, type = type, format = format)
  alnPhyDat = alnfile
  #read in the treefile
  #zjt commented out#genetree = read.tree(treefile)
  genetree = treefile
  #eliminate species in the alignment but not the tree and vice versa
  inboth = intersect(names(alnPhyDat),genetree$tip.label)
  todropg = genetree$tip.label[genetree$tip.label %in% inboth == FALSE]
  if (length(todropg) > 0) {
    genetree = drop.tip(genetree, todropg)
  }
  if (length(inboth) < length(names(alnPhyDat))) {
    alnPhyDat = subset(alnPhyDat, subset = inboth)
  }
  #unroot the tree
  genetree = unroot(genetree)
  #just in case, set all branches to 1 first (pml abhors a vacuum... or a zero)
  genetree$edge.length = c(rep(1,length(genetree$edge.length)))
  #Run distance estimation using submodel
  #generate an initial pml tree
  lgptree = pml(genetree, alnPhyDat, model = submodel, k = k, rearrangement="none", ...)
  #generate a tree
  #use capture.output to suppress optimization output?
  lgopttree = optim.pml(lgptree,optInv=T,optGamma=T,optEdge=T,rearrangement="none",
                    model=submodel, ...)
  return(list("results.init"=lgptree,"results.opt"=lgopttree, "tree.opt"=lgopttree$tree))
```

```
}

#estimating gene branch lengths using timetree master tree
master_tree_fasta <-
read.newick(file="19genes_tip_labels_105taxa_master_tree_match_fasta.nwk")

alx3_phan <- estimatePhangornTree(alx3_phyDat, master_tree_fasta, type="DNA",
submodel="JC")
bmp3_phan <- estimatePhangornTree(bmp3_phyDat, master_tree_fasta, type="DNA",
submodel="JC")
cited1_phan <- estimatePhangornTree(cited1_phyDat, master_tree_fasta, type="DNA",
submodel="JC")
dlx1_phan <- estimatePhangornTree(dlx1_phyDat, master_tree_fasta, type="DNA",
submodel="JC")
dlx2_phan <- estimatePhangornTree(dlx2_phyDat, master_tree_fasta, type="DNA",
submodel="JC")
dlx3_phan <- estimatePhangornTree(dlx3_phyDat, master_tree_fasta, type="DNA",
submodel="JC")
dlx4_phan <- estimatePhangornTree(dlx4_phyDat, master_tree_fasta, type="DNA",
submodel="JC")
dlx5_phan <- estimatePhangornTree(dlx5_phyDat, master_tree_fasta, type="DNA",
submodel="JC")
dlx6_phan <- estimatePhangornTree(dlx6_phyDat, master_tree_fasta, type="DNA",
submodel="JC")
gbx2_phan <- estimatePhangornTree(gbx2_phyDat, master_tree_fasta, type="DNA",
submodel="JC")
gsc_phan <- estimatePhangornTree(gsc_phyDat, master_tree_fasta, type="DNA",
submodel="JC")
hand1_phan <- estimatePhangornTree(hand1_phyDat, master_tree_fasta, type="DNA",
submodel="JC")
hand2_phan <- estimatePhangornTree(hand2_phyDat, master_tree_fasta, type="DNA",
submodel="JC")
hgf_phan <- estimatePhangornTree(hgf_phyDat, master_tree_fasta, type="DNA",
submodel="JC")
meis2_phan <- estimatePhangornTree(meis2_phyDat, master_tree_fasta, type="DNA",
submodel="JC")
nkx3_phan <- estimatePhangornTree(nkx3_phyDat, master_tree_fasta, type="DNA",
submodel="JC")
plagl1_phan <- estimatePhangornTree(plagl1_phyDat, master_tree_fasta, type="DNA",
submodel="JC")
rgs5_phan <- estimatePhangornTree(rgs5_phyDat, master_tree_fasta, type="DNA",
submodel="JC")
unc5c_phan <- estimatePhangornTree(unc5c_phyDat, master_tree_fasta, type="DNA",
submodel="JC")
```

```
phan_trees <-
c(alx3_phan$tree,bmp3_phan$tree,cited1_phan$tree,dlx1_phan$tree,dlx2_phan$tree,
                          dlx3_phan$tree,dlx4_phan$tree,

        dlx5_phan$tree,dlx6_phan$tree,gbx2_phan$tree,gsc_phan$tree,hand1_phan$tree,
                          hand2_phan$tree,hgf_phan$tree,

        meis2_phan$tree,nkx3_phan$tree,plagl1_phan$tree,rgs5_phan$tree,unc5c_phan$tree)
write.tree(phan_trees,file="19genes_phan_gene_trees.nwk")

fc=file("19genes_phan_gene_trees.nwk", "wt")
gene_list <-
c("ALX3","BMP3","CITED1","DLX1","DLX2","DLX3","DLX4","DLX5","DLX6","GBX2","
GSC",  "HAND1","HAND2","HGF","MEIS2","NKX3","PLAGL1","RGS5","UNC5C")

  for(i in 1:length(phan_trees)){
    writeLines(paste(gene_list[i], write.tree(phan_trees[[i]]), sep = "\t"), fc)
  }
  close(fc)

#read master tree
master_tree_fasta <-
read.newick(file="19genes_tip_labels_105taxa_master_tree_match_fasta.nwk")

#read gene trees
gene_trees_n19 <- readTrees(file="19genes_phan_gene_trees.nwk",
        minTreesAll=19,minSpecs=105)

RERw = getAllResiduals(gene_trees_n19, useSpecies=gene_trees_n19$masterTree$tip.label,
        transform ="sqrt", weighted = T, scale = T)

saveRDS(RERw, file="RERw.rds")
loadRERw = readRDS("RERw.rds")

#make average and gene tree plots
par(mfrow=c(1,2))
#plot average tree
avgtree=plotTreeHighlightBranches(gene_trees_n19$masterTree,
hlspecies=c("Canis_lupus_dingo","Cebus_imitator"),
        hlcols=c("blue","red"), main="Average tree")

#plot individual gene tree
bmp3tree=plotTreeHighlightBranches(gene_trees_n19$trees$bmp3,
hlspecies=c("Canis_lupus_dingo","Cebus_imitator"),
        hlcols=c("blue","red"), main="BMP3 tree")
```

```
#plot RERs
par(mfrow=c(1,1))
phenvExample <-
foreground2Paths(c("Heterocephalus_glaber","Mus_musculus"),gene_trees_n19,
                    clade="terminal")
plotRers(RERw,"bmp3",phenv=phenvExample)

multirers = returnRersAsTreesAll(gene_trees_n19,RERw)
write.tree(multirers, file='n19_genes_RERs.nwk', tree.names=TRUE)

#visualize RERs along branches as a heatmap
new_alx3_rers = treePlotRers(treesObj=gene_trees_n19, rermat=RERw, index="alx3", type="c",
          nlevels=20, figwid=1)

#import vetted unique genera from gene trees
unique_genera <- read.csv("master_tree_fasta_tips_unique_genera.csv",header=F)

#import continuous trait data
cont_trait_data <- read.csv("n1318_interp_9-8-2021.csv", row.names=1)

#subsetting trait_data with unique genera from genes only
subset_cont_trait <- cont_trait_data[cont_trait_data[,1] %in% unique_genera[,1],]
cont_trait_trimmed <- aggregate(subset_cont_trait[, 3:7], list(subset_cont_trait$Genus), mean)
write.csv(cont_trait_trimmed,file="gene_tree_cont_trait.csv")


##################################
#CORE RER ANALYSES FROM HERE ON#
#importing vetted trimmed and fused cont trait dataset
cont_trait_fused <- read.csv("gene_tree_cont_trait_corresponded_fused.csv", row.names=1)

#read master tree
master_tree_fasta <-
read.newick(file="19genes_tip_labels_105taxa_master_tree_match_fasta.nwk")

#read gene trees
gene_trees_n19 <- readTrees(file="19genes_phan_gene_trees.nwk",
          minTreesAll=19,minSpecs=105)

#cont.trait path to rer: 5 traits
tip.vals <- t(as.vector(cont_trait_fused[,3,drop=F])) #1=pc1, 2=pc2, 3=disp, 4=ar, 5=ma#
names(tip.vals) <- colnames(tip.vals)
charpaths=char2Paths(tip.vals, gene_trees_n19)

#get residuals to vetted species only
RERw = getAllResiduals(gene_trees_n19, useSpecies=names(tip.vals),
```

```
        transform ="log", weighted = F, scale = T)

#saveRDS(RERw, file="RERw.rds")
#RERw = readRDS("RERw.rds")

#cont trait to genome element table
res <-correlateWithContinuousPhenotype(RERw, charpaths,
        min.sp =10, winsorizeRER=3, winsorizetrait=3)

res[order(res$P),]

#correlation by gene
x=charpaths
y=RERw['ALX3',]
pathnames=namePathsWSpecies(gene_trees_n19$masterTree)
names(y)=pathnames
plot(x,y, cex.axis=1, cex.lab=1, cex.main=1, xlab="Weight Change",
        ylab="Evolutionary Rate", main="Gene ALX3 Pearson Correlation w AR", pch=19,
cex=0.5, xlim=c(-2,2))
text(x,y, labels=names(y), pos=4, cex=0.75)
abline(lm(y~x), col='red',lwd=3)

##
#visual examination of gene specific taxon evo rates#
##
#plot individual gene tree
gene_taxon_tree=plotTreeHighlightBranches(gene_trees_n19$trees$alx3,
        hlspecies=c("Heterocephalus_glaber","Mus_musculus"),
        hlcols=c("blue","red"), main="alx3 tree")

#plot RERs
par(mfrow=c(1,1))
phenvExample <-
foreground2Paths(c("Heterocephalus_glaber","Mus_musculus"),gene_trees_n19,
                clade="terminal")
plotRers(RERw,"alx3",phenv=phenvExample)

#plot RERs as tree
par(mfrow=c(1,1))
bend3rers = returnRersAsTree(gene_trees_n19, RERw,"alx3", plot =TRUE,
phenv=phenvExample)

#visualize RERs along branches as a heatmap
new_rgs5_rers = treePlotRers(treesObj=gene_trees_n19, rermat=RERw, index="rgs5", type="c",
        nlevels=20, figwid=1)
#laurasiatheres are reconstructed as having ancestrally accelerated evo rate for alx3
```

```
#euarchontoglires have ancestrally accelerated plagl1 evo rate.
# both clades recon to have relative low rates of bmp3 evo rate
#no ancestral acceleration/decrease in rgs5 evo rate
###################################
#4 genes alx3, plagl1, bmp3, dlx5 are significantly (by p.adj) correlated to AR evo rate.
#no genes are sig correlated to MA evo rate.
#3 genes alx3, rgs5, and bmp3 are sig correlated w displacement evo rate
```

**Appendix S7. R Script for plotting network graph of gene-biomechanics correlations**

```
#Set working directory
setwd(choose.dir())

#required library
library(igraph)

#import gene-morphology significant linkage quantities
gene_morph <- read.csv("gene_morphology_edge_list.csv", header=FALSE)

g <- graph.data.frame(gene_morph, directed=FALSE)

bipartite.mapping(g)
V(g)$type <- bipartite_mapping(g)$type

plot(g)

V(g)$color <- ifelse(V(g)$type, "lightblue", "salmon")
V(g)$shape <- ifelse(V(g)$type, "circle", "square")
E(g)$color <- "black"
E(g)$weight <- edge.betweenness(g)

V(g)$size <- 22
plot(g, vertex.label.cex = 0.8, vertex.label.color = "black")

#simplified edges
g2 = simplify(g)
V(g2)$color <- ifelse(V(g2)$type, "lightblue", "salmon")
V(g2)$shape <- ifelse(V(g2)$type, "circle", "square")
E(g2)$color <- "black"
E(g2)$weight = sapply(E(g2), function(e) {
   length(all_shortest_paths(g, from=ends(g2, e)[1], to=ends(g2, e)[2])$res) } )
V(g2)$size <- 23
l <- layout.fruchterman.reingold(g2,niter=500,area=vcount(g2)^2.3,repulserad=vcount(g2)^2.8)

plot(g2, vertex.label.cex = 0.8, vertex.label.color = "black",
edge.width=E(g2)$weight*2,layout=l)
```

**Appendix S8. Entanglement analysis for pairwise gene tree comparisons of phenogram similarity (script shown is for one example pairwise analysis)**

```
#set working directory
setwd(choose.dir())

#load libraries
library(ggdendro)
library(ggplot2)
library(corrplot)
library(datelife)
library(rotl)
library(gmodels)
library(ape)
library (maps)
library(phytools)
library(geiger)
library(cluster)
library(dendextend)
library(MASS)
library(mvtnorm)
library(caper)
library(TreeTools)
library(zoom)

#read and parse phylogenetic data; these are phenograms already built using UPGMA, NJ, etc.
methods.
treeA <-read.newick("Hand2_sequence_branch.nwk")
treeB <-read.newick("Rgs5_sequence_branch.nwk")
#repeat for additional trees

#keep matching tips only
treeA<-drop.tip(treeA,treeA$tip.label[-na.omit(match(treeB$tip.label, treeA$tip.label))])
treeB<-drop.tip(treeB,treeB$tip.label[-na.omit(match(treeA$tip.label, treeB$tip.label))])

#setting negative branch lengths to zero
#(only necessary to correct for in NJ tree building method)
treeA$edge.length[treeA$edge.length<0]<-0
treeB$edge.length[treeB$edge.length<0]<-0

#dichotomize trees
treeA <- multi2di(treeA, random= FALSE)
treeB <- multi2di(treeB, random= FALSE)

#export tips for loading into timetree to identify outgroup
write.table(treeA$tip, file="Hand2_tips.txt",row.names=F,col.names=F)
```

```
write.table(treeB$tip, file="Rgs5_tips.txt",row.names=F,col.names=F)

#(then, process the txt file and load into timetree.org to obtain outgroup)

#To find out the tip number for given species with name "species" type
#treeA
which(treeA$tip.label=="'Acanthopagrus_latus'")

#treeB
which(treeB$tip.label=="'Acanthopagrus_latus'")

#look the position of the taxa in the trees
zoom(treeA, c(109:110,112:123,125:129), cex = 0.75, main = "treeA", subtree = TRUE
windows()
zoom(treeB, c(135:140,157:160,162:170), cex = 0.75, main = "treeB", subtree = TRUE)

#then drop from both files those outgroup taxa nested with other taxa in treeB
treeA_drop<-drop.tip(treeA,c("'Notolabrus_celidotus'", "'Danio_rerio'",
"'Electrophorus_electricus'", "'Pygocentrus_nattereri'", "'Colossoma_macropomum'",
"'Pangasianodon_hypophthalmus'"))
treeB_drop<-drop.tip(treeB,c("'Notolabrus_celidotus'", "'Danio_rerio'",
"'Electrophorus_electricus'", "'Pygocentrus_nattereri'", "'Colossoma_macropomum'",
"'Pangasianodon_hypophthalmus'"))

#Check and verify the inconsistent taxa were removed

#look for the most recent common ancester of the outgroup members using different functions
getMRCA(treeA_drop,c(109:110,112:122))
findMRCA(treeA_drop,c(109:110,112:122))

getMRCA(treeB_drop,c(151:154,156:164))
findMRCA(treeB_drop,c(151:154,156:164))

#test that the clade includes the same number of taxa
clade.members(340, treeA_drop)
clade.members(340, treeA_drop, tip.labels=TRUE)

clade.members(380, treeB_drop)
clade.members(380, treeB_drop, tip.labels=TRUE)

#Export the results to notepad, then open it in Excel to order alphabetically the names and
compare them

#root the tree
#choose random outgroup to test whether outgroup definition changes outcomes
treeA_rooted <- root(treeA_drop,node=340,resolve.root=T)
```

```
treeB_rooted <- root(treeB_drop,node=380,resolve.root=T)

#look for the root node
RootNode(treeA_rooted)
RootNode(treeB_rooted)

#remove node labels
treeA_rooted$node.label <- NULL
treeB_rooted$node.label <- NULL

#renumbering tree to conform to phylo format
treeA_rooted_renum <- TreeTools::Renumber(treeA_rooted)
treeB_rooted_renum <- TreeTools::Renumber(treeB_rooted)

#making phylogeny into ultrametric dendrogram
treeA_rooted_ultra <- chronos(treeA_rooted_renum,lambda=0)
treeB_rooted_ultra <- chronos(treeB_rooted_renum,lambda=0)

is.ultrametric(treeA_rooted_ultra)
is.ultrametric(treeB_rooted_ultra)

#function to compare trees to make sure both have same tips and are rooted and ultrametric
comparePhylo(treeA_rooted_ultra,treeB_rooted_ultra)
is.binary(treeA_rooted_ultra)
is.binary(treeB_rooted_ultra)

# Create dendrograms
dendA <- as.hclust (treeA_rooted_ultra)
dendB <- as.hclust (treeB_rooted_ultra)
dendA <- as.dendrogram (dendA)
dendB <- as.dendrogram (dendB)

#can try plotting to make sure trees/dendrograms look ok
plot(dendA)
windows()
plot(dendB)

# Align and plot two dendrograms side by side
dendAB <- dendlist(dendA, dendB)
dendAB_untangled <- dendextend::untangle(dendAB, method = "step1side") # Find the best
alignment layout
tanglegram(dendAB_untangled)                    # Draw the two dendrograms
entanglement(dendAB_untangled)
```

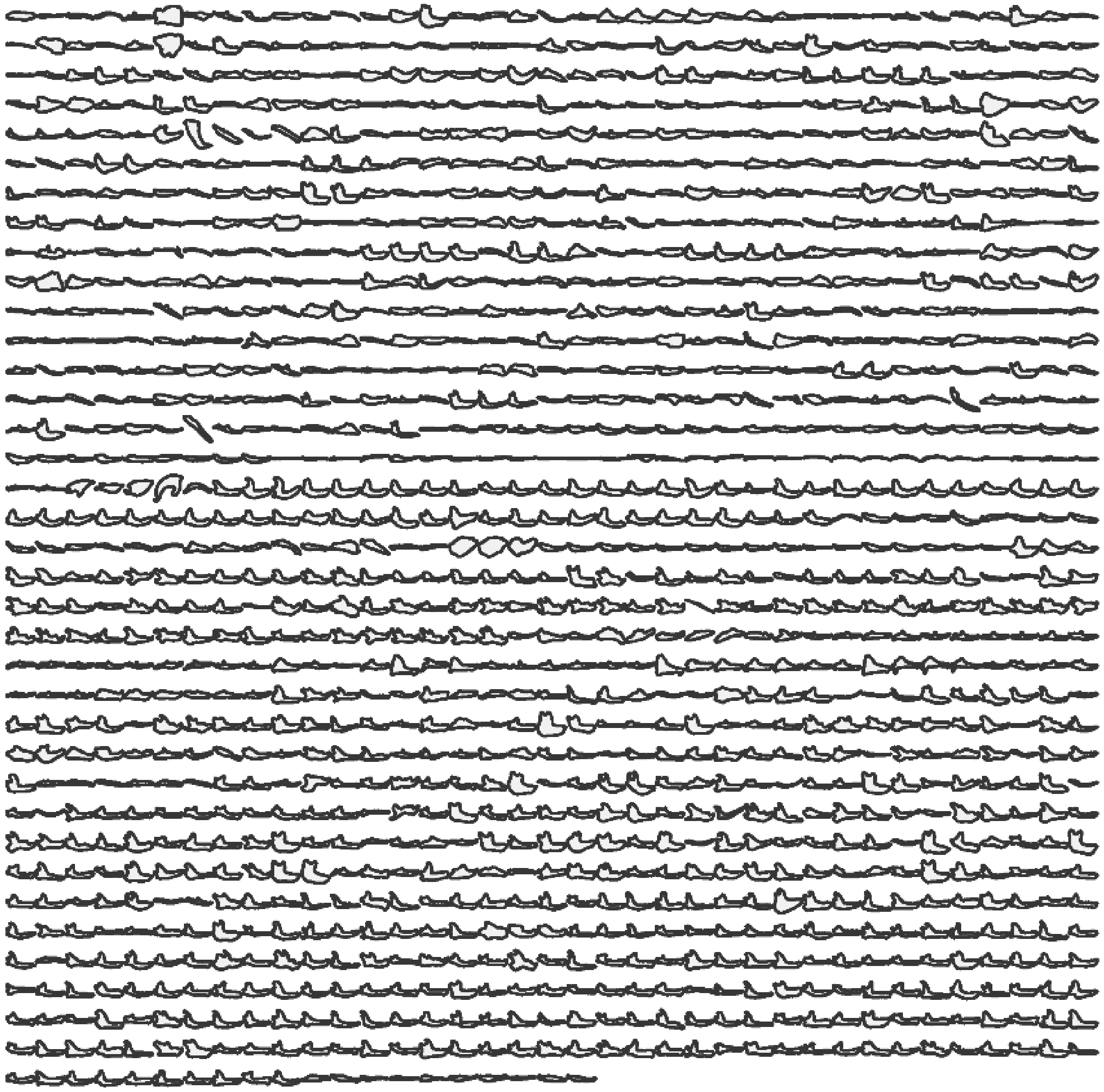

**Fig. S1.**

Panel showing the ~1,350 vertebrate jaw shape outlines included in the shape and biomechanical analyses.

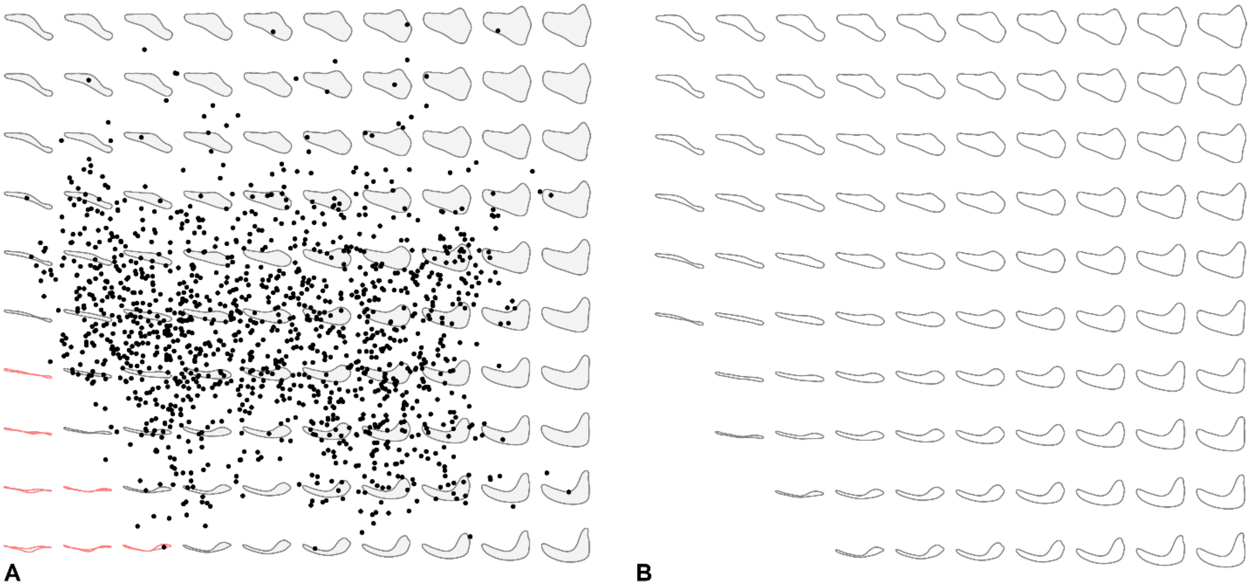

**Fig. S2.**

Jaw shape morphospace. (**A**) Mandible shape morphospace of PC1 (38.9% of variance) and PC2 (20.2% of variance) values from a PCA of elliptic Fourier transform data of >1,000 genera of jawed vertebrates. Warped theoretical jaw shape models used for biomechanical trait measurements and analyses are plotted in the background. (**B**) The 93 warped theoretical jaw models used in experimental and simulation analysis of biomechanical characteristics. Note the removal of several warped models in the lower left corner of the line up, indicating regions in the jaw shape morphospace representing impossible jaw shapes.

| | | | | | | | | | |
|---|---|---|---|---|---|---|---|---|---|
| 1.82 | 1.60 | 1.42 | 1.42 | 1.36 | 1.48 | 1.44 | 1.04 | 1.18 | 0.92 |
| 2.66 | 2.26 | 1.62 | 1.36 | 1.40 | 1.38 | 1.34 | 1.10 | 1.14 | 1.24 |
| 4.28 | 3.08 | 1.86 | 1.54 | 1.38 | 1.20 | 1.44 | 1.06 | 1.08 | 1.20 |
| FLEXIBLE | 3.90 | 2.20 | 1.84 | 1.28 | 1.18 | 1.26 | 1.24 | 1.24 | 1.26 |
| FLEXIBLE | 4.44 | 3.58 | 2.18 | 1.60 | 1.24 | 1.34 | 1.28 | 1.16 | 1.66 |
| FLEXIBLE | 3.64 | 3.42 | 2.96 | 1.54 | 1.28 | 1.54 | 1.44 | 1.54 | 1.48 |
| IMPOSSIBLE | 5.20 | 3.98 | 3.72 | 1.92 | 1.52 | 1.44 | 1.48 | 1.38 | 1.66 |
| IMPOSSIBLE | FLEXIBLE | 5.98 | 4.02 | 2.46 | 1.86 | 1.66 | 1.54 | 1.70 | 1.80 |
| IMPOSSIBLE | IMPOSSIBLE | FLEXIBLE | 8.78 | 4.24 | 2.48 | 2.04 | 1.98 | 1.92 | 2.32 |
| IMPOSSIBLE | IMPOSSIBLE | IMPOSSIBLE | FLEXIBLE | FLEXIBLE | 3.60 | 2.23 | 2.18 | 2.00 | 1.86 |

**A**

| | | | | | | | | | |
|---|---|---|---|---|---|---|---|---|---|
| 83.72 | 75.29 | 71.44 | 71.00 | 75.25 | 80.96 | 79.65 | 55.90 | 65.35 | 51.85 |
| 133.60 | 113.50 | 86.85 | 71.17 | 66.65 | 78.20 | 75.04 | 59.23 | 63.58 | 65.99 |
| 176.20 | 162.70 | 99.51 | 81.53 | 77.28 | 64.29 | 83.25 | 60.07 | 61.06 | 70.80 |
| FLEXIBLE | 199.70 | 119.20 | 95.68 | 68.69 | 63.54 | 69.72 | 69.53 | 66.87 | 69.30 |
| FLEXIBLE | 153.00 | 215.30 | 126.30 | 89.33 | 69.97 | 76.38 | 70.40 | 64.25 | 90.86 |
| FLEXIBLE | 81.42 | 159.10 | 171.40 | 83.79 | 67.84 | 81.35 | 76.95 | 83.79 | 78.35 |
| IMPOSSIBLE | 87.81 | 158.20 | 183.10 | 100.10 | 79.58 | 80.55 | 84.86 | 77.62 | 90.84 |
| IMPOSSIBLE | FLEXIBLE | 191.90 | 161.80 | 134.80 | 99.05 | 89.55 | 81.53 | 92.61 | 94.05 |
| IMPOSSIBLE | IMPOSSIBLE | FLEXIBLE | 337.90 | 245.20 | 131.20 | 116.70 | 99.91 | 101.00 | 135.50 |
| IMPOSSIBLE | IMPOSSIBLE | IMPOSSIBLE | FLEXIBLE | FLEXIBLE | 195.60 | 123.40 | 115.40 | 107.70 | 105.40 |

**B**

**Fig. S3.**

Experimentally derived biomechanical traits for warped theoretical jaw models covering the PC1-PC2 morphospace. (**A**) Maximum displacement values (in mm) and relative magnitudes indicated by heatmap. (**B**) strain energy (N-mm) and relative magnitudes indicated by heatmap. Models indicated "flexible" deflected without fracture, and thus did not have a maximum displacement reference point for calculating strain energy. Models indicated "impossible" are biologically impossible given the crossover of the upper and lower edges of the jaw shape.

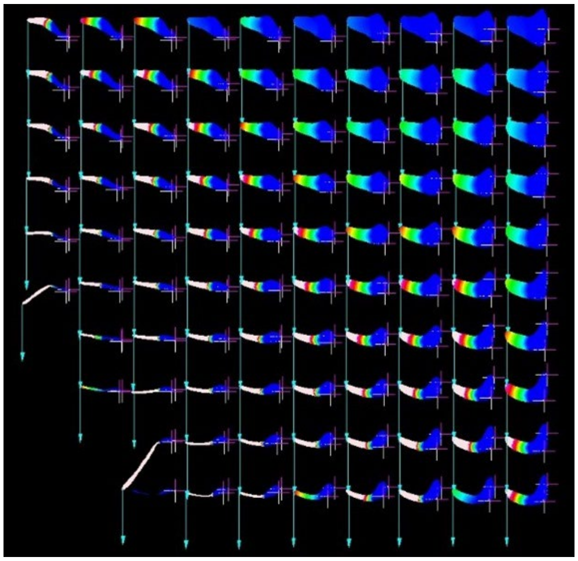

**Fig. S4.**

Outputs of cantilever bending finite element simulations conducted on the extruded 2D jaw models representing the vertebrate jaw shape morphospace. Heatmaps indicate the amount of vertical (dorsoventral) displacement (in mm) experienced by different regions of each warped theoretical jaw shape. The blue vector indicates the direction of ventrally directed bending force placed on the anterior-most node in each model. Cross-hair symbols indicate the location of nodal constraints placed on each model to prevent free body movement.

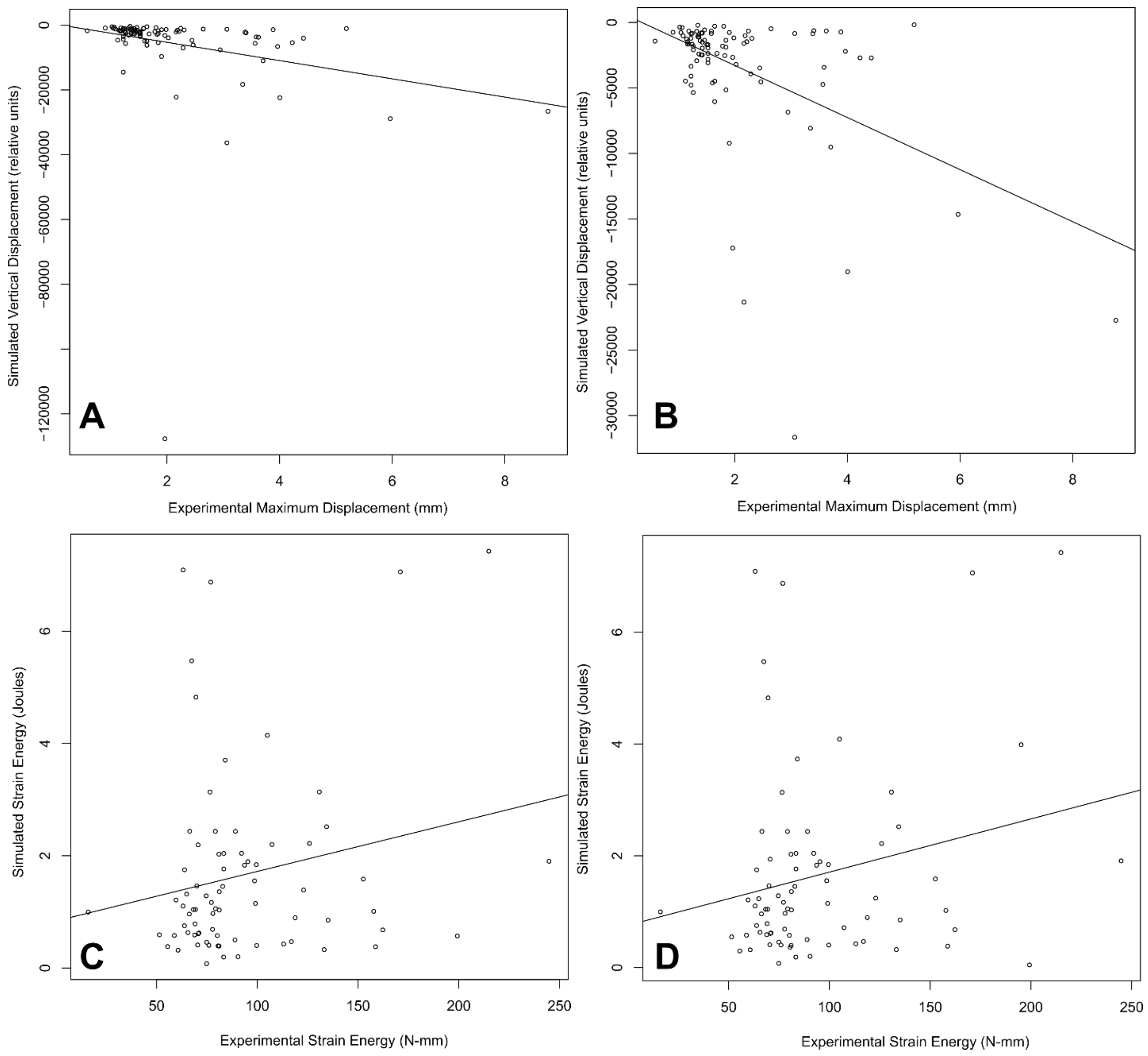


**Fig. S5.**

Linear regression analysis and bivariate plot between experiment and simulation generated biomechanical trait values. Experimentally derived maximum displacement values (x axis) and finite element simulation-based displacement values using models with (**A**) built-in intramandibular joint boundaries and (**B**) those without. Experimentally measured strain energy values versus simulated strain energy values in (**C**) models with intramandibular joints and (**D**) models without intramandibular joints.

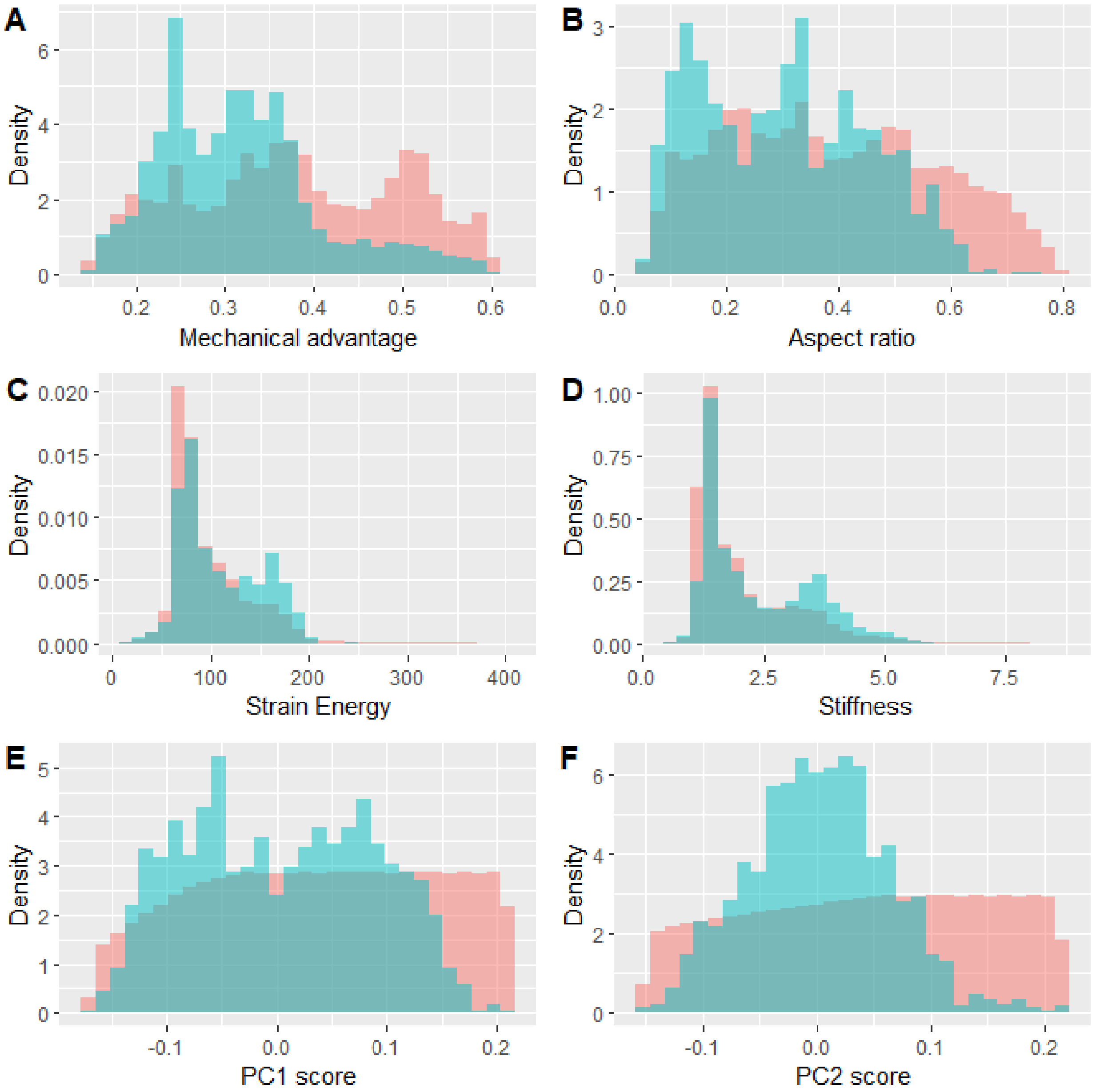


**Fig. S6.**

Comparison between distributions of bootstrapped samples of biomechanical and morphological variables and those derived from actual sampled vertebrate taxa. (**A**) Distribution of randomly sampled MA values (pink) versus MA values from taxa (green), (**B**) AR values, (**C**) Strain energy values, (**D**) Stiffness values (as measured by interpolated maximum deflection from cantilever bending experiment data), (**E**) PC1 shape scores, and (**F**) PC2 shape scores.

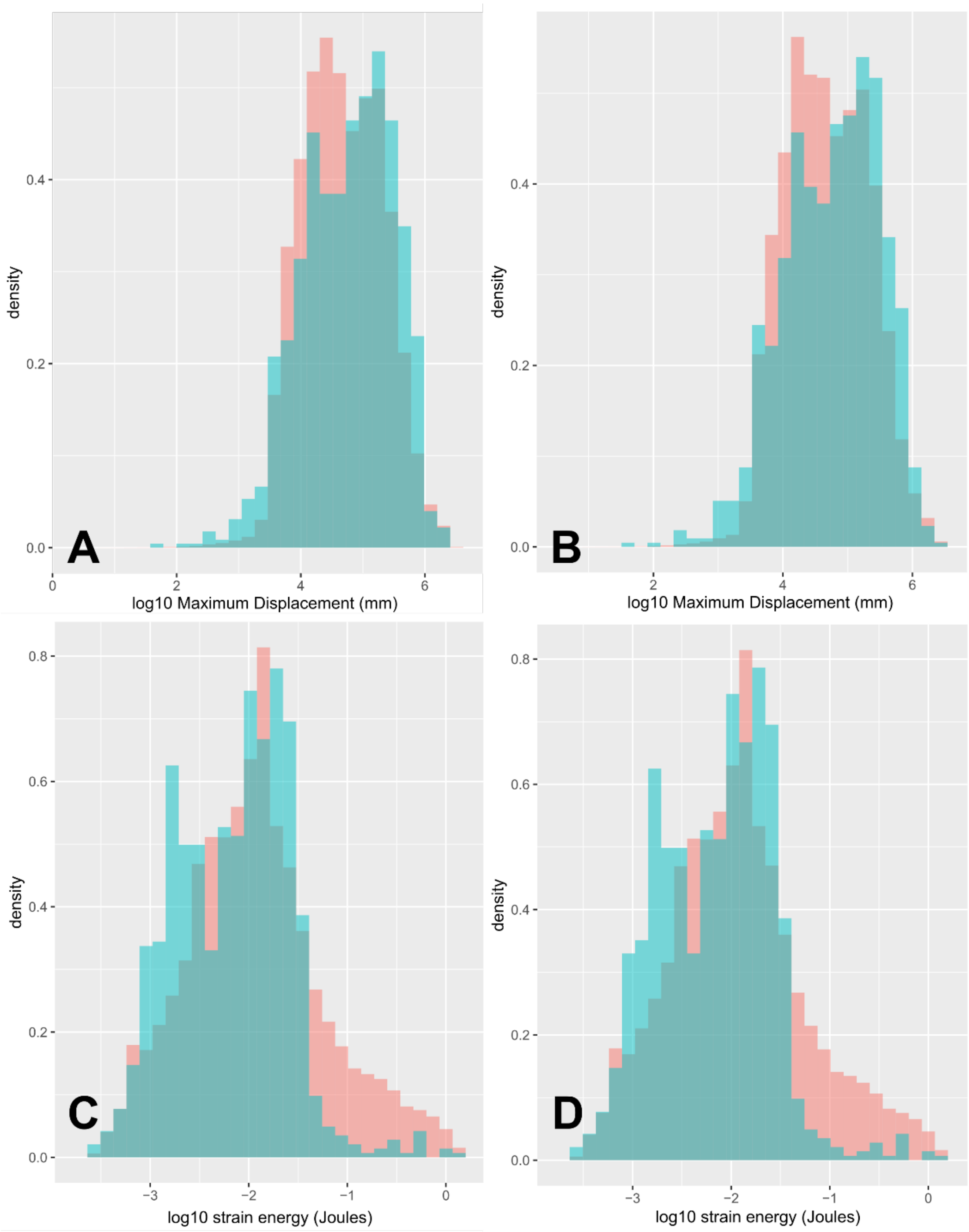


**Fig. S7.**
Comparison between distributions of bootstrapped samples of biomechanical and morphological variables and those derived from actual sampled vertebrate taxa. (A) Distribution of randomly sampled log displacement values (pink) versus log displacement values from interpolated taxonomic values based on solid finite element models (green), (B) log displacement values based on finite element models with intramandibular joints, (C) log strain energy values estimated from solid finite element models, and (D) log strain energy values estimated from intramandibular joint models.

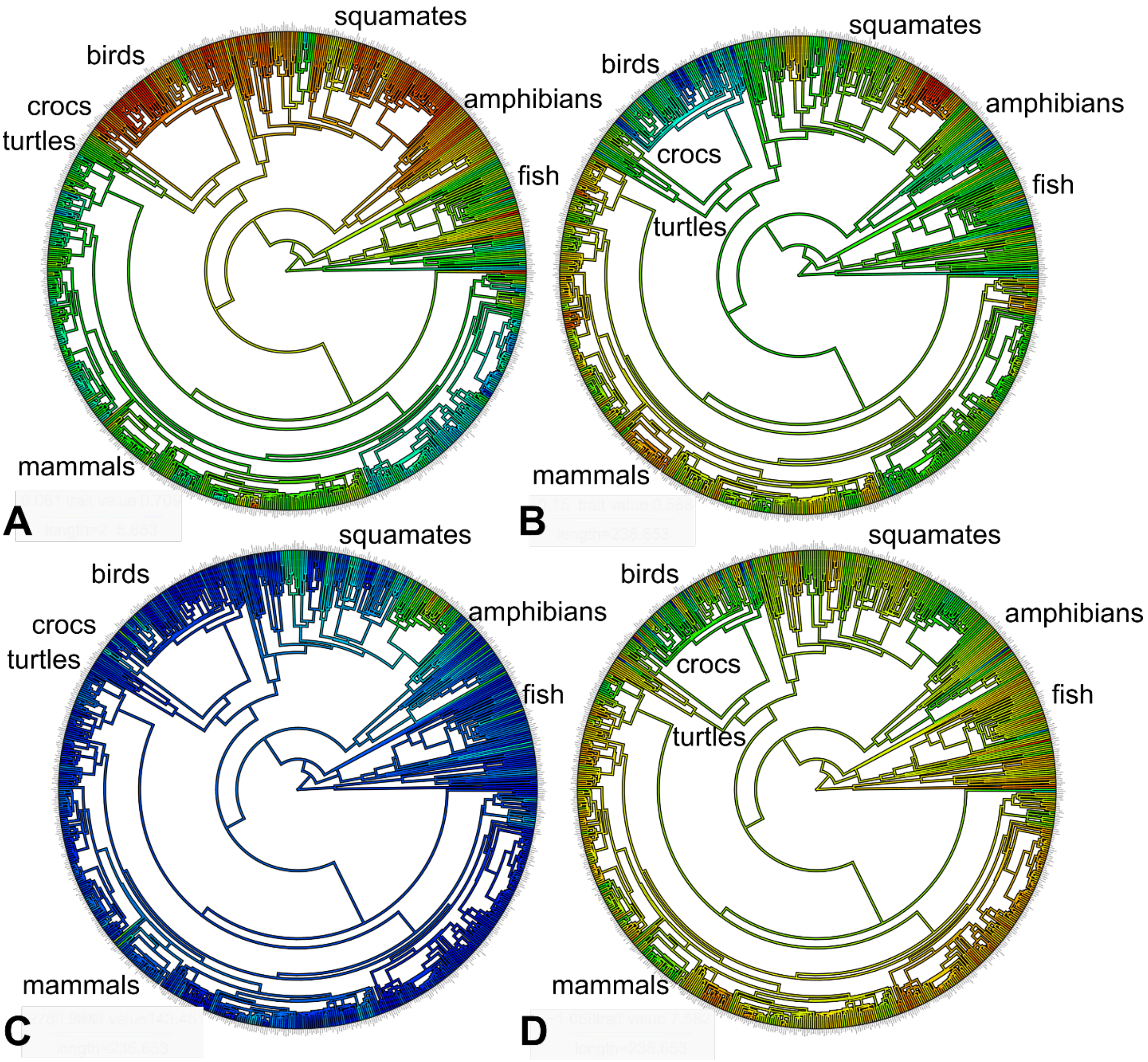


**Fig. S8.**
Mapped biomechanical trait values for extant genera sampled in the jaw functional morphological analyses. (**A**) Aspect ratio (AR, inverse of jaw speed), (**B**) mechanical advantage, (**C**) Maximum displacement values estimated from FEA, (**D**) strain energy values estimated from FEA. Blue/cooler colors indicate larger trait values, red/warmer colors indicate smaller trait values. See Appendix S4 for R script.

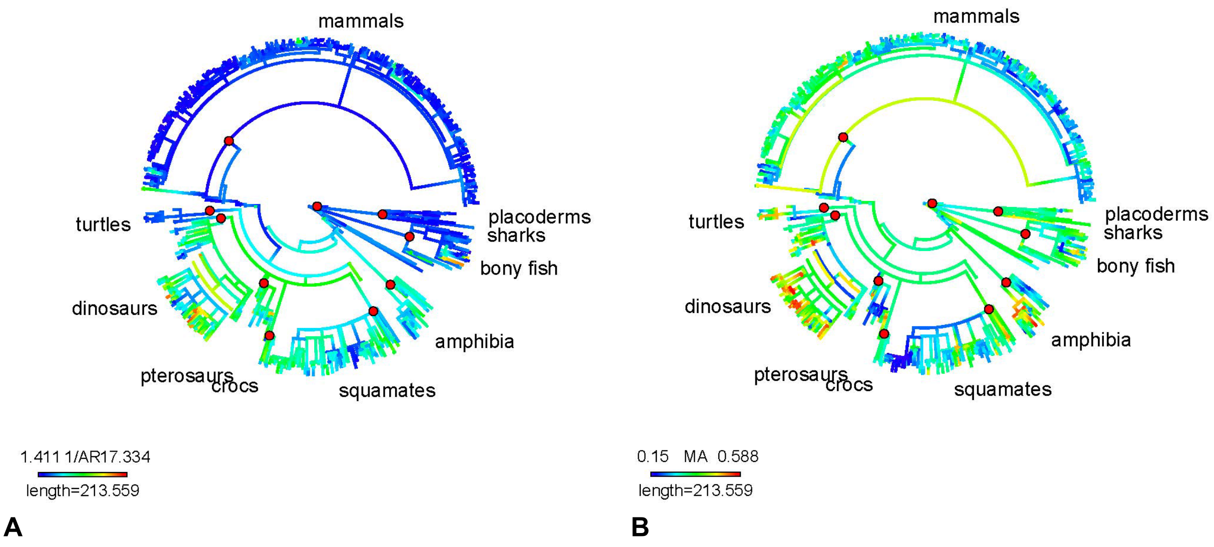


**Fig. S9.**
Jaw shape-derived biomechanical metrics mapped onto the composite phylogeny using ancestral state reconstructions, both extant and fossil taxa. (**A**) Aspect ratio (specifically, 1/AR or "jaw speed"), (**B**) mechanical advantage (MA). See Appendix S4 for R script

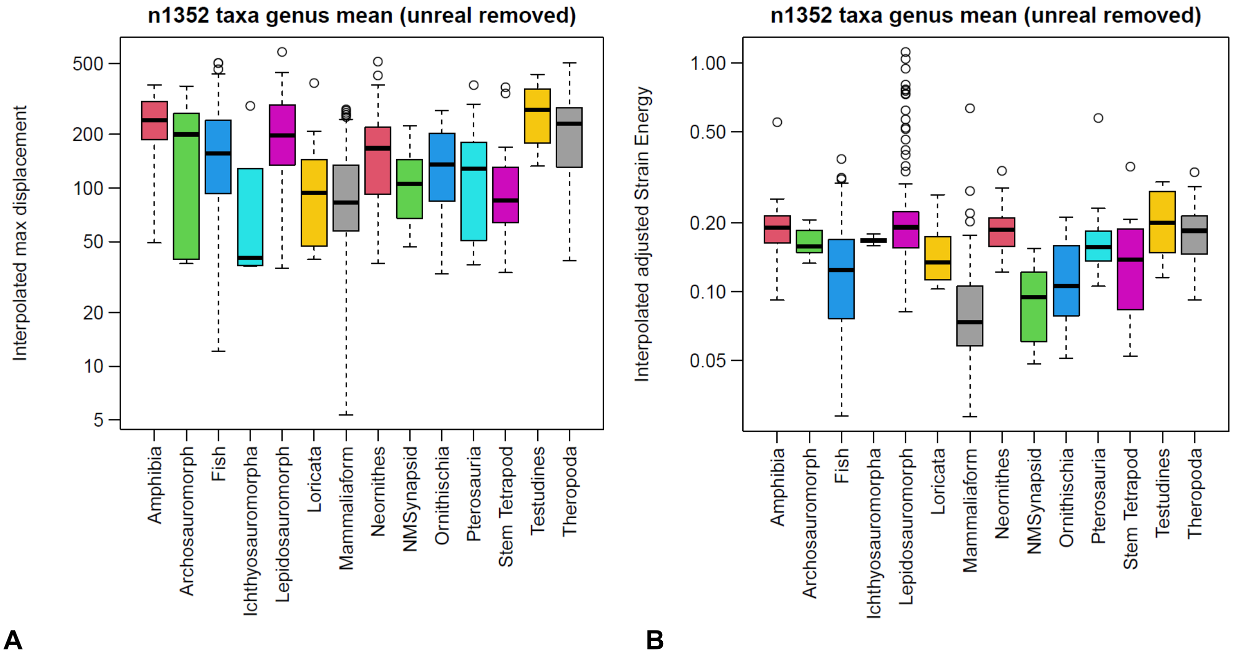


**Fig. S10.**
Box plots of biomechanical trait values estimated using finite element simulations for different vertebrate clades sampled. (**A**) maximum displacement values (in mm), (**B**) strain energy values (in Joules).

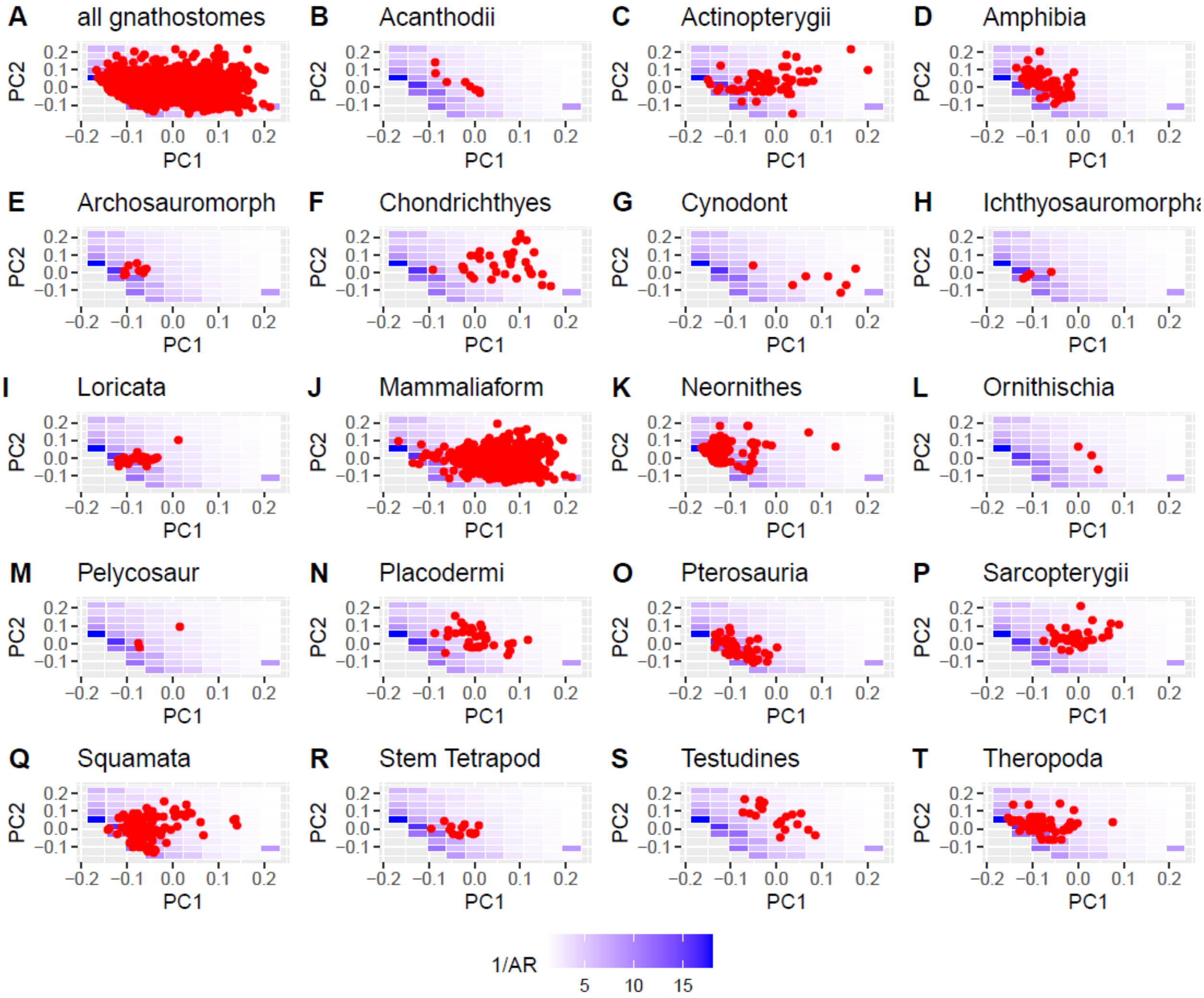


**Fig. S11.**
Location of individual vertebrate clades in the shape morphospace and corresponding estimated values of bite speed (visualized as 1/aspect ratio). Statistical comparisons were made on binned mammal versus non-mammal groups only, given the small sample sizes of most non-mammal clades. (**A**) All specimens sampled, (**B**) Acanthodii, (**C**) Actinopterygii, (**D**) Amphibia, (**E**) Archosauromorpha, (**F**) Chondrichthyes, (**G**) Cynodonts, (**H**) Ichthyomorpha, (**I**) Loricata, (**J**) Mammals, (**K**) Neornithes, (**L**) Ornithischian dinosaurs, (**M**) Pelycosaurs, (**N**) Placodermi, (**O**) Pterosauria, (**P**) Sarcopterygia, (**Q**) Squamata, (**R**) Stem tetrapodomorphs, (**S**) Testudines, (**T**) Theropod dinosaurs.

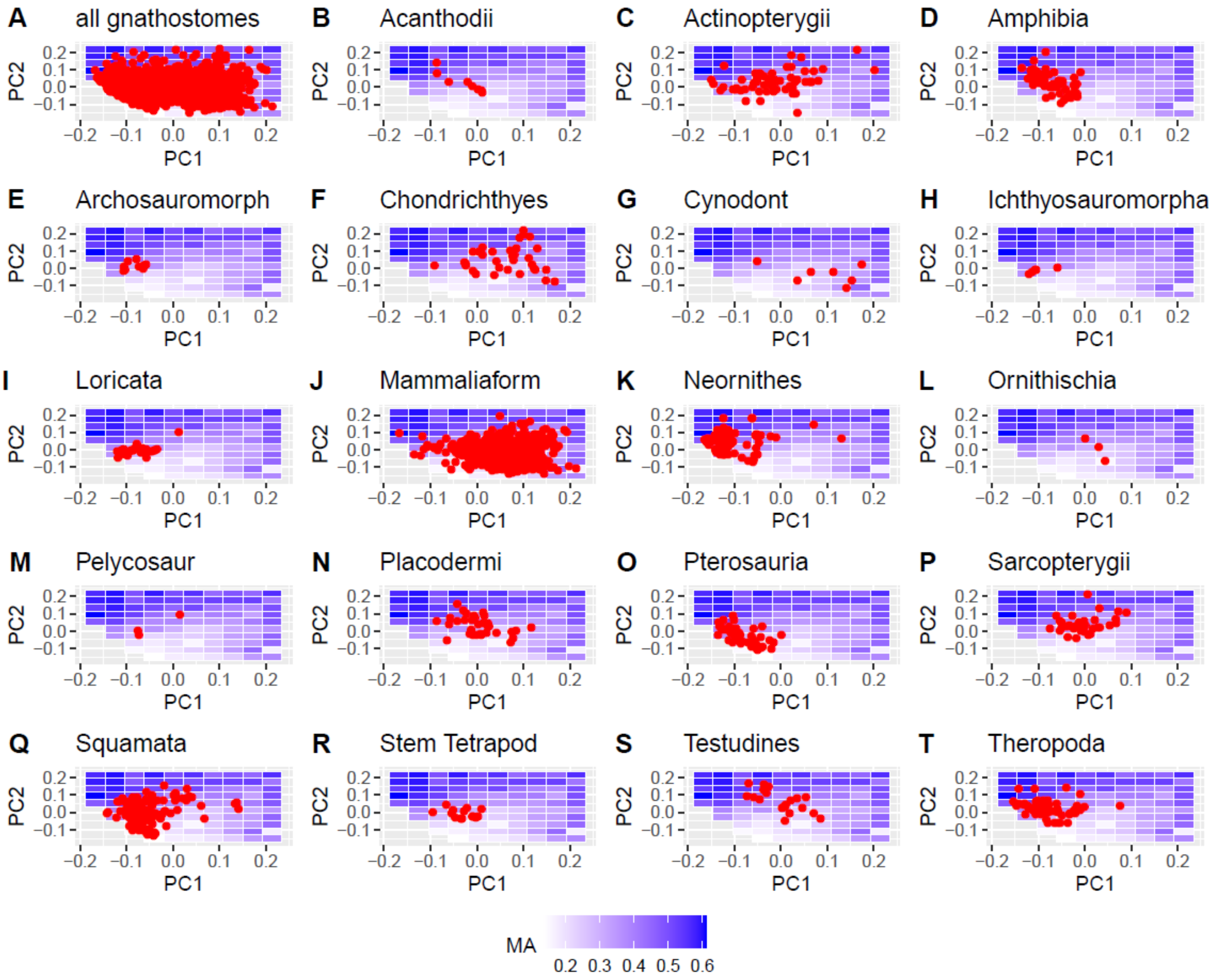


**Fig. S12.**
Location of individual vertebrate clades in the shape morphospace and corresponding estimated values of mechanical advantage (MA). Statistical comparisons were made on binned mammal versus non-mammal groups only, given the small sample sizes of most non-mammal clades. (**A**) All specimens sampled, (**B**) Acanthodii, (**C**) Actinopterygii, (**D**) Amphibia, (**E**) Archosauromorpha, (**F**) Chondrichthyes, (**G**) Cynodonts, (**H**) Ichthyomorpha, (**I**) Loricata, (**J**) Mammals, (**K**) Neornithes, (**L**) Ornithischian dinosaurs, (**M**) Pelycosaurs, (**N**) Placodermi, (**O**) Pterosauria, (**P**) Sarcopterygia, (**Q**) Squamata, (**R**) Stem tetrapodomorphs, (**S**) Testudines, (**T**) Theropod dinosaurs.

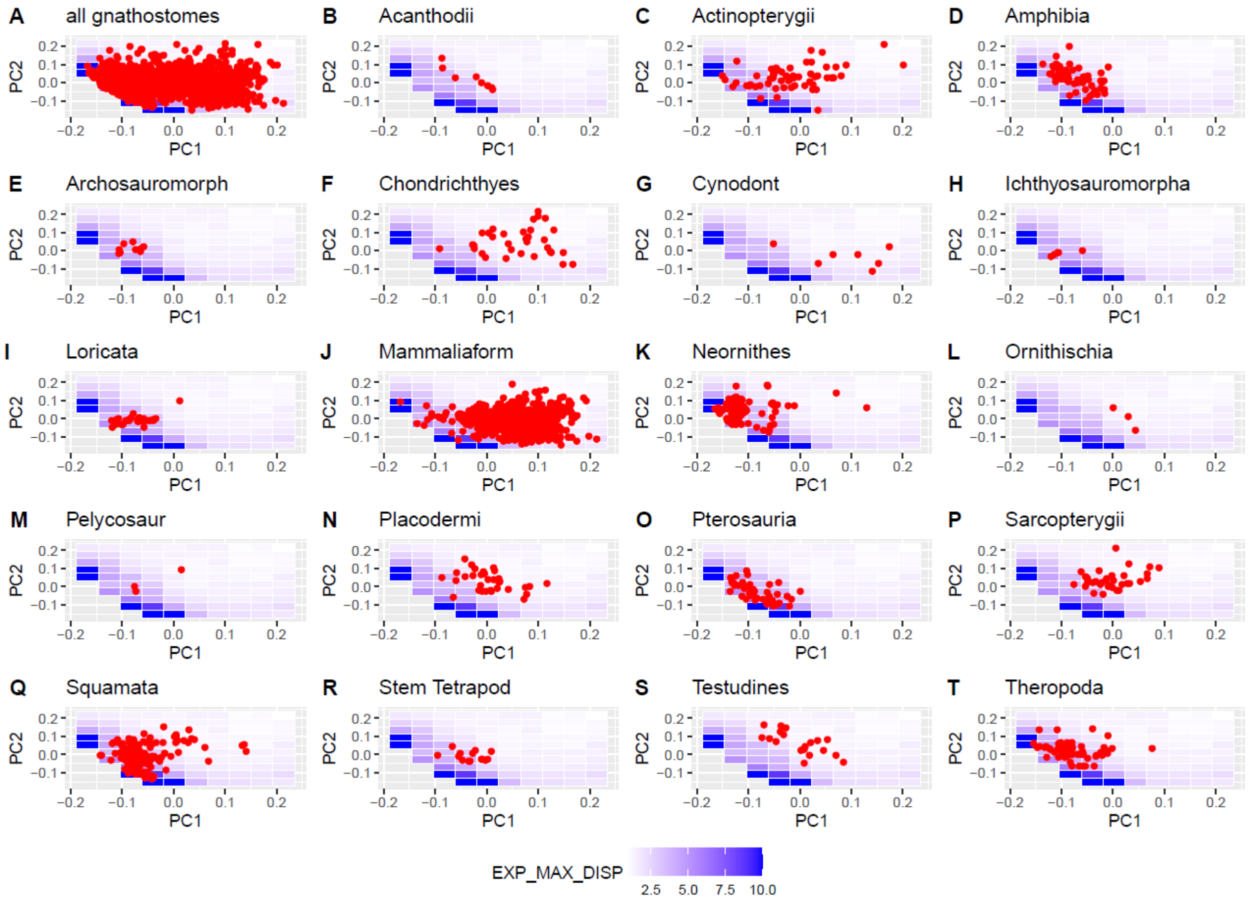


**Fig S13.**
Location of individual vertebrate clades in the shape morphospace and corresponding estimated values of experimental measured maximum displacement values from cantilever bending experiments. Statistical comparisons were made on binned mammal versus non-mammal groups only, given the small sample sizes of most non-mammal clades. (**A**) All specimens sampled, (**B**) Acanthodii, (**C**) Actinopterygii, (**D**) Amphibia, (**E**) Archosauromorpha, (**F**) Chondrichthyes, (**G**) Cynodonts, (**H**) Ichthyomorpha, (**I**) Loricata, (**J**) Mammals, (**K**) Neornithes, (**L**) Ornithischian dinosaurs, (**M**) Pelycosaurs, (**N**) Placodermi, (**O**) Pterosauria, (**P**) Sarcopterygia, (**Q**) Squamata, (**R**) Stem tetrapodomorphs, (**S**) Testudines, (**T**) Theropod dinosaurs.

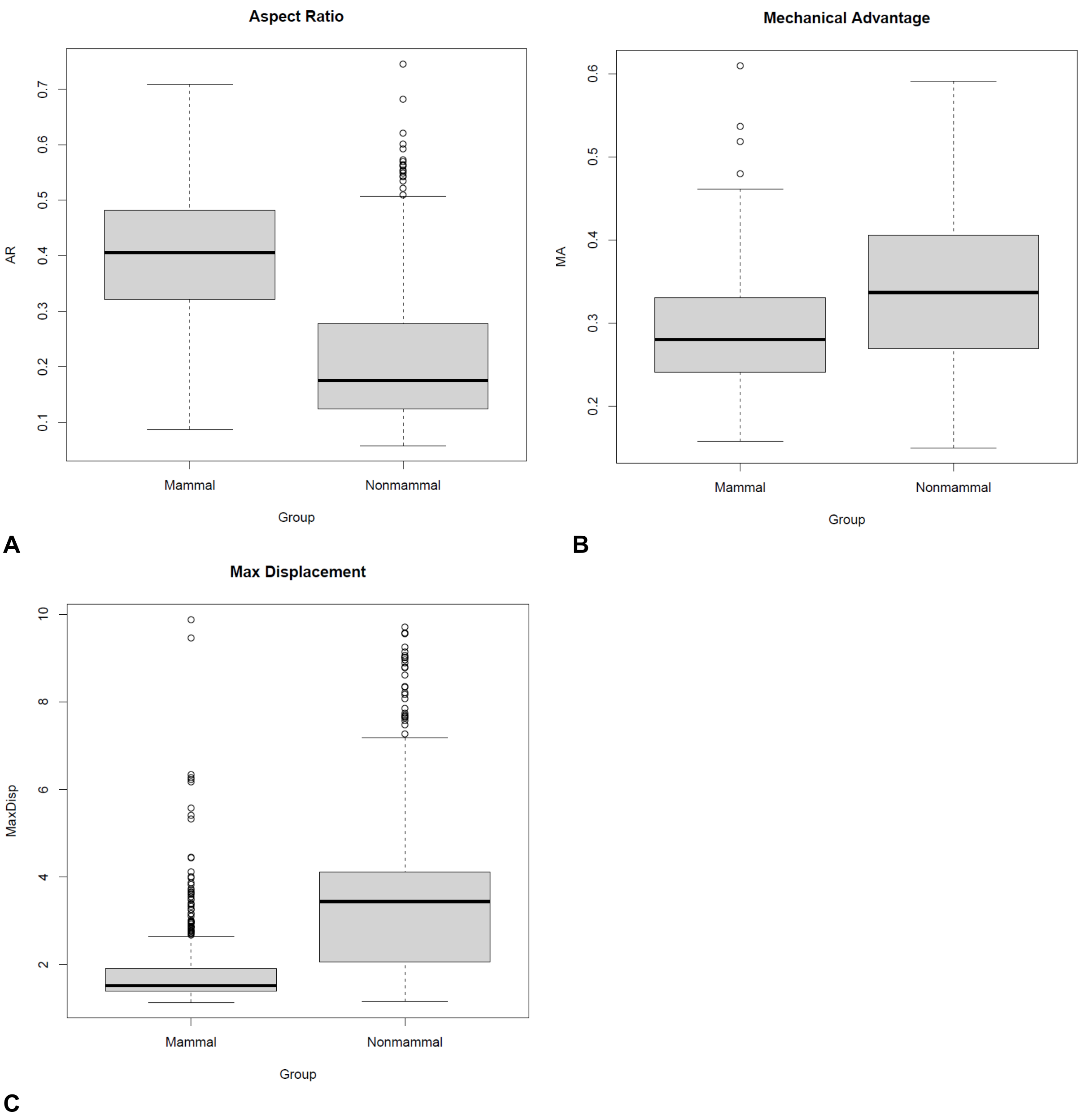


**Fig. S14.**
Box plots comparing biomechanical trait values between mammals and non-mammals in the jaw functional morphology dataset. (**A**) aspect ratio, (**B**) mechanical advantage, (**C**) experimentally measured maximum displacement values. Differences between mammals and non-mammals are significant for all three biomechanical variables based on ANOVA (all $p < 0.0001$) and Wilcoxon rank sum (all $p < 0.0001$) tests.

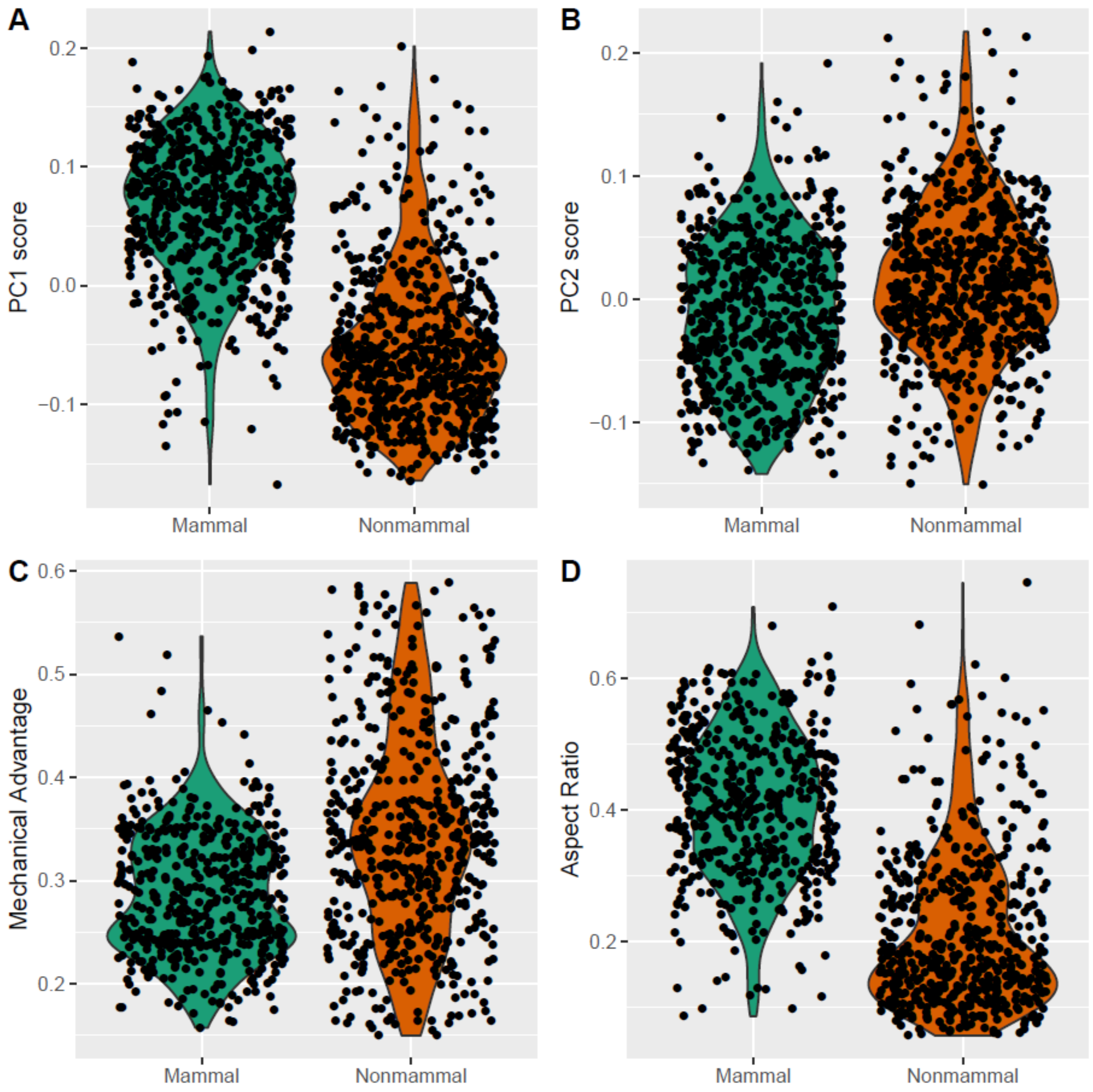


**Fig. S15.**

Violin plots showing the distribution of shape and biomechanical trait values in the mammal and non-mammal partitions of the dataset. (**A**) PC1 jaw shape score, (**B**) PC2 jaw shape score, (**C**) mechanical advantage, (**D**) aspect ratio.

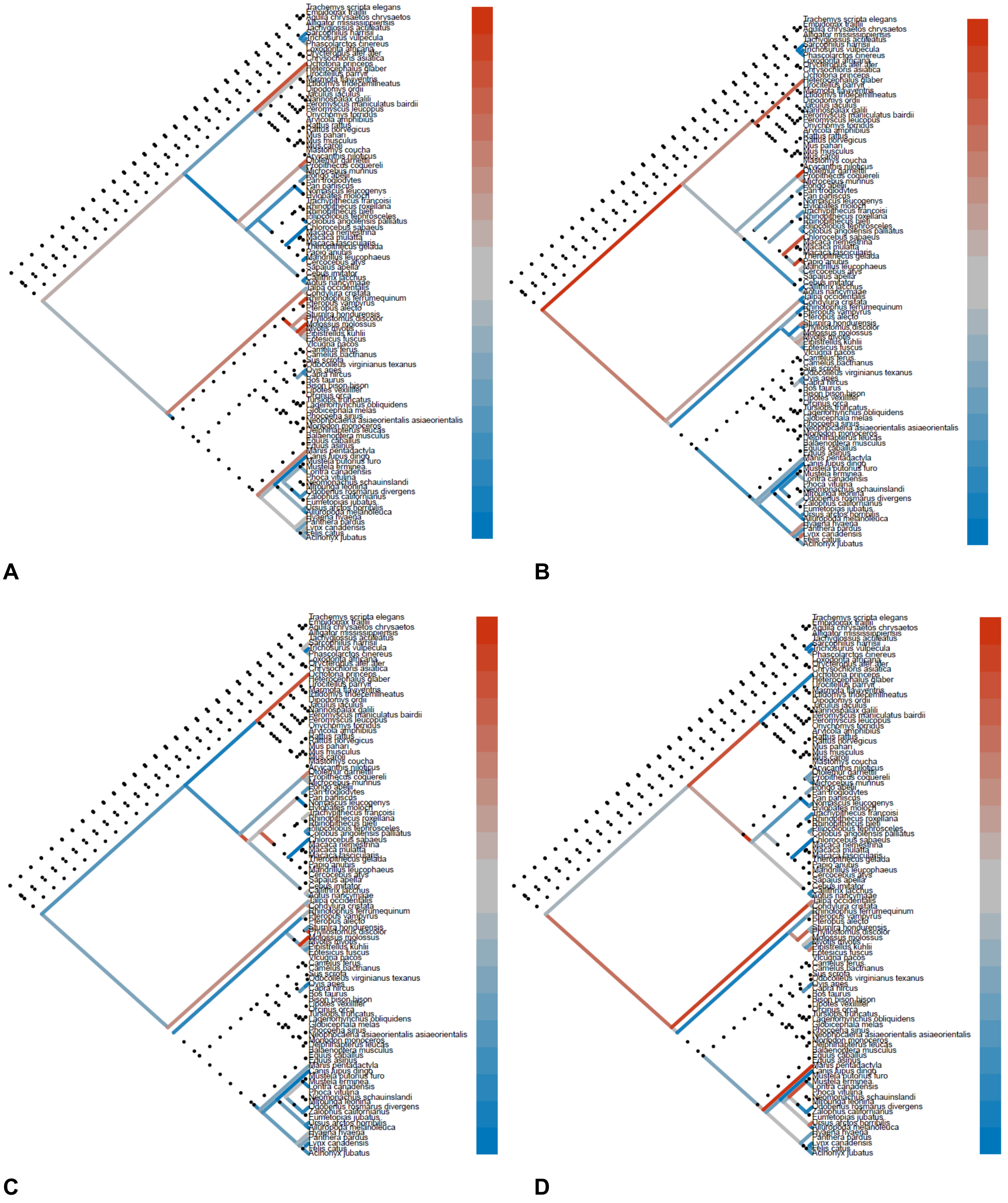


**Fig. S16.**

Heatmapped trees showing different reconstructed evolutionary rates in extant mammal clades in (**A**) *BMP3*, (**B**) *ALX3*, (**C**) *RGS5*, and (**D**) *PLAGL1* nDNA sequences. Dotted lines indicate insufficient data to estimate evolutionary rates.

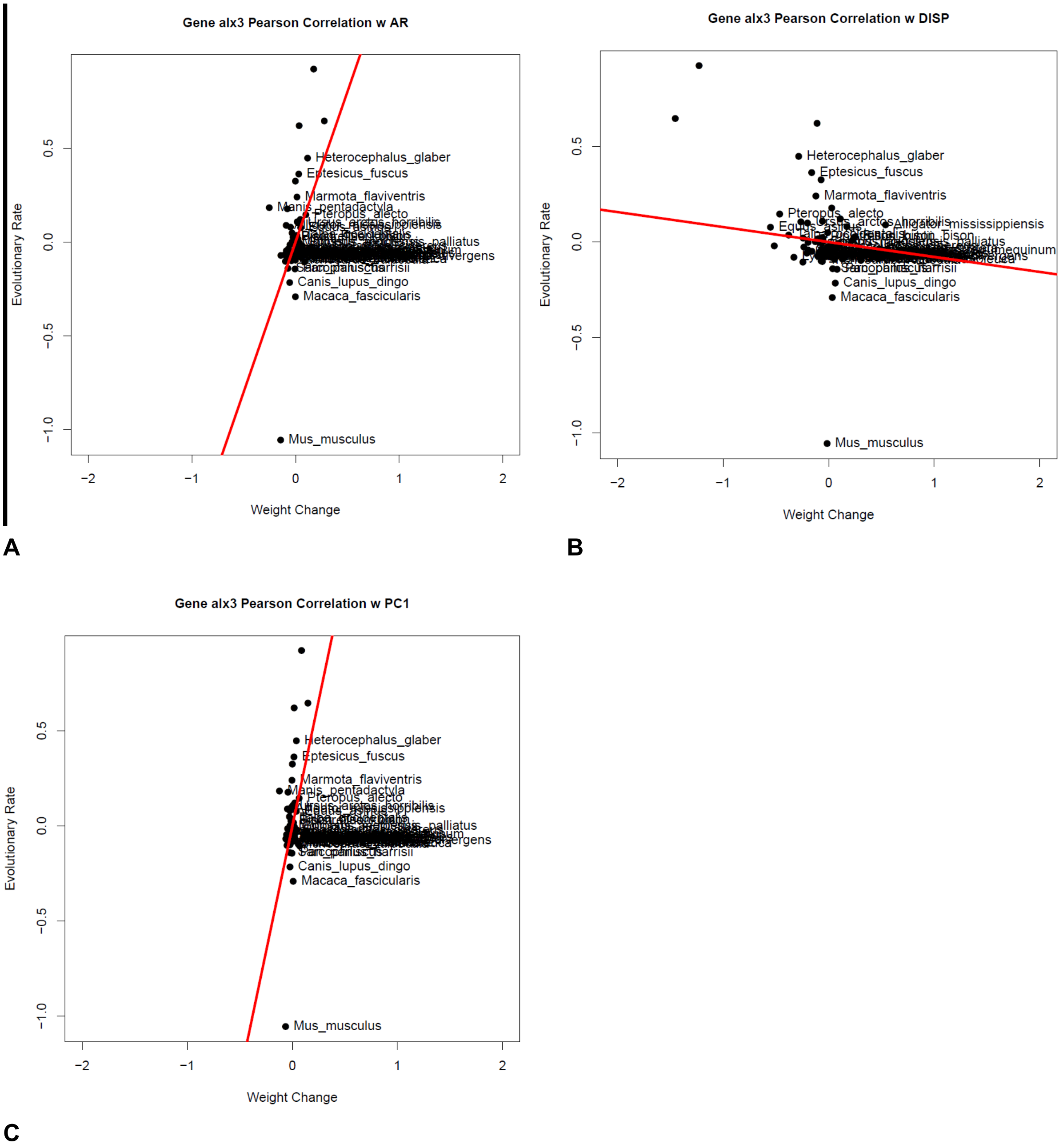


**Fig. S17.**
Relative evolutionary rate correlation for the *ALX3* gene with (**A**) aspect ratio, (**B**) maximum displacement, and (**C**) PC1 jaw shape score. Only statistically significant gene-morphology correlations are shown.

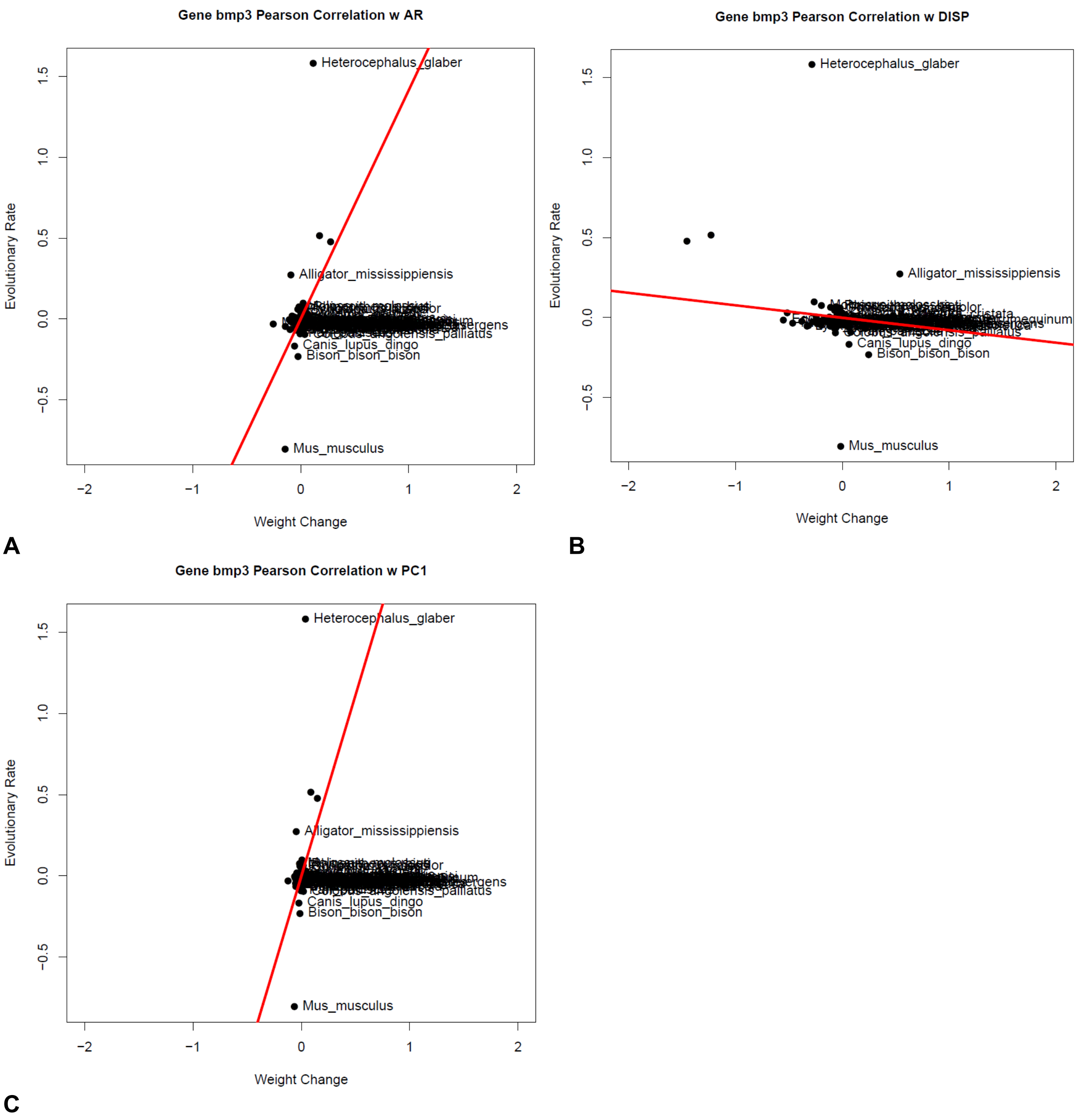


**Fig. S18.**
Relative evolutionary rate correlation for the *BMP3* gene with (**A**) aspect ratio, (**B**) maximum displacement, and (**C**) PC1 jaw shape score. Only statistically significant gene-morphology correlations are shown.

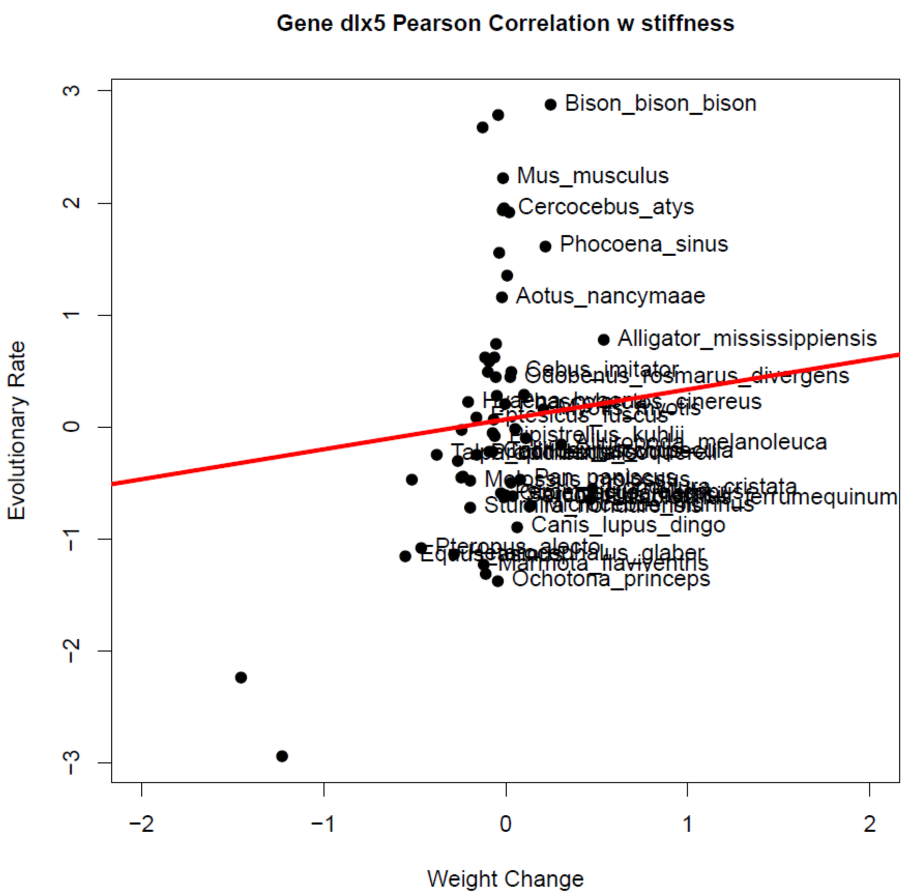


**Fig. S19.**
Relative evolutionary rate correlation for the *DLX5* gene with stiffness (experimental maximum displacement interpolation) values. Only statistically significant gene-morphology correlations are shown.

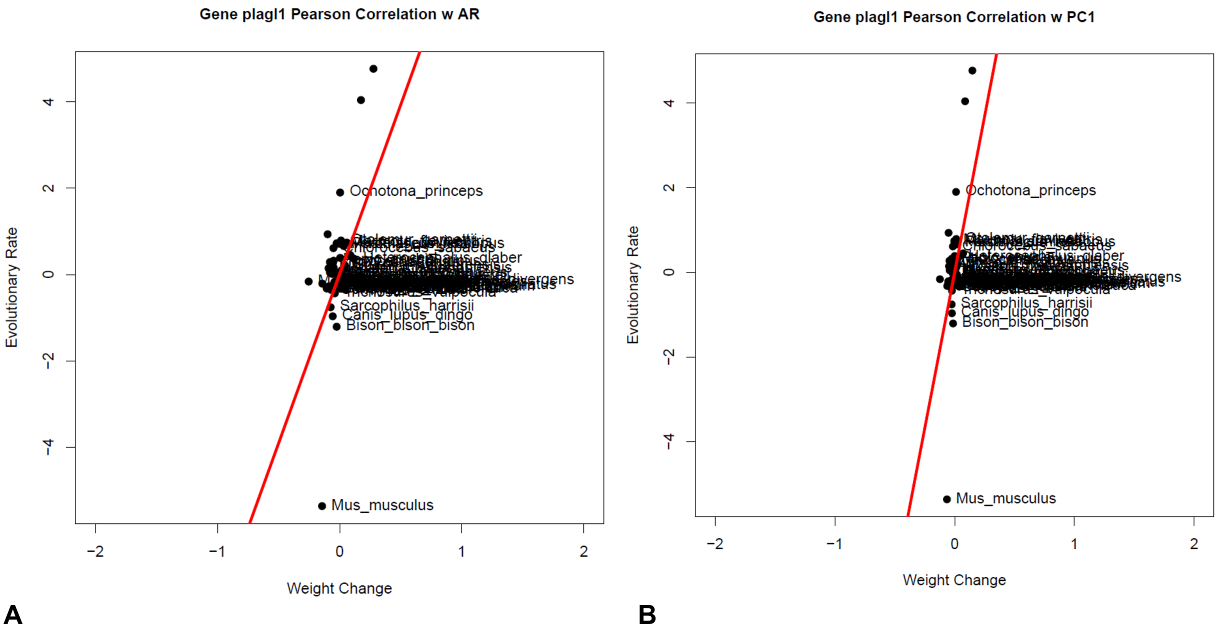


**Fig. S20.**
Relative evolutionary rate correlation for the *PLAGL1* gene with (**A**) aspect ratio, (**B**) PC1 jaw shape score. Only statistically significant gene-morphology correlations are shown.

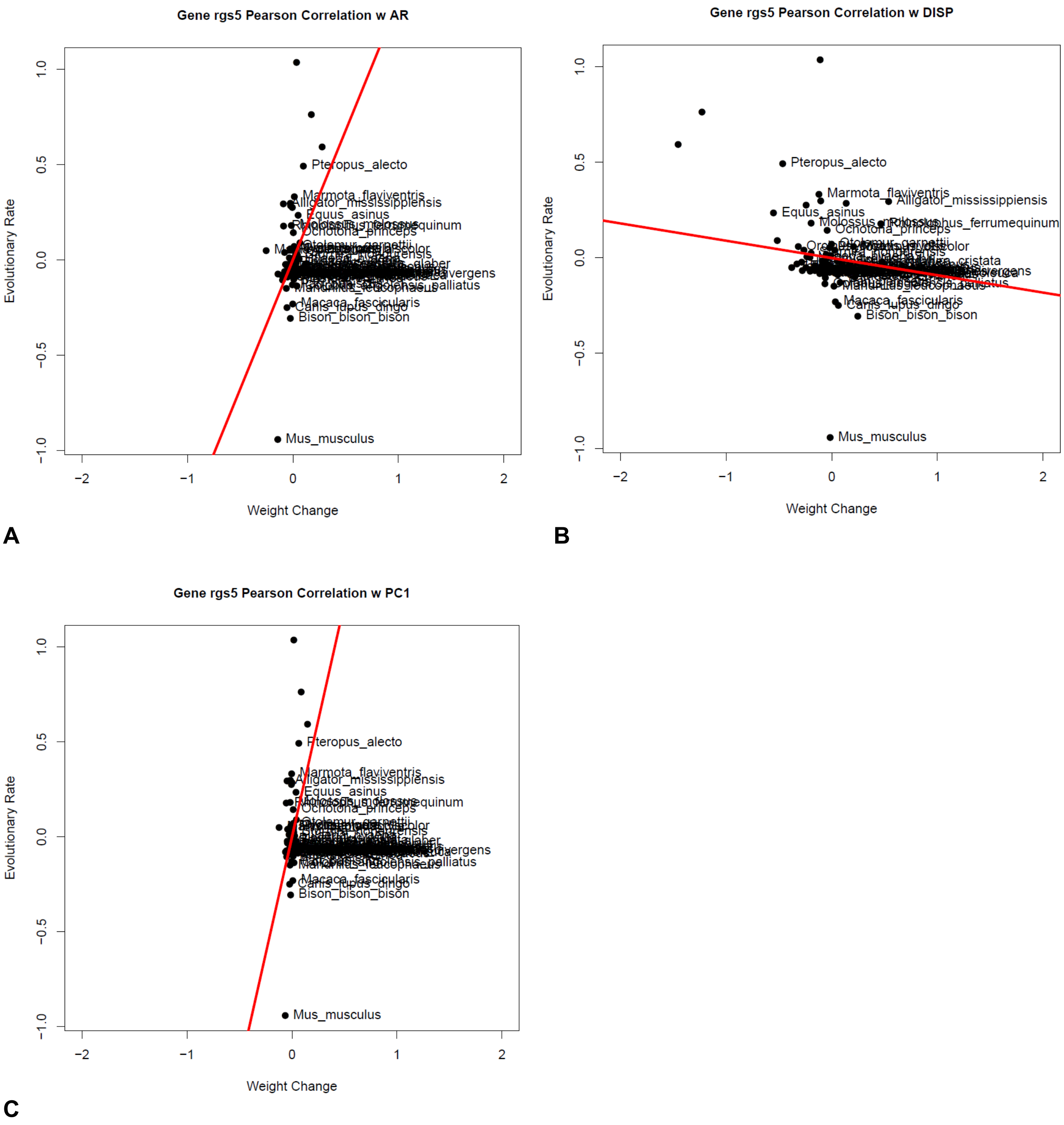


**Fig. S21.**
Relative evolutionary rate correlation for the *RGS5* gene with (**A**) aspect ratio, (**B**) maximum displacement, and (**C**) PC1 jaw shape score. Only statistically significant gene-morphology correlations are shown.

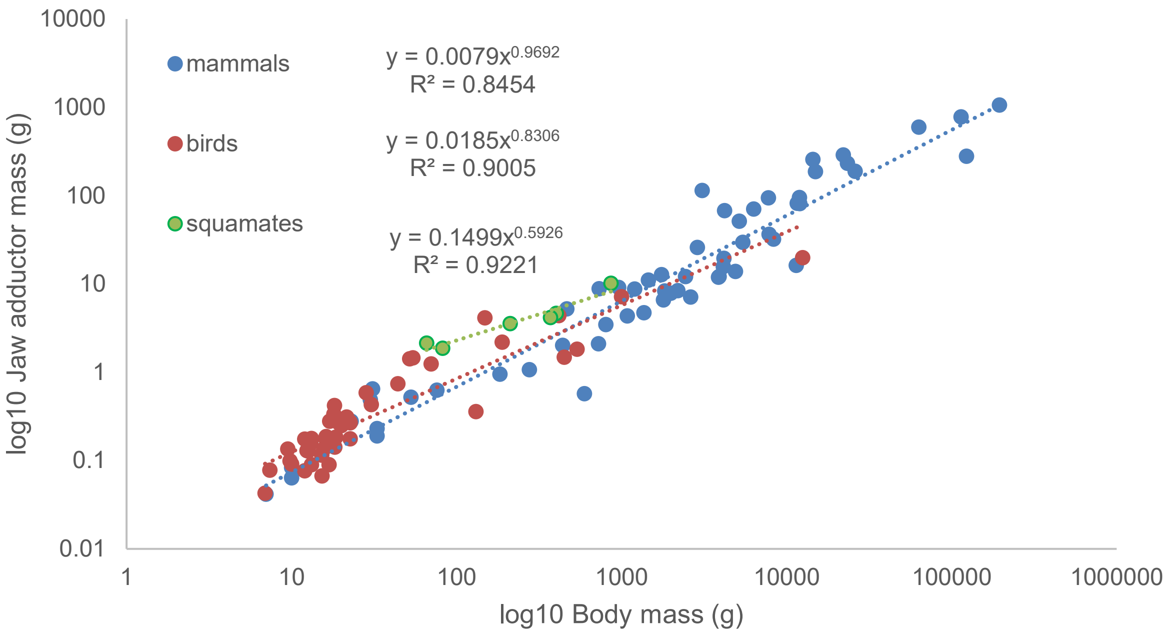


**Fig. S22.**

Regression analysis between log10 body mass (grams) and log10 jaw adductor mass (grams) for a subset of mammals, birds, and squamates sampled in the overall dataset. Regression equations are shown next to the legend for each clade.

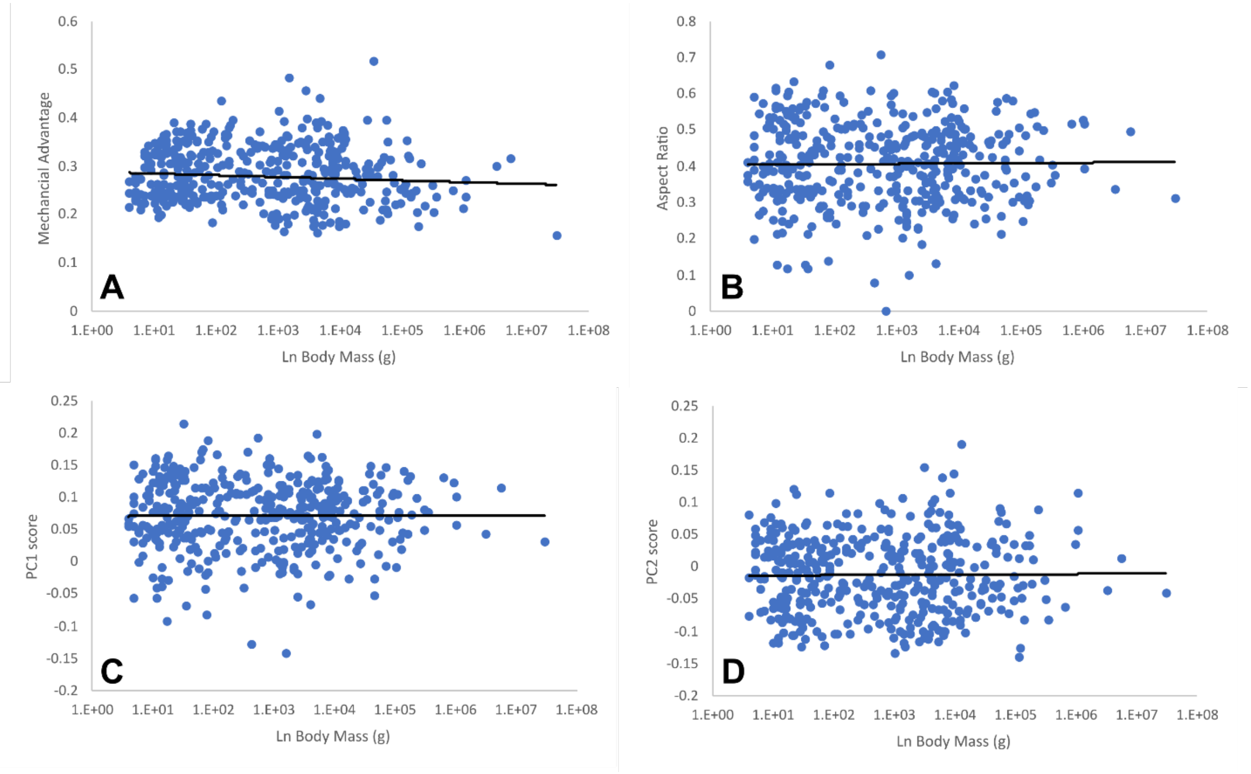


**Fig. S23.**

Scatterplots of log10 body mass versus biomechanical and shape trait values for extant mammal genera. (**A**) mechanical advantage, (**B**) aspect ratio, (**C**) PC1 jaw shape score, (**D**) PC2 jaw shape score. Body mass data are genus adult average values extracted from the PanTHERIA database.

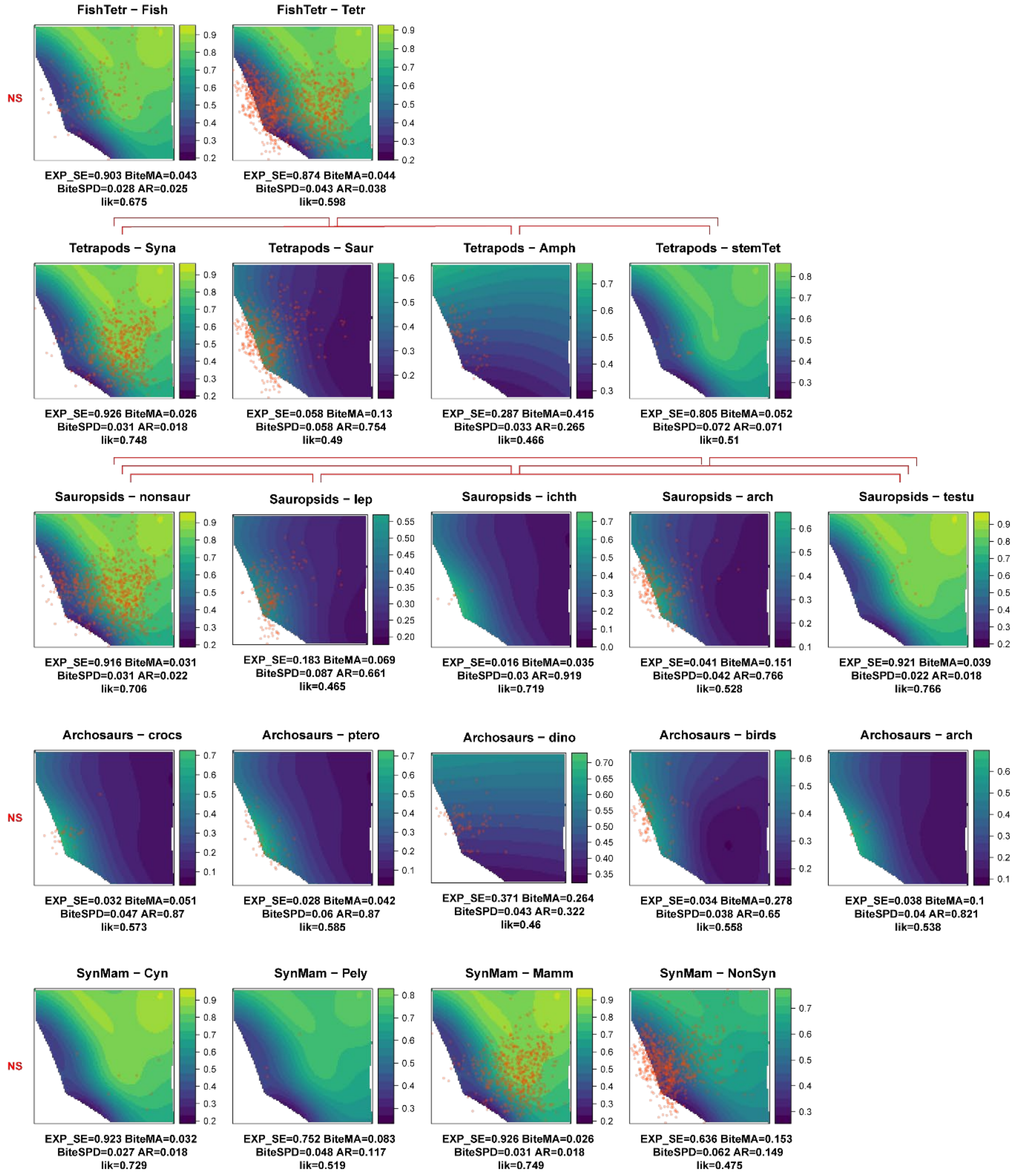


### Fig. S24.

Adaptive landscape regimes over the shape morphospace, showing pairwise comparisons between major vertebrate clades with adequate sample sizes. Statistically significant landscape differences are indicated by brackets over each pair of analyses. EXP_SE, experimentally derived strain energy values; BiteMA, mechanical advantage (in-leve/out-lever ratio); BiteSPD, jaw length/jaw depth ratio; AR, aspect ratio (jaw depth/jaw length); like, likelihood.

| Clade | Genus | Species |
|---|---|---|
| Acanthodii | *Cambaracanthus* | *goodhopensis* |
| Acanthodii | *Cheiracanthus* | *sp* |
| Acanthodii | *Diplacanthus* | *sp* |
| Acanthodii | *Howittacanthus* | *kentoni* |
| Acanthodii | *Ischnacanthus* | *kingi* |
| Acanthodii | *Rockycampacanthus* | *milesi* |
| Acanthodii | *Taemasacanthus* | *erroli* |
| Acanthodii | *Taemasacanthus* | *porca* |
| Actinopterygii | *Acipenser* | *sturio* |
| Actinopterygii | *Alepisaurus* | *ferox* |
| Actinopterygii | *Alopias* | *vulpinus* |
| Actinopterygii | *Amiidae* | *calva* |
| Actinopterygii | *Anarhichas* | *lupus* |
| Actinopterygii | *Anarhichas* | *denticulatus* |
| Actinopterygii | *Anguilla* | *rostrata* |
| Actinopterygii | *Aphanopus* | *carbo* |
| Actinopterygii | *Arapaima* | *gigas* |
| Actinopterygii | *Ariopsis* | *assimilis* |
| Actinopterygii | *Arothron* | *meleagris* |
| Actinopterygii | *Atractosteus* | *spatula* |
| Actinopterygii | *Atractosteus* | *spatula* |
| Actinopterygii | *Atractosteus* | *tristoechus* |
| Actinopterygii | *Atractosteus* | *tropicus* |
| Actinopterygii | *Callionymus* | *lyra* |
| Actinopterygii | *Cnidoglanis* | *megastomus* |
| Actinopterygii | *Cynoscion* | *regalis* |
| Actinopterygii | *Cyprinus* | *carpio* |
| Actinopterygii | *Dapedium* | *politum* |
| Actinopterygii | *Dicentrarchus* | *labrax* |
| Actinopterygii | *Diodon* | *hystrix* |
| Actinopterygii | *Donnrosenia* | *schaefferi* |
| Actinopterygii | *Esox* | *lucius* |
| Actinopterygii | *Eurypharynx* | *pelecanoides* |
| Actinopterygii | *Forcipiger* | *longirostris* |
| Actinopterygii | *Gadus* | *morhua* |
| Actinopterygii | *Gaidropsarus* | *vulgaris* |
| Actinopterygii | *Gnathonemus* | *petersii* |
| Actinopterygii | *Gogosardina* | *coatesi* |
| Actinopterygii | *Gonatodus* | *punctatus* |
| Actinopterygii | *Gymnothorax* | *ocellatus* |
| Actinopterygii | *Halosauropsis* | *macrochir* |
| Actinopterygii | *Hyporhamphus* | *ihi* |
| Actinopterygii | *Lepisosteus* | *oculatus* |
| Actinopterygii | *Lepisosteus* | *osseous* |
| Actinopterygii | *Lepisosteus* | *osseus* |
| Actinopterygii | *Lepisosteus* | *platostomus* |
| Actinopterygii | *Lepisosteus* | *platyrhincus* |
| Actinopterygii | *Lophius* | *piscatorius* |
| Actinopterygii | *Malapterurus* | *microstoma* |

| | | |
|---|---|---|
| Actinopterygii | *Micropterus* | *salmoides* |
| Actinopterygii | *Mimia* | *toombsi* |
| Actinopterygii | *Moxostoma* | *macrolepidotum* |
| Actinopterygii | *Moythomasia* | *durgaringa* |
| Actinopterygii | *Myripristis* | *jacobus* |
| Actinopterygii | *Oncorhynchus* | *mykiss* |
| Actinopterygii | *Pleuronectes* | *platessa* |
| Actinopterygii | *Polypterus* | *bichir* |
| Actinopterygii | *Porichthys* | *myriaster* |
| Actinopterygii | *Rhinocephalus* | *planiceps* |
| Actinopterygii | *Salmo* | *sp.* |
| Actinopterygii | *Scarus* | *sp.* |
| Actinopterygii | *Silurid* | *sp* |
| Actinopterygii | *Sphyraena* | *barracuda* |
| Actinopterygii | *Strongylura* | *timucu* |
| Actinopterygii | *Synodus* | *sp* |
| Actinopterygii | *Thunnus* | *albacares* |
| Actinopterygii | *Watsonichthys* | *pectinatus* |
| Actinopterygii | *Xiphias* | *gladius* |
| Amphibia | *Ambystoma* | *gracile* |
| Amphibia | *Ambystoma* | *tigrinum* |
| Amphibia | *Amphiuma* | *means* |
| Amphibia | *Amphiuma* | *pholeter* |
| Amphibia | *Aneides* | *lugubris* |
| Amphibia | *Atretochoana* | *eiselti* |
| Amphibia | *Bombina* | *orientalis* |
| Amphibia | *Boulengerula* | *taitanus* |
| Amphibia | *Chiropterotriton* | *chiropterus* |
| Amphibia | *Chiropterotriton* | *magnipes* |
| Amphibia | *Dermophis* | *mexicanus* |
| Amphibia | *Desmognathus* | *apalachicolae* |
| Amphibia | *Desmognathus* | *auriculatus* |
| Amphibia | *Dicamptodon* | *ensatus* |
| Amphibia | *Eoherpeton* | *watsoni* |
| Amphibia | *Epicrionops* | *bicolor* |
| Amphibia | *Eurycea* | *rathbuni* |
| Amphibia | *Eurycea* | *robusta* |
| Amphibia | *Eurycea* | *sosorum* |
| Amphibia | *Eurycea* | *waterlooensis* |
| Amphibia | *Gastrophryne* | *usta* |
| Amphibia | *Geotrypetes* | *seraphini* |
| Amphibia | *Gephyrostegus* | *bohemicus* |
| Amphibia | *Gephyrostegus* | *bohemicus* |
| Amphibia | *Hemiphractus* | *elioti* |
| Amphibia | *Hemiphractus* | *elioti* |

| | | |
|---|---|---|
| Amphibia | *Hemiphractus* | *fasciatus* |
| Amphibia | *Hemiphractus* | *kaylockae* |
| Amphibia | *Hemiphractus* | *panamensis* |
| Amphibia | *Hemiphractus* | *proboscideus* |
| Amphibia | *Herpele* | *squalostoma* |
| Amphibia | *Hydromantes* | *platycephalus* |
| Amphibia | *Ichthyophis* | *bannanicus* |
| Amphibia | *Ichthyophis* | *kohtaoensis* |
| Amphibia | *Karsenia* | *koreana* |
| Amphibia | *Microcaecilia* | *iwokramae* |
| Amphibia | *Necturus* | *maculosus* |
| Amphibia | *Necturus* | *maculosus* |
| Amphibia | *Onychodactylus* | *japonicus* |
| Amphibia | *Paradactylodon* | *persicus* |
| Amphibia | *Phaeognathus* | *hubrichti* |
| Amphibia | *Proterogyrinus* | *scheelei* |
| Amphibia | *Rhinatrema* | *bivittatum* |
| Amphibia | *Rhyacotriton* | *variegatus* |
| Amphibia | *Schistometopum* | *thomense* |
| Amphibia | *Scolecomorphus* | *sp* |
| Amphibia | *Silvanerpeton* | *miripedes* |
| Amphibia | *Siphonops* | *annulatus* |
| Amphibia | *Siren* | *lacertina* |
| Amphibia | *Theloderma* | *stellatum* |
| Amphibia | *Thorius* | *adelos* |
| Amphibia | *Thorius* | *aureus* |
| Amphibia | *Thorius* | *longicaudus* |
| Amphibia | *Thorius* | *minutissimus* |
| Amphibia | *Thorius* | *narisovalis* |
| Amphibia | *Thorius* | *pinicola* |
| Amphibia | *Thorius* | *tlaxiacus* |
| Amphibia | *Typhlonectes* | *compressicauda* |
| Amphibia | *Typhlonectes* | *natans* |
| Amphibia | *Typhlonectes* | *natans* |
| Amphibia | *Uraeotyphlus* | *narayani* |
| Amphibia | *Xenopus* | *laevis* |
| Archosauromorph | *Azendohsaurus* | *madagaskarensis* |
| Archosauromorph | *Boreopricea* | *funerea* |
| Archosauromorph | *Chanaresuchus* | *bonapartei* |
| Archosauromorph | *Doswellia* | *sixmilensis* |
| Archosauromorph | *Euparkeria* | *capensis* |
| Archosauromorph | *Garjainia* | *prima* |
| Archosauromorph | *Macrocnemus* | *bassanii* |
| Archosauromorph | *Mesosuchus* | *browni* |
| Archosauromorph | *Proterosuchus* | *fergusi* |

| | | |
|---|---|---|
| Chondrichthyes | *Aetobatus* | *narinari* |
| Chondrichthyes | *Aetobatus* | *narinari* |
| Chondrichthyes | *Aetomylaeus* | *bovinus* |
| Chondrichthyes | *Carcharhinus* | *galapagensis* |
| Chondrichthyes | *Carcharhinus* | *plumbeus* |
| Chondrichthyes | *Carcharias* | *taurus* |
| Chondrichthyes | *Carcharodon* | *sp.* |
| Chondrichthyes | *Cephaloscyllium* | *ventriosum* |
| Chondrichthyes | *Chiloscyllium* | *plagiosum* |
| Chondrichthyes | *Chimaera* | *sp* |
| Chondrichthyes | *Cladodoides* | *wildungensis* |
| Chondrichthyes | *Dasyatis* | *centroura* |
| Chondrichthyes | *Diplodoselache* | *woodi (gen.et sp. Nov.)* |
| Chondrichthyes | *Dipturus* | *laevis* |
| Chondrichthyes | *Elasmodectes* | *(cf.) willetti* |
| Chondrichthyes | *Ginglymostoma* | *cirratum* |
| Chondrichthyes | *Glyphis* | *gangeticus* |
| Chondrichthyes | *Heterodontus* | *francisci* |
| Chondrichthyes | *Hexanchus* | *griseus* |
| Chondrichthyes | *Hydrolagus* | *colliei* |
| Chondrichthyes | *Isistius* | *brasiliensis* |
| Chondrichthyes | *Mustelus* | *canis* |
| Chondrichthyes | *Myliobatis* | *californica* |
| Chondrichthyes | *Myriacanthus* | *paradoxus* |
| Chondrichthyes | *Orectolobus* | *ornatus* |
| Chondrichthyes | *Pastinachus* | *sephen* |
| Chondrichthyes | *Pseudobatos* | *lentiginosus* |
| Chondrichthyes | *Rhinoptera* | *bonasus* |
| Chondrichthyes | *Saivodus* | *magnificus* |
| Chondrichthyes | *Scyliorhinus* | *retifer* |
| Chondrichthyes | *Sphyrna* | *mokarran* |
| Chondrichthyes | *Squalus* | *acanthias* |
| Chondrichthyes | *Squatina* | *aculeata* |
| Cynodont | *Bocatherium* | *sp* |
| Cynodont | *Cynognathus* | *sp* |
| Cynodont | *Dvinia* | *sp* |
| Cynodont | *Exaeretodon* | *sp* |
| Cynodont | *Probainognathus* | *sp* |
| Cynodont | *Procynosuchus* | *sp* |
| Cynodont | *Thrinaxodon* | *sp* |
| Ichthyosauromorpha | *Cartorhynchus* | *lenticarpus* |
| Ichthyosauromorpha | *Chaohusaurus* | *chaoxianensis* |
| Ichthyosauromorpha | *Cymbospondylus* | *piscosus* |
| Ichthyosauromorpha | *Hupehsuchus* | *sp* |
| Ichthyosauromorpha | *Ophthalmosaurus* | *icenicus* |

| | | |
|---|---|---|
| Ichthyosauromorpha | *Temnodontosaurus* | *platyodon* |
| Loricata | *Alligator* | *mississippiensis* |
| Loricata | *Alligator* | *sinensis* |
| Loricata | *Batrachotomus* | *kupferzellensis* |
| Loricata | *Caiman* | *crocodilus* |
| Loricata | *Caiman* | *crocodilus* |
| Loricata | *Caiman* | *latirostris* |
| Loricata | *Caiman* | *yacare* |
| Loricata | *Crocodylus* | *intermedius* |
| Loricata | *Crocodylus* | *johnstoni* |
| Loricata | *Crocodylus* | *moreletii* |
| Loricata | *Crocodylus* | *moreletti* |
| Loricata | *Crocodylus* | *niloticus* |
| Loricata | *Crocodylus* | *novaeguinae* |
| Loricata | *Crocodylus* | *porosus* |
| Loricata | *Crocodylus* | *rhombifer* |
| Loricata | *Gavialis* | *gangeticus* |
| Loricata | *Gavialis* | *gangeticus* |
| Loricata | *Gavialis* | *gangeticus* |
| Loricata | *Laganosuchus* | *thaumastos* |
| Loricata | *Mecistops* | *cataphractus* |
| Loricata | *Mecistops* | *cataphractus* |
| Loricata | *Melanosuchus* | *niger* |
| Loricata | *Osteolaemus* | *tetraspis* |
| Loricata | *Paleosuchus* | *palpebrosus* |
| Loricata | *Paleosuchus* | *trigonatus* |
| Loricata | *Paleosuchus* | *trigonatus* |
| Loricata | *Simosuchus* | *clarki* |
| Loricata | *Stomatosuchus* | *inermis* |
| Loricata | *Tomistoma* | *schlegelii* |
| Loricata | *Tomistoma* | *schlegelii* |
| Mammaliaform | *Abrawayaomys* | *ruschii* |
| Mammaliaform | *Acerodon* | *celebensis* |
| Mammaliaform | *Acerodon* | *jubatus* |
| Mammaliaform | *Acinonyx* | *jubatus* |
| Mammaliaform | *Acomys* | *cahirinus* |
| Mammaliaform | *Aconaemys* | *sagei* |
| Mammaliaform | *Acrobates* | *pygmaeus* |
| Mammaliaform | *Acrobates* | *pygmaeus* |
| Mammaliaform | *Aepyceros* | *melampus* |
| Mammaliaform | *Ailuropoda* | *melanoleuca* |
| Mammaliaform | *Ailurus* | *fulgens* |
| Mammaliaform | *Akodon* | *kofordi* |
| Mammaliaform | *Alcelaphus* | *buselaphus* |

| | | |
|---|---|---|
| Mammaliaform | *Alouatta* | *seniculus* |
| Mammaliaform | *Amblonyx* | *cinerea* |
| Mammaliaform | *Ametrida* | *centurio* |
| Mammaliaform | *Ametrida* | *centurio* |
| Mammaliaform | *Ammospermophilus* | *nelsoni* |
| Mammaliaform | *Ammotragus* | *lervia* |
| Mammaliaform | *Andalgalomys* | *pearsoni* |
| Mammaliaform | *Anoura* | *caudifer* |
| Mammaliaform | *Anoura* | *geoffroyi* |
| Mammaliaform | *Antilocapra* | *Antilocapra Americana* |
| Mammaliaform | *Antrozous* | *pallidus* |
| Mammaliaform | *Antrozous* | *pallidus* |
| Mammaliaform | *Aonyx* | *capensis* |
| Mammaliaform | *Aotus* | *trivirgatus* |
| Mammaliaform | *Aplodontia* | *rufa* |
| Mammaliaform | *Arctocebus* | *aureus* |
| Mammaliaform | *Arctonyx* | *collaris* |
| Mammaliaform | *Ariteus* | *flavescens* |
| Mammaliaform | *Artibeus* | *anderseni* |
| Mammaliaform | *Artibeus* | *cinereus* |
| Mammaliaform | *Asellia* | *tridens* |
| Mammaliaform | *Atelerix* | *albiventris* |
| Mammaliaform | *Ateles* | *geoffroyi* |
| Mammaliaform | *Atilax* | *paludinosus* |
| Mammaliaform | *Babyrousa* | *celebensis* |
| Mammaliaform | *Balantiopteryx* | *plicata* |
| Mammaliaform | *Balantiopteryx* | *plicata* |
| Mammaliaform | *Bassaricyon* | *gabbii* |
| Mammaliaform | *Bassariscus* | *astutus* |
| Mammaliaform | *Bettongia* | *penicillata* |
| Mammaliaform | *Bison* | *antiquus* |
| Mammaliaform | *Bison* | *bison* |
| Mammaliaform | *Bison* | *Bison bison* |
| Mammaliaform | *Blarina* | *brevicauda* |
| Mammaliaform | *Borhyaena* | *tuberata* |
| Mammaliaform | *Bos* | *taurus* |
| Mammaliaform | *Boselaphus* | *tragocamelus* |
| Mammaliaform | *Brachyphylla* | *cavernarum* |
| Mammaliaform | *Brachyphylla* | *cavernarum* |
| Mammaliaform | *Brachyteles* | *arachnoides* |
| Mammaliaform | *Cabassous* | *unicinctus* |
| Mammaliaform | *Caenolestes* | *convelatus* |
| Mammaliaform | *Calippus* | *martini* |
| Mammaliaform | *Callicebus* | *moloch* |
| Mammaliaform | *Callimico* | *goeldii* |

| | | |
|---|---|---|
| Mammaliaform | *Callimico* | *goeldii* |
| Mammaliaform | *Callithrix* | *jacchus* |
| Mammaliaform | *Callithrix* | *pygmaea* |
| Mammaliaform | *Callorhinus* | *ursinus* |
| Mammaliaform | *Caluromys* | *derbianus* |
| Mammaliaform | *Caluromysiops* | *irrupta* |
| Mammaliaform | *Canis* | *lupus* |
| Mammaliaform | *Cannomys* | *badius* |
| Mammaliaform | *Capra* | *hircus* |
| Mammaliaform | *Capromys* | *pilorides* |
| Mammaliaform | *Caracal* | *caracal* |
| Mammaliaform | *Carlito* | *syrichta* |
| Mammaliaform | *Carollia* | *brevicauda* |
| Mammaliaform | *Carollia* | *subrufa* |
| Mammaliaform | *Casinycteris* | *ophiodon* |
| Mammaliaform | *Castor* | *canadensis* |
| Mammaliaform | *Castor* | *canadensis* |
| Mammaliaform | *Cebus* | *capucinus* |
| Mammaliaform | *Cebus* | *versicolor* |
| Mammaliaform | *Centurio* | *senex* |
| Mammaliaform | *Cephalopachus* | *bancanus* |
| Mammaliaform | *Cephalophus* | *ogilbly* |
| Mammaliaform | *Cercocebus* | *torquatus* |
| Mammaliaform | *Cercopithecus* | *neglectus* |
| Mammaliaform | *Cerdocyon* | *thous* |
| Mammaliaform | *Cervalces* | *scotti* |
| Mammaliaform | *Chaerephon* | *jobensis* |
| Mammaliaform | *Chaerephon* | *plicatus* |
| Mammaliaform | *Chaetodipus* | *intermedius* |
| Mammaliaform | *Chaetophractus* | *vellerosus* |
| Mammaliaform | *Chaetophractus* | *villosus* |
| Mammaliaform | *Cheiromeles* | *parvidens* |
| Mammaliaform | *Chiroderma* | *salvini* |
| Mammaliaform | *Chiroderma* | *villosum* |
| Mammaliaform | *Chironectes* | *minimus* |
| Mammaliaform | *Chiropotes* | *satanas* |
| Mammaliaform | *Chlamyphorus* | *truncatus* |
| Mammaliaform | *Chlorocebus* | *aethiops* |
| Mammaliaform | *Chlorocebus* | *sabaeus* |
| Mammaliaform | *Chlorotalpa* | *sclateri* |
| Mammaliaform | *Chodsigoa* | *hypsibia* |
| Mammaliaform | *Choeroniscus* | *godmani* |
| Mammaliaform | *Choeroniscus* | *godmani* |
| Mammaliaform | *Choeronycteris* | *mexicana* |

| | | |
|---|---|---|
| Mammaliaform | *Choeronycteris* | *mexicana* |
| Mammaliaform | *Chrotopterus* | *auritus* |
| Mammaliaform | *Chrotopterus* | *auritus* |
| Mammaliaform | *Chrysocyon* | *brachyurus* |
| Mammaliaform | *Civettictis* | *civetta* |
| Mammaliaform | *Cladosictis* | *patagonica* |
| Mammaliaform | *Clethrionomys* | *dawsoni* |
| Mammaliaform | *Colobus* | *angolensis* |
| Mammaliaform | *Colobus* | *polykomos* |
| Mammaliaform | *Condylura* | *cristata* |
| Mammaliaform | *Conepatus* | *humboldti* |
| Mammaliaform | *Connochaetes* | *taurinus* |
| Mammaliaform | *Corynorhinus* | *macrotis* |
| Mammaliaform | *Corynorhinus* | *rafinesquii* |
| Mammaliaform | *Cricetomys* | *gambianus* |
| Mammaliaform | *Crocidura* | *foetida* |
| Mammaliaform | *Crocuta* | *crocuta* |
| Mammaliaform | *Crossarchus* | *obscurus* |
| Mammaliaform | *Cryptomys* | *hottentotus* |
| Mammaliaform | *Cryptoprocta* | *ferox* |
| Mammaliaform | *Cryptotis* | *magna* |
| Mammaliaform | *Ctenomys* | *opimus* |
| Mammaliaform | *Ctenomys* | *sociabilis* |
| Mammaliaform | *Cynictis* | *penicillata* |
| Mammaliaform | *Cynocephalus* | *sp.* |
| Mammaliaform | *Cynocephalus* | *volans* |
| Mammaliaform | *Cynogale* | *bennettii* |
| Mammaliaform | *Cynomops* | *abrasus* |
| Mammaliaform | *Cynomops* | *planirostris* |
| Mammaliaform | *Cynopterus* | *brachyotis* |
| Mammaliaform | *Cynopterus* | *brachyotis* |
| Mammaliaform | *Dama* | *dama* |
| Mammaliaform | *Damaliscus* | *lunatus* |
| Mammaliaform | *Dasyprocta* | *punctata* |
| Mammaliaform | *Dasypus* | *kappleri* |
| Mammaliaform | *Dasypus* | *novemcinctus* |
| Mammaliaform | *Dasypus* | *sabanicola* |
| Mammaliaform | *Dasyurus* | *hallucatus* |
| Mammaliaform | *Daubentonia* | *lateral* |
| Mammaliaform | *Daubentonia* | *madagascariensis* |
| Mammaliaform | *Dendrohyrax* | *dorsalis* |
| Mammaliaform | *Dendrohyrax* | *arboreus* |
| Mammaliaform | *Desmodus* | *rotundus* |
| Mammaliaform | *Desmodus* | *rotundus* |

| | | |
|---|---|---|
| Mammaliaform | *Dicerorhinus* | *sumatrensis* |
| Mammaliaform | *Diclidurus* | *albus* |
| Mammaliaform | *Didelphis* | *sp* |
| Mammaliaform | *Didelphis* | *virginiana* |
| Mammaliaform | *Diphylla* | *ecaudata* |
| Mammaliaform | *Diphylla* | *ecaudata* |
| Mammaliaform | *Dipodomys* | *leucogenys* |
| Mammaliaform | *Dipus* | *sagitta* |
| Mammaliaform | *Dobsonia* | *minor* |
| Mammaliaform | *Dorcopsis* | *muelleri* |
| Mammaliaform | *Dymecodon* | *pilirostris* |
| Mammaliaform | *Echinosorex* | *gymnura* |
| Mammaliaform | *Echymipera* | *rufescens* |
| Mammaliaform | *Eidolon* | *helvum* |
| Mammaliaform | *Eira* | *barbara* |
| Mammaliaform | *Elephantulus* | *brachyrhynchus* |
| Mammaliaform | *Elephas* | *maximus* |
| Mammaliaform | *Eliomys* | *quecinus* |
| Mammaliaform | *Eliomys* | *quecinus* |
| Mammaliaform | *Emballonura* | *alecto* |
| Mammaliaform | *Emballonura* | *alecto* |
| Mammaliaform | *Enhydra* | *lutris* |
| Mammaliaform | *Eonycteris* | *major* |
| Mammaliaform | *Eonycteris* | *robusta* |
| Mammaliaform | *Epomophorus* | *gambianus* |
| Mammaliaform | *Epomophorus* | *labiatus* |
| Mammaliaform | *Epomops* | *franqueti* |
| Mammaliaform | *Epomops* | *franqueti* |
| Mammaliaform | *Eptesicus* | *andinus* |
| Mammaliaform | *Eptesicus* | *andinus* |
| Mammaliaform | *Equus* | *burchelli* |
| Mammaliaform | *Equus* | *ferus* |
| Mammaliaform | *Equus* | *caballus* |
| Mammaliaform | *Erethizon* | *epixanthum* |
| Mammaliaform | *Erethizon* | |
| Mammaliaform | *Erignathus* | *barbatus* |
| Mammaliaform | *Erinaceus* | *europaeus* |
| Mammaliaform | *Erophylla* | *bombifrons* |
| Mammaliaform | *Erophylla* | *sezekorni* |
| Mammaliaform | *Erophylla* | *sezekorni* |
| Mammaliaform | *Erythrocebus* | *patas* |
| Mammaliaform | *Eulemur* | *fulvus* |
| Mammaliaform | *Eumops* | *sonoriensis* |

| | | |
|---|---|---|
| Mammaliaform | *Euoticus* | *elegantulus* |
| Mammaliaform | *Euphractus* | *sexcinctus* |
| Mammaliaform | *Eupleres* | *goudotii* |
| Mammaliaform | *Eurohippus* | *parvulus* |
| Mammaliaform | *Euroscaptor* | *micrura* |
| Mammaliaform | *Eutatus* | *sequini* |
| Mammaliaform | *Felis* | *silvestris* |
| Mammaliaform | *Fossa* | *fossana* |
| Mammaliaform | *Galago* | *senegalensis* |
| Mammaliaform | *Galemys* | *pyrenaicus* |
| Mammaliaform | *Galeopterus* | *variegatus* |
| Mammaliaform | *Galerella* | *sanguinea* |
| Mammaliaform | *Galidia* | *elegans* |
| Mammaliaform | *Galidictis* | *fasciata* |
| Mammaliaform | *Gazella* | *subgutturosa* |
| Mammaliaform | *Genetta* | *piscivora* |
| Mammaliaform | *Gerbillus* | *watersi* |
| Mammaliaform | *Giraffa* | *camelopardalis* |
| Mammaliaform | *Giraffa* | *camelopardalis* |
| Mammaliaform | *Glaucomys* | *sabrinus* |
| Mammaliaform | *Glossophaga* | *longirostris* |
| Mammaliaform | *Glossophaga* | *soricina* |
| Mammaliaform | *Glyphonycteris* | *sylvestris* |
| Mammaliaform | *Gorilla* | *gorilla* |
| Mammaliaform | *Gorilla* | *gorilla* |
| Mammaliaform | *Gracilinanus* | *agilis* |
| Mammaliaform | *Graphiurus* | *nagtglasii* |
| Mammaliaform | *Gulo* | *gulo* |
| Mammaliaform | *Hadrocodium* | *sp* |
| Mammaliaform | *Hadrocodium* | *sp* |
| Mammaliaform | *Haldanodon* | *sp* |
| Mammaliaform | *Haldanodon* | *sp* |
| Mammaliaform | *Hapalemur* | *griseus* |
| Mammaliaform | *Haplonycteris* | *fischeri* |
| Mammaliaform | *Haplonycteris* | *fischeri* |
| Mammaliaform | *Haramiyavia* | *sp* |
| Mammaliaform | *Harpyionycteris* | *celebensis* |
| Mammaliaform | *Helogale* | *undulata* |
| Mammaliaform | *Hemibelideus* | *lemuroides* |
| Mammaliaform | *Hemicentetes* | *semispinosus* |
| Mammaliaform | *Hemiechinus* | *auritus* |
| Mammaliaform | *Hemigalus* | *derbyanus* |
| Mammaliaform | *Herpestes* | *javanicus* |

| | | |
|---|---|---|
| Mammaliaform | *Hesperoptenus* | *tickelli* |
| Mammaliaform | *Hesperoptenus* | *tickelli* |
| Mammaliaform | *Heterocephalus* | *glaber* |
| Mammaliaform | *Hipposideros* | *diadema* |
| Mammaliaform | *Hipposideros* | *diadema* |
| Mammaliaform | *Hipposideros* | *maggietaylorae* |
| Mammaliaform | *Holochilus* | *sciureus* |
| Mammaliaform | *Homo* | *sapiens* |
| Mammaliaform | *Homo* | *sapiens* |
| Mammaliaform | *Hoolock* | *hoolock* |
| Mammaliaform | *Hyaena* | *brunnea* |
| Mammaliaform | *Hydrictis* | *macullicollis* |
| Mammaliaform | *Hydrochoerus* | *hydrochaeris* |
| Mammaliaform | *Hydropotes* | *inermis* |
| Mammaliaform | *Hydrurga* | *leptonyx* |
| Mammaliaform | *Hylobates* | *lar* |
| Mammaliaform | *Hylobates* | *moloch* |
| Mammaliaform | *Hylobates* | *muelleri* |
| Mammaliaform | *Hylobates* | *sp* |
| Mammaliaform | *Hylonycteris* | *underwoodi* |
| Mammaliaform | *Hylonycteris* | *underwoodi* |
| Mammaliaform | *Hypnomys* | *morpheus* |
| Mammaliaform | *Hypsugo* | *savii* |
| Mammaliaform | *Hyracotherium* | *sp.* |
| Mammaliaform | *Hystrix* | *cristata* |
| Mammaliaform | *Hystrix* | *indica* |
| Mammaliaform | *Ichneumia* | *albicauda* |
| Mammaliaform | *Idionycteris* | *phyllotis* |
| Mammaliaform | *Idionycteris* | *phyllotis* |
| Mammaliaform | *Iomys* | *horsefieldii* |
| Mammaliaform | *Kerivoula* | *pellucida* |
| Mammaliaform | *Kerivoula* | *pellucida* |
| Mammaliaform | *Kobus* | *kob* |
| Mammaliaform | *Kuehneotherium* | *prasecursoris* |
| Mammaliaform | *Kuehneotherium* | *sp* |
| Mammaliaform | *Lagostomus* | *maximus* |
| Mammaliaform | *Lagothrix* | *lagotricha* |
| Mammaliaform | *Lasionycteris* | *noctivagans* |
| Mammaliaform | *Lasionycteris* | *noctivagans* |
| Mammaliaform | *Lasiorhinus* | *krefftii* |
| Mammaliaform | *Lasiorhinus* | *latifrons* |
| Mammaliaform | *Lasiurus* | *borealis* |
| Mammaliaform | *Lasiurus* | *intermedius* |

| | | |
|---|---|---|
| Mammaliaform | *Lavia* | *frons* |
| Mammaliaform | *Lemur* | *catta* |
| Mammaliaform | *Leontocebus* | *fuscicollis* |
| Mammaliaform | *Leontopithecus* | *rosalia* |
| Mammaliaform | *Leopardus* | *geoffroyi* |
| Mammaliaform | *Leopardus* | *pardalis* |
| Mammaliaform | *Lepilemur* | *mustelinus* |
| Mammaliaform | *Leptailurus* | *serval* |
| Mammaliaform | *Leptonycteris* | *yerbabuenae* |
| Mammaliaform | *Leptonycteris* | *yerbabuenae* |
| Mammaliaform | *Lepus* | *bairdii* |
| Mammaliaform | *Lestoros* | *inca* |
| Mammaliaform | *Liomys* | *irroratus* |
| Mammaliaform | *Lionycteris* | *spurrelli* |
| Mammaliaform | *Lionycteris* | *spurrelli* |
| Mammaliaform | *Lonchophylla* | *robusta* |
| Mammaliaform | *Lonchophylla* | *robusta* |
| Mammaliaform | *Lonchorhina* | *aurita* |
| Mammaliaform | *Lontra* | *canadensis* |
| Mammaliaform | *Lontra* | *felina* |
| Mammaliaform | *Lontra* | *longicaudis* |
| Mammaliaform | *Lophocebus* | *albigena* |
| Mammaliaform | *Lophocebus* | *albigena* |
| Mammaliaform | *Lophostoma* | *brasiliense* |
| Mammaliaform | *Loris* | *lydekkerianus* |
| Mammaliaform | *Lutra* | *lutra* |
| Mammaliaform | *Lutreolina* | *crassicaudata* |
| Mammaliaform | *Lutrogale* | *perspicillata* |
| Mammaliaform | *Lycalopex* | *culpaeus* |
| Mammaliaform | *Lycaon* | *pictus* |
| Mammaliaform | *Lynx* | *lynx* |
| Mammaliaform | *Macaca* | *fascicularis* |
| Mammaliaform | *Macaca* | *fuscata* |
| Mammaliaform | *Macaca* | *mulatta* |
| Mammaliaform | *Macaca* | *nemestrina* |
| Mammaliaform | *Macroeuphractus* | *outesi* |
| Mammaliaform | *Macroglossus* | *minimus* |
| Mammaliaform | *Macroglossus* | *minimus* |
| Mammaliaform | *Macropus* | *rufus* |
| Mammaliaform | *Macropus* | *sp* |
| Mammaliaform | *Macrotis* | *lagotis* |
| Mammaliaform | *Macrotus* | *waterhousii* |
| Mammaliaform | *Macrotus* | *waterhousii* |
| Mammaliaform | *Madoqua* | *kirkii* |

| | | |
|---|---|---|
| Mammaliaform | *Mammut* | *pacificus* |
| Mammaliaform | *Mandrillus* | *leucophaeus* |
| Mammaliaform | *Manis* | *tricuspis* |
| Mammaliaform | *Maotherium* | *sp* |
| Mammaliaform | *Marmosops* | *fuscatus* |
| Mammaliaform | *Marmota* | *flaviventris* |
| Mammaliaform | *Martes* | *pennanti* |
| Mammaliaform | *Megaconus* | *sp* |
| Mammaliaform | *Megaderma* | *lyra* |
| Mammaliaform | *Megaderma* | *spasma* |
| Mammaliaform | *Megaptera* | *novaeangliae* |
| Mammaliaform | *Megaptera* | *novaeangliae* |
| Mammaliaform | *Megasorex* | *gigas* |
| Mammaliaform | *Meles* | *meles* |
| Mammaliaform | *Mellivora* | *capensis* |
| Mammaliaform | *Melogale* | *moschata* |
| Mammaliaform | *Mephitis* | *mephitis* |
| Mammaliaform | *Merychippus* | *insignis* |
| Mammaliaform | *Merychippus* | *sp.* |
| Mammaliaform | *Mesohippus* | *sp.* |
| Mammaliaform | *Mesohippus* | *sp.* |
| Mammaliaform | *Metachirus* | *nudicaudatus* |
| Mammaliaform | *Micoureus* | *regina* |
| Mammaliaform | *Microcebus* | *murinus* |
| Mammaliaform | *Microdipodops* | *pallidus* |
| Mammaliaform | *Microdocodon* | *sp* |
| Mammaliaform | *Microgale* | *brevicaudata* |
| Mammaliaform | *Micronycteris* | *hirsuta* |
| Mammaliaform | *Micronycteris* | *hirsuta* |
| Mammaliaform | *Microtus* | *californicus* |
| Mammaliaform | *Mimon* | *cozumelae* |
| Mammaliaform | *Mimon* | *cozumelae* |
| Mammaliaform | *Miniopterus* | *australis* |
| Mammaliaform | *Miniopterus* | *schreibersii* |
| Mammaliaform | *Miopithecus* | *talapoin* |
| Mammaliaform | *Mogera* | *wogura* |
| Mammaliaform | *Molossops* | *temminckii* |
| Mammaliaform | *Molossops* | *temminckii* |
| Mammaliaform | *Molossus* | *currentium* |
| Mammaliaform | *Molossus* | *molossus* |
| Mammaliaform | *Monodelphis* | *sp* |
| Mammaliaform | *Monophyllus* | *redmani* |
| Mammaliaform | *Monophyllus* | *redmani* |
| Mammaliaform | *Morganucodon* | *sp* |
| Mammaliaform | *Morganucodon* | *sp* |

| | | |
|---|---|---|
| Mammaliaform | *Morganucodon* | *watsoni* |
| Mammaliaform | *Mormoops* | *megalophylla* |
| Mammaliaform | *Mormoops* | *megalophylla* |
| Mammaliaform | *Mormopterus* | *acetabulosus* |
| Mammaliaform | *Mormopterus* | *acetabulosus* |
| Mammaliaform | *Mungos* | *mungo* |
| Mammaliaform | *Murina* | *aurata* |
| Mammaliaform | *Murina* | *aurata* |
| Mammaliaform | *Mus* | *musculus* |
| Mammaliaform | *Musonycteris* | *harrisoni* |
| Mammaliaform | *Musonycteris* | *harrisoni* |
| Mammaliaform | *Mustela* | *frenata* |
| Mammaliaform | *Mydaus* | *javanensis* |
| Mammaliaform | *Myocastor* | *coypus* |
| Mammaliaform | *Myodes* | *gapperi* |
| Mammaliaform | *Myodes* | *phaeus* |
| Mammaliaform | *Myotis* | *albescens* |
| Mammaliaform | *Myotis* | *californicus* |
| Mammaliaform | *Myrmecobius* | *fasciatus* |
| Mammaliaform | *Nandinia* | *binotata* |
| Mammaliaform | *Nandinia* | *binotata* |
| Mammaliaform | *Nanger* | *dama* |
| Mammaliaform | *Nasalis* | *larvatus* |
| Mammaliaform | *Nasalis* | *larvatus* |
| Mammaliaform | *Nasua* | *narica* |
| Mammaliaform | *Natalus* | *tumidirostris* |
| Mammaliaform | *Neacomys* | *musseri* |
| Mammaliaform | *Neofelis* | *nebulosa* |
| Mammaliaform | *Neomys* | *fodiens* |
| Mammaliaform | *Neoromicia* | *capensis* |
| Mammaliaform | *Neoromicia* | *capensis* |
| Mammaliaform | *Neotoma* | *nevadensis* |
| Mammaliaform | *Neotragus* | *batesi* |
| Mammaliaform | *Neovison* | *vison* |
| Mammaliaform | *Neurotrichus* | *gibbsii* |
| Mammaliaform | *Noctilio* | *albiventris* |
| Mammaliaform | *Noctilio* | *leporinus* |
| Mammaliaform | *Nomascus* | *concolor* |
| Mammaliaform | *Notiosorex* | *crawfordi* |
| Mammaliaform | *Nyctalus* | *leisleri* |
| Mammaliaform | *Nyctereutes* | *procyonoides* |
| Mammaliaform | *Nycteris* | *hispida* |
| Mammaliaform | *Nycticebus* | *bengalensis* |
| Mammaliaform | *Nycticebus* | *coucang* |
| Mammaliaform | *Nycticeius* | *humeralis* |

| | | |
|---|---|---|
| Mammaliaform | *Nyctiellus* | *lepidus* |
| Mammaliaform | *Nyctimene* | *major* |
| Mammaliaform | *Nyctimene* | *rabori* |
| Mammaliaform | *Nyctinomops* | *femorosaccus* |
| Mammaliaform | *Nyctinomops* | *laticaudatus* |
| Mammaliaform | *Nyctomys* | *sumichrasti* |
| Mammaliaform | *Ochotona* | *princeps* |
| Mammaliaform | *Octodon* | *degus* |
| Mammaliaform | *Odobenus* | *rosmarus* |
| Mammaliaform | *Odocoileus* | *Odocoileus hemionus* |
| Mammaliaform | *Oecomys* | *mamorae* |
| Mammaliaform | *Okapia* | *johnstoni* |
| Mammaliaform | *Oligokyphus* | *sp* |
| Mammaliaform | *Ondatra* | |
| Mammaliaform | *Onychogalea* | *frenata* |
| Mammaliaform | *Orcinus* | *orca* |
| Mammaliaform | *Oreamnos* | *americanus* |
| Mammaliaform | *Ornithorhynchus* | *anatinus* |
| Mammaliaform | *Ornithorhynchus* | *sp* |
| Mammaliaform | *Oryctolagus* | *cuniculus* |
| Mammaliaform | *Oryx* | *dammah* |
| Mammaliaform | *Otocyon* | *megalotis* |
| Mammaliaform | *Otolemur* | *crassicaudatus* |
| Mammaliaform | *Otolemur* | *crassicaudatus* |
| Mammaliaform | *Otopteropus* | *cartilagonodus* |
| Mammaliaform | *Otopteropus* | *cartilagonodus* |
| Mammaliaform | *Ovibos* | *moschatus* |
| Mammaliaform | *Ovis* | *canadensis* |
| Mammaliaform | *Ovis Linnaeus* | |
| Mammaliaform | *Oxymycterus* | *hiska* |
| Mammaliaform | *Paguma* | *larvata* |
| Mammaliaform | *Pan* | *troglodytes* |
| Mammaliaform | *Panthera* | *leo* |
| Mammaliaform | *Panthera* | *pardus* |
| Mammaliaform | *Panthera* | *uncia* |
| Mammaliaform | *Papio* | *cynocephalus* |
| Mammaliaform | *Papio* | *ursinus* |
| Mammaliaform | *Paracynictis* | *selousi* |
| Mammaliaform | *Paradoxurus* | *hermaphroditus* |
| Mammaliaform | *Paraechinus* | *micropus* |
| Mammaliaform | *Parahyaena* | *brunnea* |
| Mammaliaform | *Paralomys* | *gerbillus* |
| Mammaliaform | *Parascalops* | *breweri* |
| Mammaliaform | *Pedetes* | *capensis* |
| Mammaliaform | *Penthetor* | *lucasi* |

| | | |
|---|---|---|
| Mammaliaform | *Penthetor* | *lucasi* |
| Mammaliaform | *Perameles* | *gunnii* |
| Mammaliaform | *Perameles* | *obesula* |
| Mammaliaform | *Perodicticus* | *potto* |
| Mammaliaform | *Perodicticus* | *potto* |
| Mammaliaform | *Peromyscus* | *crinitus* |
| Mammaliaform | *Peropteryx* | *kappleri* |
| Mammaliaform | *Peropteryx* | *kappleri* |
| Mammaliaform | *Petaurista* | *petaurista* |
| Mammaliaform | *Petaurus* | *breviceps* |
| Mammaliaform | *Petrodromus* | *sultani* |
| Mammaliaform | *Petrogale* | *brachyotis* |
| Mammaliaform | *Petrogale* | *penicillata* |
| Mammaliaform | *Petrogale* | *Petrogale penicillata* |
| Mammaliaform | *Phascogale* | *tapoatafa* |
| Mammaliaform | *Phascolarctos* | *cinereus* |
| Mammaliaform | *Philander* | *andersoni* |
| Mammaliaform | *Philantomba* | *monticola* |
| Mammaliaform | *Philantomba* | *monticolor* |
| Mammaliaform | *Philetor* | *brachypterus* |
| Mammaliaform | *Philetor* | *brachypterus* |
| Mammaliaform | *Phoca* | *vitulina* |
| Mammaliaform | *Phocoena* | *phocoena* |
| Mammaliaform | *Phocoena* | *sinus* |
| Mammaliaform | *Phyllops* | *falcatus* |
| Mammaliaform | *Phyllops* | *falcatus* |
| Mammaliaform | *Phyllostomus* | *discolor* |
| Mammaliaform | *Phyllostomus* | *discolor* |
| Mammaliaform | *Phyllotis* | *caprinus* |
| Mammaliaform | *Piliocolobus* | *badius* |
| Mammaliaform | *Piliocolobus* | *badius* |
| Mammaliaform | *Pipistrellus* | *coromandra* |
| Mammaliaform | *Pipistrellus* | *sp* |
| Mammaliaform | *Pithecia* | *pithecia* |
| Mammaliaform | *Platyrrhinus* | *brachycephalus* |
| Mammaliaform | *Platyrrhinus* | *helleri* |
| Mammaliaform | *Plecotus* | *auritus* |
| Mammaliaform | *Plecotus* | *auritus* |
| Mammaliaform | *Pongo* | *abelii* |
| Mammaliaform | *Pongo* | *pygmaeus* |
| Mammaliaform | *Potorous* | *tridactylus* |
| Mammaliaform | *Potos* | *flavus* |
| Mammaliaform | *Presbytis* | *femoralis* |
| Mammaliaform | *Priodontes* | *maximus* |
| Mammaliaform | *Prionailurus* | *viverrinus* |

| | | |
|---|---|---|
| Mammaliaform | *Prionodon* | *pardicolor* |
| Mammaliaform | *Procavia* | *capensis* |
| Mammaliaform | *Procolobus* | *verus* |
| Mammaliaform | *Procyon* | *lotor* |
| Mammaliaform | *Promops* | *centralis* |
| Mammaliaform | *Promops* | *nasutus* |
| Mammaliaform | *Propithecus* | *coquereli* |
| Mammaliaform | *Proteles* | *cristatus* |
| Mammaliaform | *Prothylacinus* | *patagonicus* |
| Mammaliaform | *Pseudocheirus* | *cooki* |
| Mammaliaform | *Pseudocheirus* | *peregrinus* |
| Mammaliaform | *Ptenochirus* | *jagori* |
| Mammaliaform | *Ptenochirus* | *jagori* |
| Mammaliaform | *Pteromys* | *volans* |
| Mammaliaform | *Pteronotus* | *davyi* |
| Mammaliaform | *Pteronotus* | *davyi* |
| Mammaliaform | *Pteronotus* | *parnellii* |
| Mammaliaform | *Pteronura* | *brasiliensis* |
| Mammaliaform | *Pteropus* | *hypomelanus* |
| Mammaliaform | *Pteropus* | *vampyrus* |
| Mammaliaform | *Ptilocercus* | *lowii* |
| Mammaliaform | *Puma* | *concolor* |
| Mammaliaform | *Punomys* | *kofordi* |
| Mammaliaform | *Pygathrix* | *nemaeus* |
| Mammaliaform | *Pygoderma* | *bilabiatum* |
| Mammaliaform | *Pygoderma* | *bilabiatum* |
| Mammaliaform | *Redunca* | *arundinum* |
| Mammaliaform | *Reithrodontomys* | *raviventris* |
| Mammaliaform | *Repenomamus* | *sp* |
| Mammaliaform | *Rhagomys* | *longilingua* |
| Mammaliaform | *Rheithrosciurus* | *small* |
| Mammaliaform | *Rhinolophus* | *arcuatus* |
| Mammaliaform | *Rhinolophus* | *ferrumequinum* |
| Mammaliaform | *Rhinophylla* | *pumilio* |
| Mammaliaform | *Rhinophylla* | *pumilio* |
| Mammaliaform | *Rhinopithecus* | *roxellana* |
| Mammaliaform | *Rhinopoma* | *hardwickii* |
| Mammaliaform | *Rhipidomys* | *gardneri* |
| Mammaliaform | *Rhogeessa* | *tumida* |
| Mammaliaform | *Rhogeessa* | *tumida* |
| Mammaliaform | *Rhynchonycteris* | *naso* |
| Mammaliaform | *Rousettus* | *aegyptiacus* |
| Mammaliaform | *Rousettus* | *aegyptiacus* |
| Mammaliaform | *Saccolaimus* | *saccolaimus* |
| Mammaliaform | *Saccopteryx* | *canescens* |

| | | |
|---|---|---|
| Mammaliaform | *Saccopteryx* | *leptura* |
| Mammaliaform | *Saguinus* | *geoffroyi* |
| Mammaliaform | *Saimiri* | *sciureus* |
| Mammaliaform | *Sapajus* | *apella* |
| Mammaliaform | *Sarcophilus* | *harrisii* |
| Mammaliaform | *Sarcophilus* | *laniarius* |
| Mammaliaform | *Scalopus* | *aquaticus* |
| Mammaliaform | *Scapanus* | *latimanus* |
| Mammaliaform | *Scotophilus* | *heathi* |
| Mammaliaform | *Scotophilus* | *heathii* |
| Mammaliaform | *Selevinia* | *betpakdalaensis* |
| Mammaliaform | *Semnopithecus* | *johnii* |
| Mammaliaform | *Shuotherium* | *sp* |
| Mammaliaform | *Sigmodon* | *ochrognathus* |
| Mammaliaform | *Sigmodontomys* | *aphrastus* |
| Mammaliaform | *Simias* | *concolor* |
| Mammaliaform | *Sinobaatar* | *sp* |
| Mammaliaform | *Sinoconodon* | *sp* |
| Mammaliaform | *Sinoconodon* | *sp* |
| Mammaliaform | *Smilodon* | *fatalis* |
| Mammaliaform | *Solenodon* | *paradoxus* |
| Mammaliaform | *Sorex* | *ornatus* |
| Mammaliaform | *Speothos* | *venaticus* |
| Mammaliaform | *Spermophilus* | *franklinii* |
| Mammaliaform | *Spilogale* | *putorius* |
| Mammaliaform | *Stenoderma* | *rufum* |
| Mammaliaform | *Stenoderma* | *rufum* |
| Mammaliaform | *Sturnira* | *ludovici* |
| Mammaliaform | *Sturnira* | *magna* |
| Mammaliaform | *Suncus* | *etruscus* |
| Mammaliaform | *Suricata* | *suricatta* |
| Mammaliaform | *Syconycteris* | *australis* |
| Mammaliaform | *Sylvilagus* | *bachmani* |
| Mammaliaform | *Symphalangus* | *syndactylus* |
| Mammaliaform | *Symphalangus* | *syndactylus* |
| Mammaliaform | *Tachyglossus* | *aculeatus* |
| Mammaliaform | *Tachyglossus* | *sp* |
| Mammaliaform | *Tachyoryctes* | *ibeanus* |
| Mammaliaform | *Tadarida* | *aegyptiaca* |
| Mammaliaform | *Tadarida* | *aegyptiaca* |
| Mammaliaform | *Talpa* | *europaea* |
| Mammaliaform | *Tamias* | *panamintinus* |
| Mammaliaform | *Tamiasciurus* | *hudsonicus* |
| Mammaliaform | *Taphozous* | *philippinensis* |
| Mammaliaform | *Taphozous* | *philippinensis* |

| | | |
|---|---|---|
| Mammaliaform | *Tapirella* | *bairdii* |
| Mammaliaform | *Tapirus* | *terrestris* |
| Mammaliaform | *Tarsius* | *syrichta* |
| Mammaliaform | *Tarsius* | *tarsier* |
| Mammaliaform | *Taxidea* | *taxus* |
| Mammaliaform | *Tayassu* | |
| Mammaliaform | *Tenrec* | *ecaudatus* |
| Mammaliaform | *Theropithecus* | *gelada* |
| Mammaliaform | *Theropithecus* | *gelada* |
| Mammaliaform | *Thomasomys* | *sp* |
| Mammaliaform | *Thomomys* | *bottae* |
| Mammaliaform | *Thoopterus* | *nigrescens* |
| Mammaliaform | *Thylacinus* | *cynocephalus* |
| Mammaliaform | *Thylacinus* | *cynocephalus* |
| Mammaliaform | *Thylogale* | *wilcoxi* |
| Mammaliaform | *Thyroptera* | *tricolor* |
| Mammaliaform | *Thyroptera* | *tricolor* |
| Mammaliaform | *Tolypeutes* | *matacus* |
| Mammaliaform | *Tonatia* | *bidens* |
| Mammaliaform | *Trachops* | *cirrhosus* |
| Mammaliaform | *Trachops* | *cirrhosus* |
| Mammaliaform | *Trachypithecus* | *cristatus* |
| Mammaliaform | *Trachypithecus* | *cristatus* |
| Mammaliaform | *Tragelaphus* | *strepsicerus* |
| Mammaliaform | *Tragulus* | *javanicus* |
| Mammaliaform | *Tremarctos* | *ornatus* |
| Mammaliaform | *Trichosurus* | *vulpecula* |
| Mammaliaform | *Trichosurus* | *vulpecula* |
| Mammaliaform | *Tupaia* | *belangeri* |
| Mammaliaform | *Tupaia* | *chrysogaster* |
| Mammaliaform | *Tupaia* | *dorsalis* |
| Mammaliaform | *Tylonycteris* | *pachypus* |
| Mammaliaform | *Tylonycteris* | *pachypus* |
| Mammaliaform | *Urocyon* | *cinereoargenteus* |
| Mammaliaform | *Urocyon* | *cinereoargenteus* |
| Mammaliaform | *Uroderma* | *bilobatum* |
| Mammaliaform | *Uroderma* | *bilobatum* |
| Mammaliaform | *Urotrichus* | *talpoides* |
| Mammaliaform | *Ursus* | *arctos* |
| Mammaliaform | *Vampyressa* | *nymphaea* |
| Mammaliaform | *Vampyriscus* | *nymphaea* |
| Mammaliaform | *Vampyrodes* | *caraccioli* |
| Mammaliaform | *Varecia* | *variegata* |
| Mammaliaform | *Varecia* | *variegata* |
| Mammaliaform | *Vassallia* | *maxima* |

| | | |
|---|---|---|
| Mammaliaform | *Vespertilio* | *murinus* |
| Mammaliaform | *Vilevolodon* | *sp* |
| Mammaliaform | *Viverra* | *tangalunga* |
| Mammaliaform | *Viverricula* | *indica* |
| Mammaliaform | *Vombatus* | *ursinus* |
| Mammaliaform | *Vulpes* | *vulpes* |
| Mammaliaform | *Zaedyus* | *pichiy* |
| Mammaliaform | *Zalophus* | *californicus* |
| Mammaliaform | *Zapus* | *trinotatus* |
| Mammaliaform | *Zygodontomys* | *brevicauda* |
| Neornithes | *Aechmophorus* | *occidentalis* |
| Neornithes | *Aegolius* | *acadicus* |
| Neornithes | *Aepyornis* | *sp* |
| Neornithes | *Alle* | *Alle* |
| Neornithes | *Amazona* | *ochrocephala* |
| Neornithes | *Anhinga* | *rufa* |
| Neornithes | *Anthracoceros* | *malayanus* |
| Neornithes | *Aptenodytes* | *patagonicus* |
| Neornithes | *Apteryx* | *australis* |
| Neornithes | *Ara* | *macao* |
| Neornithes | *Ara* | *macao* |
| Neornithes | *Aramus* | *guarauna* |
| Neornithes | *Athene* | *cunicularia* |
| Neornithes | *Athene* | *cunicularia* |
| Neornithes | *Athene* | *noctua* |
| Neornithes | *Branta* | *canadensis* |
| Neornithes | *Bubo* | *virginianus* |
| Neornithes | *Bucorvus* | *abyssinicus* |
| Neornithes | *Cardinalis* | *cardinalis* |
| Neornithes | *Cardinalis* | *cardinalis* |
| Neornithes | *Casuarius* | *sp* |
| Neornithes | *Centropus* | *sinensis* |
| Neornithes | *Chauna* | *torquata* |
| Neornithes | *Chauna* | *torquata* |
| Neornithes | *Circus* | *cyanus* |
| Neornithes | *Crypturellus* | *noctivagus* |
| Neornithes | *Dacelo* | *novaequineae* |
| Neornithes | *Dendrocygna* | *autumnalis* |
| Neornithes | *Diatryma* | *gigantea* |
| Neornithes | *Dinornis* | *sp* |
| Neornithes | *Dromas* | *ardeola* |
| Neornithes | *Euaegotheles* | *insignis* |
| Neornithes | *Eudocimus* | *ruber* |
| Neornithes | *Eudyptula* | *minor* |
| Neornithes | *Eulampis* | *jugularis* |

| | | |
|---|---|---|
| Neornithes | *Falco* | *sparverius* |
| Neornithes | *Fratercula* | *arctica* |
| Neornithes | *Fregata* | *magnificens* |
| Neornithes | *Gallinago* | *delicate* |
| Neornithes | *Glaucidium* | *gnoma* |
| Neornithes | *Glaucidium* | *periatum* |
| Neornithes | *Grus* | *canadensis* |
| Neornithes | *Gyps* | *fulvus* |
| Neornithes | *Haematopus* | *ostralegus* |
| Neornithes | *Herpetotheres* | *cachinnans* |
| Neornithes | *Hesperiphona* | *vespertinus* |
| Neornithes | *Himantopus* | *mexicanus* |
| Neornithes | *Hylocichla* | *mustelina* |
| Neornithes | *Larus* | *argentatus* |
| Neornithes | *Larus* | *argentatus* |
| Neornithes | *Larus* | *delawarensis* |
| Neornithes | *Lophorina* | *superba* |
| Neornithes | *Macrocephalon* | *maleo* |
| Neornithes | *Megaceryle* | *torquata* |
| Neornithes | *Microcarbo* | *africanus* |
| Neornithes | *Morus* | *bassanus* |
| Neornithes | *Musophaga* | *violacea* |
| Neornithes | *Ninox* | *novaeseelandiae* |
| Neornithes | *Ninox* | *theomacha* |
| Neornithes | *Numenius* | *americanus* |
| Neornithes | *Numenius* | *borealis* |
| Neornithes | *Numenius* | *phaeopus* |
| Neornithes | *Nymphicus* | *hollandicus* |
| Neornithes | *Onychoprion* | *fuscata* |
| Neornithes | *Opisthocomus* | *hoazin* |
| Neornithes | *Opisthocomus* | *hoazin* |
| Neornithes | *Opisthocomus* | *hoazin* |
| Neornithes | *Oxyura* | *jamaicensis* |
| Neornithes | *Pandion* | *haliaetus* |
| Neornithes | *Pelecanus* | *occidentalis* |
| Neornithes | *Phaethon* | *rubricauda* |
| Neornithes | *Philohela* | *minor* |
| Neornithes | *Phoebastria* | *nigripes* |
| Neornithes | *Phorusrhacid* | *sp* |
| Neornithes | *Platalea* | *ajaja* |
| Neornithes | *Platalea* | *ajaja* |
| Neornithes | *Porphyrio* | *mantelli* |
| Neornithes | *Priotelus* | *temnurus* |
| Neornithes | *Pterodroma* | *lessoni* |
| Neornithes | *Pulsatrix* | *perspicillata* |

| | | |
|---|---|---|
| Neornithes | *Rhea* | *sp* |
| Neornithes | *Rissa* | *tridactyla* |
| Neornithes | *Rynchops* | *flavirostris* |
| Neornithes | *Rynchops* | *niger* |
| Neornithes | *Rynchops* | *niger* |
| Neornithes | *Spatula* | *clupeata* |
| Neornithes | *Stercorarius* | *antarcticus* |
| Neornithes | *Sterna* | *dougalli* |
| Neornithes | *Sterna* | *hirundo* |
| Neornithes | *Sternula* | *albifrons* |
| Neornithes | *Strigops* | *habroptilus* |
| Neornithes | *Strigops* | *habroptilus* |
| Neornithes | *Strix* | *nebulosa* |
| Neornithes | *Struthio* | *sp* |
| Neornithes | *Struthio* | *Struthio Linnaeus* |
| Neornithes | *Thalasseus* | *bergii* |
| Neornithes | *Thalassornis* | *leuconotus* |
| Neornithes | *Tockus* | *erythrorhynchus* |
| Neornithes | *Trachyphonus* | *darnaudii* |
| Neornithes | *Tyto* | *alba* |
| Neornithes | *Upupa* | *epops* |
| Neornithes | *Uria* | *aalgae* |
| Ornithischia | *Ankylosaurus* | *magniventris* |
| Ornithischia | *Centrosaurus* | *apertus* |
| Ornithischia | *Lambeosaurinae* | *indet.* |
| Pelycosaur | *Dimetrodon* | *sp* |
| Pelycosaur | *Secodontosaurus* | *sp* |
| Pelycosaur | *Varanosaurus* | *acutirostris* |
| Placodermi | *Brachydeirus* | *sp* |
| Placodermi | *Brachyosteus* | *sp* |
| Placodermi | *Brontichthys* | *sp* |
| Placodermi | *Bullerichthys* | *sp* |
| Placodermi | *Campbellodus* | *sp* |
| Placodermi | *Camuropiscis* | *sp* |
| Placodermi | *Coccosteus* | *decipiens* |
| Placodermi | *Compagopiscis* | *sp* |
| Placodermi | *Dinichthys* | *cf. turberculatus* |
| Placodermi | *Dinomylostoma* | *sp* |
| Placodermi | *Dunkleosteus* | *terrelli* |
| Placodermi | *Eastmanosteus* | *sp* |
| Placodermi | *Enseosteus* | *sp* |
| Placodermi | *Fallacosteus* | *sp* |
| Placodermi | *Gorgonosteus* | *sp* |
| Placodermi | *Hadrosteus* | *rapax* |
| Placodermi | *Harrytoombsia* | *sp* |

| | | |
|---|---|---|
| Placodermi | *Heintzichthys* | *gouldii* |
| Placodermi | *Hussakofia* | *sp* |
| Placodermi | *Incisoscutum* | *ritchiei* |
| Placodermi | *Kendrichthys* | *sp* |
| Placodermi | *Kimberleyichthys* | *sp* |
| Placodermi | *Latocamurus* | *sp* |
| Placodermi | *Mylostoma* | *sp* |
| Placodermi | *Oxyosteus* | *sp* |
| Placodermi | *Pachyosteus* | *bulla* |
| Placodermi | *Pholidosteus* | *friedeli* |
| Placodermi | *Plourdosteus* | *sp.* |
| Placodermi | *Selenosteus* | *brevis* |
| Placodermi | *Titanichthys* | *sp* |
| Placodermi | *Torosteus* | *sp* |
| Placodermi | *Watsonosteus* | *fletti* |
| Pterosauria | *Aetodactylus* | *halli* |
| Pterosauria | *Angustinaripterus* | *longicephalus* |
| Pterosauria | *Anhanguera* | *piscator* |
| Pterosauria | *Bakonydraco* | *galaczi* |
| Pterosauria | *Barbosania* | *gracilirostris* |
| Pterosauria | *Bergamodactylus* | *wildi* |
| Pterosauria | *Boreopterus* | *giganticus* |
| Pterosauria | *Caiuajara* | *dobruskii* |
| Pterosauria | *Campylognathoides* | *liasicus* |
| Pterosauria | *Campylognathoides* | *zitteli* |
| Pterosauria | *Coloborhynchus* | *robustus* |
| Pterosauria | *Coloborhynchus* | *spielbergi* |
| Pterosauria | *Cycnorhamphus* | *suevicus* |
| Pterosauria | *Darwinopterus* | *robustodens* |
| Pterosauria | *Dimorphodon* | *macronyx* |
| Pterosauria | *Dorygnathus* | *banthensis* |
| Pterosauria | *Dsungaripterus* | *sp* |
| Pterosauria | *Eudimorphodon* | *ranzii* |
| Pterosauria | *Feilongus* | *youngi* |
| Pterosauria | *Germanodactylus* | *cristatus* |
| Pterosauria | *Guidraco* | *venator* |
| Pterosauria | *Hamipterus* | *tianshanensis* |
| Pterosauria | *Ikrandraco* | *avatar* |
| Pterosauria | *Ikrandraco* | *avatar* |
| Pterosauria | *Istiodactylus* | *sinensis* |
| Pterosauria | *Jianchangnathus* | *robustus* |
| Pterosauria | *Ludodactylus* | *sibbicki* |
| Pterosauria | *Moganopterus* | *zhuiana* |
| Pterosauria | *Noripterus* | *sp* |
| Pterosauria | *Nurhachius* | *ignaciobritoi* |

| | | |
|---|---|---|
| Pterosauria | *Ornithocheirus* | *mesembrinus* |
| Pterosauria | *Painten* | *ptero* |
| Pterosauria | *Pteranodon* | *sp* |
| Pterosauria | *Pterodactylus* | *antiquus* |
| Pterosauria | *Pterodactylus* | *scolopaciceps* |
| Pterosauria | *Pterodaustro* | *guinazu* |
| Pterosauria | *Quetzalcoatlus* | *sp* |
| Pterosauria | *Raeticodactylus* | *filisurensis* |
| Pterosauria | *Rhamphorhynchus* | *muensteri* |
| Pterosauria | *Scaphognathus* | *crassirostris* |
| Pterosauria | *Shenzhoupterus* | *chaoyangensis* |
| Pterosauria | *Sinopterus* | *dongi* |
| Pterosauria | *Tapejara* | *wellnhoferi* |
| Pterosauria | *Tupandactylus* | *imperator* |
| Pterosauria | *Tupuxuara* | *leonardii* |
| Pterosauria | *Zhejiangopterus* | *linhaiensis* |
| Pterosauria | *Zhenyuanopterus* | *longirostris* |
| Sarcopterygii | *Allenypterus* | *sp* |
| Sarcopterygii | *Axelrodichthys* | *sp* |
| Sarcopterygii | *Chirodipterus* | *australis* |
| Sarcopterygii | *Dipterus* | *valenciennesi* |
| Sarcopterygii | *Eusthenopteron* | *foordi* |
| Sarcopterygii | *Eusthenopteron* | *foordi* |
| Sarcopterygii | *Gavinia* | *sp* |
| Sarcopterygii | *Gavinia* | *syntrips* |
| Sarcopterygii | *Griphognathus* | *whitei* |
| Sarcopterygii | *Guiyu* | *oneiros* |
| Sarcopterygii | *Holodipterus* | *sp* |
| Sarcopterygii | *Kenichthys* | *campbelli* |
| Sarcopterygii | *Latimeria* | *sp* |
| Sarcopterygii | *Lepidosiren* | *paradoxa* |
| Sarcopterygii | *Macropoma* | *sp* |
| Sarcopterygii | *Megalichthys* | *hibberti* |
| Sarcopterygii | *Megalichthys* | *laticeps* |
| Sarcopterygii | *Miguashaia* | *sp* |
| Sarcopterygii | *Neoceratodus* | *forsteri* |
| Sarcopterygii | *Orlovichthys* | *limnatis* |
| Sarcopterygii | *Osteolepis* | *sp* |
| Sarcopterygii | *Panderichthys* | *sp* |
| Sarcopterygii | *Pillararhynchus* | *sp* |
| Sarcopterygii | *Platycephalichthys* | *bischoffi* |
| Sarcopterygii | *Platyethmoidia* | *antarctica* |
| Sarcopterygii | *Polyplocodus* | *leptognathus* |
| Sarcopterygii | *Psarolepis* | *romeri* |
| Sarcopterygii | *Rhizodus* | *hibberti* |

| | | |
|---|---|---|
| Sarcopterygii | *Robinsondipterus* | *longi* |
| Sarcopterygii | *Soederberghia* | *groenlandica* |
| Sarcopterygii | *Styloichthys* | *sp* |
| Sarcopterygii | *Thursius* | *pholidotus* |
| Sarcopterygii | *Tiktaalik* | *roseae* |
| Sarcopterygii | *Tristichopterid* | *Rhipidistiangen* |
| Sarcopterygii | *Tristichopterus* | *sp* |
| Sarcopterygii | *Whiteia* | *sp* |
| Squamata | *Smaug* | *giganteus* |
| Squamata | *Acanthodactylus* | *erythrurus* |
| Squamata | *Acontias* | *lineatus* |
| Squamata | *Acrochordus* | *javanicus* |
| Squamata | *Aeluroscalabotes* | *felinus* |
| Squamata | *Agama* | *agama* |
| Squamata | *Agamodon* | *anguliceps* |
| Squamata | *Agkistrodon* | *contortrix* |
| Squamata | *Amblyodipsas* | *polylepis* |
| Squamata | *Ameiva* | *sp* |
| Squamata | *Amphiesma* | *stolata* |
| Squamata | *Amphiglossus* | *splendidus* |
| Squamata | *Amphisbaena* | *alba* |
| Squamata | *Amphisbaena* | *alba* |
| Squamata | *Amphisbaena* | *fuliginosa* |
| Squamata | *Ancylocranium* | *ionidesi* |
| Squamata | *Anniella* | *pulchra* |
| Squamata | *Anniella* | *pulchra* |
| Squamata | *Anolis* | *carolinensis* |
| Squamata | *Anops* | *kingii* |
| Squamata | *Aparallactus* | *capensis* |
| Squamata | *Aparallactus* | *modestus* |
| Squamata | *Aspidoscelis* | *tigris* |
| Squamata | *Atractaspis* | *aterrima* |
| Squamata | *Atractaspis* | *bibronii* |
| Squamata | *Bipes* | *biporus* |
| Squamata | *Bipes* | *canaliculatus* |
| Squamata | *Blanus* | *cinereus* |
| Squamata | *Boa* | *constrictor* |
| Squamata | *Boaedon* | *fuliginosus* |
| Squamata | *Bothrops* | *asper* |
| Squamata | *Brachylophus* | *fasciatus* |
| Squamata | *Bradypodion* | *pumilum* |
| Squamata | *Bronia* | *brasiliana* |
| Squamata | *Brookesia* | *sp* |
| Squamata | *Cadea* | *blanoides* |
| Squamata | *Cadea* | *palirostrata* |

| | | |
|---|---|---|
| Squamata | *Calabaria* | *reinhardtii* |
| Squamata | *Calabaria* | *reinhardtii* |
| Squamata | *Callopistes* | *maculatus* |
| Squamata | *Calotes* | *emma* |
| Squamata | *Candoia* | *carinata* |
| Squamata | *Causus* | *rhombeatus* |
| Squamata | *Chalarodon* | *madagascariensis* |
| Squamata | *Chalarodon* | *madagascariensis* |
| Squamata | *Chamaeleo* | *laevigatus* |
| Squamata | *Chilabothrus* | *angulifer* |
| Squamata | *Chilabothrus* | *striatus* |
| Squamata | *Chilorhinophis* | *geradi* |
| Squamata | *Chondrodactylus* | *bibronii* |
| Squamata | *Colobosaura* | *modesta* |
| Squamata | *Coluber* | *constrictor* |
| Squamata | *Cophotis* | *ceylanica* |
| Squamata | *Corallus* | *caninus* |
| Squamata | *Cordylosaurus* | *subtessellatus* |
| Squamata | *Cricosaura* | *sp* |
| Squamata | *Ctenomastax* | *parva* |
| Squamata | *Ctenosaura* | *pectinata* |
| Squamata | *Cylindrophis* | *ruffus* |
| Squamata | *Dalophia* | *pistillum* |
| Squamata | *Dasypeltis* | *scabra* |
| Squamata | *Delma* | *borea* |
| Squamata | *Diadophis* | *punctatus* |
| Squamata | *Dibamus* | *novaeguineae* |
| Squamata | *Dicrodon* | *lentiginosus* |
| Squamata | *Diploglossus* | *lessonae* |
| Squamata | *Diplometopon* | *zarudnyi* |
| Squamata | *Diporiphora* | *winneckei* |
| Squamata | *Dipsosaurus* | *dorsalid* |
| Squamata | *Draco* | *blanfordii* |
| Squamata | *Dyticonastis* | *rensbergeri* |
| Squamata | *Egernia* | *stokesii* |
| Squamata | *Enyalioides* | *laticeps* |
| Squamata | *Eryx* | *colubrinus* |
| Squamata | *Feylinia* | *currori* |
| Squamata | *Feylinia* | *polylepis* |
| Squamata | *Gallotia* | *galloti* |
| Squamata | *Gekko* | *gecko* |
| Squamata | *Gerrhosaurus* | *albigularis* |
| Squamata | *Gonatodes* | *albogularis* |
| Squamata | *Gyalopion* | *canum* |
| Squamata | *Gymnophthalmus* | *speciosus* |

| | | |
|---|---|---|
| Squamata | *Heloderma* | *horridum* |
| Squamata | *Heloderma* | *suspectum* |
| Squamata | *Hemitheconyx* | *caudicinctus* |
| Squamata | *Homoroselaps* | *lacteus* |
| Squamata | *Iguana* | *sp* |
| Squamata | *Japalura* | *polygonata* |
| Squamata | *Lacerta* | *viridis* |
| Squamata | *Lampropeltis* | *getula* |
| Squamata | *Lamprophis* | *fuliginosus* |
| Squamata | *Lanthanotus* | *borneensis* |
| Squamata | *Lanthanotus* | *borneensis* |
| Squamata | *Laticauda* | *colubrina* |
| Squamata | *Leiocephalus* | *barahonensis* |
| Squamata | *Leiolepis* | *belliana* |
| Squamata | *Leiosaurus* | *catamarcensis* |
| Squamata | *Leptophis* | *ahaetulla* |
| Squamata | *Lialis* | *burtonis* |
| Squamata | *Lialis* | *burtonis* |
| Squamata | *Liolaemus* | *bellii* |
| Squamata | *Liotyphlops* | *albirostris* |
| Squamata | *Loxocemus* | *bicolor* |
| Squamata | *Lycophidion* | *capense* |
| Squamata | *Masticophis* | *taeniatus* |
| Squamata | *Monopeltis* | *infuscata* |
| Squamata | *Monopeltis* | *rhodesianus* |
| Squamata | *Morunasaurus* | *annularis* |
| Squamata | *Natrix* | *natrix* |
| Squamata | *Oplurus* | *cyclurus* |
| Squamata | *Pachycalamus* | *brevis* |
| Squamata | *Pareas* | *hamptoni* |
| Squamata | *Petrosaurus* | *mearnsi* |
| Squamata | *Phelsuma* | *lineata* |
| Squamata | *Pholidobolus* | *montium* |
| Squamata | *Phrynocephalus* | *arabicus* |
| Squamata | *Platysaurus* | *guttatus* |
| Squamata | *Platysaurus* | *imperator* |
| Squamata | *Plica* | *plica* |
| Squamata | *Plotosaurus* | *bennisoni* |
| Squamata | *Pogona* | *vitticeps* |
| Squamata | *Polemon* | *christyi* |
| Squamata | *Polychrus* | *marmoratus* |
| Squamata | *Pristidactylus* | *torquatus* |
| Squamata | *Prosymna* | *meleagris* |
| Squamata | *Pseudopus* | *apodus* |
| Squamata | *Pseudopus* | *apodus* |

| | | |
|---|---|---|
| Squamata | *Ptyas* | *mucosus* |
| Squamata | *Python* | *molurus* |
| Squamata | *Rena* | *dulcis* |
| Squamata | *Rhacodactylus* | *auriculatus* |
| Squamata | *Rhampholeon* | *spectrum* |
| Squamata | *Rhineura* | *floridana* |
| Squamata | *Rhinoeura* | *floridana* |
| Squamata | *Saltuarius* | *cornutus* |
| Squamata | *Sceloporus* | *variabilis* |
| Squamata | *Shinisaurus* | *crocodilurus* |
| Squamata | *Smaug* | *giganteus* |
| Squamata | *Sonora* | *semiannulata* |
| Squamata | *Sphenodon* | *punctatus* |
| Squamata | *Sphenomorphus* | *solomonis* |
| Squamata | *Stenocercus* | *guentheri* |
| Squamata | *Takydromus* | *ocellatus* |
| Squamata | *Takydromus* | *wolteri* |
| Squamata | *Teius* | *teyou* |
| Squamata | *Thamnophis* | *marcianus* |
| Squamata | *Tiliqua* | *scincoides* |
| Squamata | *Trachylepis* | *quinquetaeniata* |
| Squamata | *Trimorphodon* | *biscutatus* |
| Squamata | *Trogonophis* | *wiegmanni* |
| Squamata | *Tropidophorus* | *misaminius* |
| Squamata | *Tupinambis* | *teguixin* |
| Squamata | *Tupinambis* | *teguixin* |
| Squamata | *Typhlophis* | *squamosus* |
| Squamata | *Uma* | *scoparia* |
| Squamata | *Ungaliophis* | *continentalis* |
| Squamata | *Uromastyx* | *aegyptius* |
| Squamata | *Uromastyx* | *sp* |
| Squamata | *Urostrophus* | *vautieri* |
| Squamata | *Uta* | *stansburiana* |
| Squamata | *Varanus* | *acanthurus* |
| Squamata | *Varanus* | *olivaceus* |
| Squamata | *Varanus* | *salvator* |
| Squamata | *Xantusia* | *vigilis* |
| Squamata | *Xenocalamus* | *bicolor* |
| Squamata | *Xenochrophis* | *piscator* |
| Squamata | *Xenosaurus* | *grandis* |
| Squamata | *Xenosaurus* | *grandis* |
| Squamata | *Xenosaurus* | *rectocollaris* |
| Squamata | *Zonosaurus* | *ornatus* |
| Stem Tetrapod | *Acanthostega* | *gunnari* |
| Stem Tetrapod | *Balanerpeton* | *woodi* |

| | | |
|---|---|---|
| Stem Tetrapod | *Crassigyrinus* | *scoticus* |
| Stem Tetrapod | *Crassigyrinus* | *scoticus* |
| Stem Tetrapod | *Dendrerpeton* | *acadianum* |
| Stem Tetrapod | *Densignathus* | *rowei* |
| Stem Tetrapod | *Elginerpeton* | *pancheni* |
| Stem Tetrapod | *Greererpeton* | *burkemorani* |
| Stem Tetrapod | *Ichthyostega* | *stensioei* |
| Stem Tetrapod | *Ichthyostega* | *stensioei* |
| Stem Tetrapod | *Ichthyostega* | *stensioei* |
| Stem Tetrapod | *Megalocephalus* | *pacycephalus* |
| Stem Tetrapod | *Megalocephalus* | *pacycephalus* |
| Stem Tetrapod | *Metaxygnathus* | *denticulus* |
| Stem Tetrapod | *Phonerpeton* | *pricei* |
| Stem Tetrapod | *Ventastega* | *curonica* |
| Stem Tetrapod | *Whatcheeria* | *deltae* |
| Stem Tetrapod | *Ymeria* | *denticulata* |
| Testudines | *Apalone* | *spinifera* |
| Testudines | *Chelydra* | *serpentina* |
| Testudines | *Chelydra* | *serpentina* |
| Testudines | *Dermochelys* | *coriacea* |
| Testudines | *Dermochelys* | *coriacea* |
| Testudines | *Dermochelys* | *sp* |
| Testudines | *Elseya* | *dentata* |
| Testudines | *Elseya* | *dentata* |
| Testudines | *Glyptemys* | *muhlenbergii* |
| Testudines | *Gopherus* | *polyphemus* |
| Testudines | *Lepidochelys* | *olivacea* |
| Testudines | *Lepidochelys* | *sp* |
| Testudines | *Plesiochelys* | *planiceps* |
| Testudines | *Protostega* | *sp* |
| Testudines | *Rhinochelys* | *pulchriceps* |
| Testudines | *Sternotherus* | *minor* |
| Testudines | *Stigmochelys* | *Stigmochelys pardalis* |
| Testudines | *Terlinguachelys* | *sp* |
| Testudines | *Toxochelys* | *latiremis* |
| Testudines | *Toxochelys* | *sp* |
| Theropoda | *Alioramus* | *sp* |
| Theropoda | *Anchiornis* | *huxleyi* |
| Theropoda | *Anzu* | *wyliei* |
| Theropoda | *Archaeopteryx* | *lithographica* |
| Theropoda | *Bambiraptor* | *feinbergorum* |
| Theropoda | *Bistahieversor* | *sp* |
| Theropoda | *Caenagnathus* | *collinsi* |
| Theropoda | *Caudipteryx* | *zoui* |
| Theropoda | *Citipati* | *osmolskae* |

| | | |
|---|---|---|
| Theropoda | *Compsognathus* | *longipes* |
| Theropoda | *Confuciusornis* | *sanctus* |
| Theropoda | *Daspletosaurus* | *sp* |
| Theropoda | *Deinocheirus* | *sp* |
| Theropoda | *Dilong* | *paradoxus* |
| Theropoda | *Dilophosaurus* | *wetherilli* |
| Theropoda | *Dromaeosaurus* | *albertensis* |
| Theropoda | *Eosinopteryx* | *sp* |
| Theropoda | *Epidexipteryx* | *sp* |
| Theropoda | *Erlikosaurus* | *andrewsi* |
| Theropoda | *Gallimimus* | *bullatus* |
| Theropoda | *Garudimimus* | *brevipes* |
| Theropoda | *Gigantoraptor* | *erlianensis* |
| Theropoda | *Gobivenator* | *mongoliensis* |
| Theropoda | *Gorgosaurus* | *libratus* |
| Theropoda | *Guanlong* | *sp* |
| Theropoda | *Haplocheirus* | *sp* |
| Theropoda | *Harpymimus* | *okladnikovi* |
| Theropoda | *Hesperornis* | *sp* |
| Theropoda | *Huanansaurus* | *ganzhouensis* |
| Theropoda | *Ichthyornis* | *sp* |
| Theropoda | *Jianchangosaurus* | *yixianensis* |
| Theropoda | *Khaan* | *sp* |
| Theropoda | *Linheraptor* | *exquisitus* |
| Theropoda | *Longipteryx* | *sp.* |
| Theropoda | *Nanotyrannus* | *sp* |
| Theropoda | *Nemegtomaia* | *sp* |
| Theropoda | *Ornitholestes* | *hermanni* |
| Theropoda | *Ornithomimus* | *sp* |
| Theropoda | *Ornithomimus* | *edmontonicus* |
| Theropoda | *Oviraptor* | *philoceratops* |
| Theropoda | *Pengornis* | *houi* |
| Theropoda | *Proceratosaurus* | *bradleyi* |
| Theropoda | *Rapaxavis* | *pani* |
| Theropoda | *Raptorex* | *sp* |
| Theropoda | *Rinchenia* | *mongoliensis* |
| Theropoda | *Sapeornis* | *sp* |
| Theropoda | *Scipionyx* | *sp* |
| Theropoda | *Segnosaurus* | *galbinensis* |
| Theropoda | *Shuvuuia* | *deserti* |
| Theropoda | *Sinornithomimus* | *sp* |
| Theropoda | *Sinosauropteryx* | *prima* |
| Theropoda | *Struthiomimus* | *altus* |
| Theropoda | *Tarbosaurus* | *baatar* |
| Theropoda | *Tsaagan* | *mangas* |

| | | |
|---|---|---|
| Theropoda | *Tyrannosaurus* | *rex* |
| Theropoda | *Velociraptor* | *mongoliensis* |
| Theropoda | *Yanornis* | *martini* |
| Theropoda | *Yi* | *qi* |
| Theropoda | *yulong* | *sp* |
| Theropoda | *Yutyrannus* | *sp* |
| Theropoda | *Zhenyuanlong* | *suni* |

**Table S1.**
List of vertebrate taxa sampled in the study. The initial morphospace was constructed with the entire dataset; subsequent analyses were done by taking genus means where necessary and retaining genus level data only. For full details see Data S1.

| Gene as named in GenBank | Reference |
|---|---|
| *ALX3* – ALX homeobox 3 | Jeong *et al*. (2008) (135) |
| *BMPER* – BMP binding endothelial regulator | Jeong *et al*. (2008) (135) |
| *CITED1* – Cbp/p300 interacting transactivator with Glu/Asp rich carboxy-terminal domain 1 | Jeong *et al*. (2008) (135) |
| *DLX1* – distal-less homeobox 1 | Jeong *et al*. (2008) (135) |
| *DLX2* – distal-less homeobox 2 | Jeong *et al*. (2008) (135) |
| *DLX3* – distal-less homeobox 3 | Graham (2002), Jeong *et al*. (2008) (133, 135) |
| *DLX4* – distal-less homeobox 4 | Jeong *et al*. (2008) (135) |
| *DLX5* – distal-less homeobox 5 | Depew *et al*. (2002), Graham (2002), Jeong *et al*. (2008) (133–135) |
| *DLX6* – distal-less homeobox 6 | Depew *et al*. (2002), Graham (2002), Jeong *et al*. (2008) (133–135) |
| *Dlx6os1* – distal-less homeobox 6, opposite strand 1 | In Jeong *et al*. (2008) as *Evf1/2* (135) |
| *GBX2* – gastrulation brain homeobox 2 | Jeong *et al*. (2008) (135) |
| *GSC* – goosecoid homeobox | Jeong *et al*. (2008) (135) |
| *HAND1* – heart and neural crest derivatives expressed 1 | Jeong *et al*. (2008) (135) |
| *HAND2* – heart and neural crest derivatives expressed 2 | Jeong *et al*. (2008) (135) |
| *HGF* – hepatocyte growth factor | Jeong *et al*. (2008) (135) |
| *MEIS2* – Meis homeobox 2 | Machon *et al*. (2015) (137) |
| *NKX3-2* – NK3 homeobox 2 | In Square *et al*. (2020) as *NKX3.2* (139) |
| *PLAGL1* – PLAG1 like zinc finger 1 | In Jeong *et al*. (2008) as *Zac1* (135) |
| *RGS5* – regulator of G protein signaling 5 | Jeong *et al*. (2008) (135) |
| *UNC5C* – unc-5 netrin receptor C | Jeong *et al*. (2008) (135) |

**Table S2.**

Genes expressed in the gnathostome jaw and queried for the initial relative evolutionary analysis dataset. Note that not all genes contained adequate sample sizes (>100 spp) to be included in the subsequent RER analyses.

| Gene | Total Seqs. Recovered from BLAST | # Vetted sequences with confirmed identity | # Vetted sequences with “[name]-like” excluded | # Vetted sequences after synonymizing | # final sequences after pruning to one longest sequence per taxon |
|---|---|---|---|---|---|
| *ALX3* | 5000 | 368 | n/a | n/a | 298 |
| *BMPER* | 899 | 647 | n/a | 650 | 345 |
| *CITED1* | 1994 | 777 | n/a | 781 | 253 |
| *DLX1* | 4418 | 632 | 608 | n/a | 377 |
| *DLX2* | 5000 | 990 | 948 | 949 | 514 |
| *DLX3* | 5000 | 519 | 399 | n/a | 363 |
| *DLX4* | 4782 | 529 | 358 | 363 | 273 |
| *DLX5* | 4806 | 465 | 449 | n/a | 384 |
| *DLX6* | 3825 | 471 | 450 | n/a | 394 |
| *Dlx6os1* | 5000 | 3 | 3 | 6 | 3 |
| *GBX2* | 4441 | 435 | n/a | n/a | 330 |
| *GSC* | 1417 | 383* | n/a | n/a | 343 |
| *HAND1* | 2838 | 389 | n/a | 393 | 292 |
| *HAND2* | 3271 | 395 | n/a | 396 | 332 |
| *HGF* | 2025 | 848 | n/a | n/a | 335 |
| *MEIS2* | 5000 | 2794 | 2620 | 2622 | 372 |
| *NKX3-2* | 5000 | 332 | n/a | 343 | 312 |
| *PLAGL1* | 5000 | 1858 | n/a | 1865 | 246 |
| *RGS5* | 5000 | 339 | n/a | 343 | 263 |
| *UNC5C* | 4579 | 1039 | 692 | n/a | 335 |

*This amount after excluding 291 sequences of the paralog GSC2.

**Table S3.**

Number of resulting sequences (seqs.) after different filters were applied to the datasheet.

| Gene | Alternative Name | # Seq | Taxon |
|---|---|---|---|
| *ALX3* | *Fnd*<br>*Fnd1* | 0 | n/a |
| *BMPER* | *Cv2* | 1 | *Mus musculus* |
| | *Cv-2* | 2 | *Homo sapiens*<br>*Xenopus tropicalis* |
| | *Crim3* | 0 | n/a |
| *CITED1* | *MSG1* | 4 | *Bos Taurus*<br>*H. sapiens*<br>*M. musculus*<br>*Rattus norvegicus* |
| *DLX1* | n/a | | |
| *DLX2* | TES-1 | 1 | *M. musculus* |
| | TES1 | 0 | n/a |
| *DLX3* | *AI4*<br>*TDO* | 0 | n/a |
| *DLX4* | BP1 | 1 | *H. sapiens* |
| | DLX7 | 3 | *H. sapiens* (2)<br>*M. musculus* |
| | DLX8=1 | 1 | *Danio redio* |
| | DLX9<br>OFC15 | 0 | n/a<br>n/a |
| *DLX5* | *SHFM1*<br>*SHFM1D* | 0 | n/a |
| *DLX6* | n/a | | |
| *Dlx6os1* | Dlx6 | 1 | *Geopiza fortis* |
| | Evf1 | 1 | *R. norvegicus* |
| | Evf2 | 1 | *R. norvegicus* |
| | Dlx6as1<br>Evf<br>Evf-<br>Evf-1<br>Evf-2<br>Evf1/<br>Evf1/2<br>Sh<br>Shhrs<br>mEvf- | 0 | n/a |
| *GBX2* | n/a | | |
| *GSC* | SAMS | 0 | n/a |
| *HAND1* | Hxt | 1 | *M. musculus* |
| | Thing1 | 1 | *M. musculus* |
| | eHand | 2 | *Oryctolagus cuniculus*<br>*R. norvegicus* |

| | | | |
|---|---|---|---|
| | bHLHa27 | 0 | n/a |
| *HAND2* | *Hed* | 1 | *M. musculus* |
| | *DHAND2*<br>*Thing2*<br>*bHLHa26*<br>*dHand* | 0 | n/a |
| *HGF* | *DFNB39*<br>*F-TCF*<br>*HGFB*<br>*HPTA*<br>*SF* | 0 | n/a |
| *MEIS2* | MRG1 | 2 | *M. musculus* (Mrg1a and Mrg1b) |
| *NKX3-2* | *BAPX1* | 9 | *H. sapiens* (4)<br>*L. camtschaticum* (1)<br>*M. musculus* (1)<br>*Petromyzon* marinus (1)<br>*Scyliorhinus canicular* (1)<br>*Scyliorhinus torazame* (1) |
| | NKX3.2 | 2 | *Gallus gallus* |
| | NKX3B<br>SMMD | 0 | n/a |
| *PLAGL1* | LOT1 | 2 | *H. sapiens*<br>*M. musculus* |
| | ZAC | 4 | *H. sapiens* (3)<br>*Bos taurus* |
| | ZAC1 | 1 | *M. musculus* |
| *RGS5* | MSTP032 | 1 | *H. sapiens* |
| | MSTP092 | 1 | *H. sapiens* |
| | MSTP106 | 1 | *H. sapiens* |
| | MSTP129 | 1 | *H. sapiens* |
| | MST092<br>MST106<br>MST129 | 0 | n/a |
| *UNC5C* | UNC5H3 | 0 | n/a |

**Table S4.**
Synonyms of genes as recorded in GenBank and number of sequences for gene and taxon.

| Gene | Seqs. With scientific name missing | Presumed taxon | Kept (K)/ Deleted (D) | Justification |
|---|---|---|---|---|
| *ALX3* | n/a | | | |
| *BMPER* | n/a | | | |
| *CITED1* | 8 | *Mauremys reevesii* XM_039487110.1 | D | It was not the longest sequence for the species |
| | | *M. reevesii* XM_039487111.1 | D | It was not the longest sequence for the species |
| | | *M. reevesii* XM_039487108.1 | K | It was the longest sequence for the species and there was not a record properly filled out |
| | | *Pipra filicauda* | K | It was the only specimen for the specie and genus |
| | | *Saimiri boliviensis boliviensis* XM_010332397.2 | D | It was not the longest sequence for the species |
| | | *S. boliviensis. b.* XM_010332395.2 | D | It was not the longest sequence for the species |
| | | *S. boliviensis. b.* XM_039475681.1 | D | It was not the longest sequence for the species |
| | | *S. boliviensis. b.* XM_039475680.1 | K | It was the longest sequence for the species and there was not a record properly filled out |
| *DLX1* | 1 | *Pipistrellus kuhlii* | D | It was not the longest sequence for the species |
| *DLX2* | 8 | *Scarus compressus* | D | There was other record properly filled out with the same length |
| | | *Sc. ghobban* | D | It was not the longest sequence for the species |
| | | *Sc. perrico* | D | There was other record properly filled out with the same length |
| | | *Sc. rubroviolaceus* | D | It was not the longest sequence for the species |
| | | *Crotalus tigris* | K | It was the only specimen for the specie. |
| | | *M. reevesii* | K | It was the only specimen for the specie. |

| | | | | |
|---|---|---|---|---|
| | | *P. filicauda* | K | It was the only specimen for the specie. |
| | | *S. boliviensis b.* | K | It was the only specimen for the specie. |
| *DLX3* | n/a | | | |
| *DLX4* | n/a | | | |
| *DLX5* | n/a | | | |
| *DLX6* | 3 | *Acipenser ruthenus* | D | It was not the longest sequence for the species |
| | | *P. filicauda* | K | It was the only specimen for the specie. |
| | | *S. boliviensis b.* | K | It was the only specimen for the specie. |
| *GBX2* | n/a | | | |
| *GSC* | | *Corvus cornix cornix* | K | It was the only specimen for the specie |
| | | *Cr. tigris* | K | It was the only specimen for the specie |
| | | *M. reevesii* | K | It was the only specimen for the specie |
| | | *P. filicauda* | K | It was the only specimen for the specie |
| *HAND1* | n/a | | | |
| *HAND2* | n/a | | | |
| *HGF* | n/a | | | |
| *MEIS2* | n/a | | | |
| *KX3-2* | n/a | | | |
| *PLAGL1* | | *C. cornix c.* | K | It was the only specimen for the specie |
| | | *M. reevesii* | K | It was the only specimen for the specie |
| | | *P. filicauda* | K | It was the only specimen for the specie |
| | | *S. boliviensis b.* | K | It was the only specimen for the specie |
| *RGS5* | n/a | | | |
| *UNC5C* | | *C. cornix c.* | K | It was the only specimen for the specie |
| | | *P. filicauda XM_027731747.1* | D | It was not the longest sequence for the species |
| | | *P. filicauda* XM_027731746.1 | K | It was the longest sequence for the species and there was not a record properly filled out |

| | | |
|---|---|---|
| *S. boliviensis b.* *XM_003929536.2* | D | It was not the longest sequence for the species |
| *S. boliviensis b.* XM_010340136.1 | K | It was the only specimen for the specie |

**Table S5.**
Presumed scientific name of sequences for each included gene that did not have a properly annotated scientific name field.

| residual transform-weight-scale | PC1 | PC2 | Stiffness | AR | MA |
|---|---|---|---|---|---|
| none-F-F | bmp3,plagl1,alx3,rgs5,dlx2 | none | **alx3,rgs5**,bmp3 | **alx3**,bmp3,rgs5,plagl1 | none |
| none-T-F | none | none | none | **alx3**,bmp3 | **dlx1** |
| none-T-T | none | none | none | none | none |
| none-F-T | none | none | none | none | none |
| sqrt-F-F | none | none | **rgs5,alx3** | **alx3**,bmp3,dlx2 | **dlx1** |
| sqrt-T-F | none | none | none | **alx3** | **dlx1** |
| sqrt-T-T | none | none | none | none | none |
| sqrt-F-T | none | none | none | none | none |
| log-F-F | none | none | none | none | none |
| log-T-F | none | none | none | **alx3** | **dlx1** |
| log-T-T | none | none | none | none | none |
| log-F-T | none | none | none | none | none |

**Table S6.**
Relative evolutionary rate sensitivity analysis with different transformation parameters.

Non-tetrapod-Tetrapoda Comparison

| | Non-tetrapod | Tetrapods |
|---|---|---|
| Non-tetrapod | - | 7950 |
| Tetrapods | 0.899 | - |

Within-tetrapod Comparison

| | Synapsids | Sauropsids | Amphibia | Stem Tetrapodomorphs |
|---|---|---|---|---|
| Synapsids | - | 15 | 34 | 7466 |
| Sauropsids | **0.002** | - | 1774 | 164 |
| Amphibia | **0.004** | 0.201 | - | 0 |
| Stem Tetrapodomorphs | 0.844 | **0.019** | 0 | - |

Within-sauropsids vs. non-sauropsids

| | Non-sauropsids | Lepidosaurs | Ichthyosaurs | Archosaurs | Testudines |
|---|---|---|---|---|---|
| Non-sauropsids | - | 110 | 0 | 0 | 7755 |
| Lepidosaurs | **0.012** | - | 6437 | 5239 | 82 |
| Ichthyosaurs | 0 | 0.728 | - | 6775 | 0 |
| Archosaurs | 0 | 0.592 | 0.766 | - | 0 |
| Testudines | 0.877 | **0.009** | 0 | 0 | - |

Archosauromorpha

| | Crocodylia/ Loricata | Pterosauria | Non-avian Dinosauria | Neornithes | Stem Archosauromorphs |
|---|---|---|---|---|---|
| Crocodylia/Loricata | - | 7985 | 3018 | 4471 | 7292 |
| Pterosauria | 0.903 | - | 2733 | 3973 | 6542 |
| Non-avian Dinosauria | 0.341 | 0.309 | - | 3615 | 3602 |
| Neornithes | 0.506 | 0.449 | 0.409 | - | 5812 |
| Stem Archosauromorphs | 0.825 | 0.74 | 0.407 | 0.657 | - |

Within-synapsids vs. non-synapsids

| | Cynodonta | Pelycosaurs | Mammaliaformes | Non-Synapsids |
|---|---|---|---|---|
| Cynodonta | - | 6240 | 8202 | 5868 |
| Pelycosaurs | 0.706 | - | 6117 | 8334 |
| Mammaliaformes | 0.928 | 0.692 | - | 5742 |
| Non-Synapsids | 0.664 | 0.942 | 0.649 | - |

**Table S7.**

Summary statistics for differences in adaptive landscape optimizations between pairwise clade groups. Lower diagonal values are *p* values; upper diagonal values are goodness of fit. Bold fonts indicate statistical significance at $p < 0.02$.

**Movie S1. (separate file)**
Movie_S1_model21_10-100N_8-13-21_Step00-68.avi
10.6084/m9.figshare.21232022

**Data S1. (separate file)**
Data_S1_Supplemental_Table_Specimen_List.xlsx
10.6084/m9.figshare.21232079

**Data S2. (separate file)**
Data_S2_Specimen_Usage_Permission_Documents.zip
10.6084/m9.figshare.21232082

**Data S3. (separate file)**
Data_S3_Tsengetal_jaw_shape_outlines_n1352.zip
10.6084/m9.figshare.21232055Data S4. (separate file)
Data_S4_Tsengetal_EFA_datasets_R_objects.zip
10.6084/m9.figshare.21232058

**Data S5. (separate file)**
Data_S5_Supplemental_Table_PC_Scores_Labeled_All.csv
10.6084/m9.figshare.21232043

**Data S6. (separate file)**
Data_S6_Supplemental_Table_theoretical_models_MA&AR.csv
10.6084/m9.figshare.21232049

**Data S7. (separate file)**
Data_S7_Supplemental_Table_theoretical_models_FEA_output.xlsx
10.6084/m9.figshare.21232085

**Data S8. (separate file)**
Data_S8_Tsengetal_theoretical_jaw_FE_models.zip
10.6084/m9.figshare.21232067

**Data S9. (separate file)**
Data_S9_Supplemental_Table_bending_test_validation_data.xlsx
10.6084/m9.figshare.21232073

**Data S10. (separate file)**
Data_S10_Supplemental_Table_Wilcoxon_Test_by_Clade_Displacement&SE.xlsx
10.6084/m9.figshare.21232052

**Data S11. (separate file)**
Data_S11_Supplemental_Table_Genera_List.xlsx
10.6084/m9.figshare.21232037

**Data S12. (separate file)**
Data_S12_Supplemental_Table_Fossil_Topology_Source.xlsx
10.6084/m9.figshare.21232061

**Data S13. (separate file)**
Data_S13_Supplemental_File_Extant_Genus_Tree.pdf
10.6084/m9.figshare.21232070

**Data S14. (separate file)**
Data_S14_Supplemental_Table_Extant_Gnathostome_Diversity.xlsx
10.6084/m9.figshare.21232064

**Data S15. (separate file)**
Data_S15_Tsengetal_composite_tree_samples&subsamples.zip
10.6084/m9.figshare.21232076

**Data S16. (separate file)**
Data_S16_Supplemental_Table_BLAST_Parameters.xlsx
10.6084/m9.figshare.21232028

**Data S17. (separate file)**
Data_S17_Supplemental_Table_RER_Gene_Sequence_Accessions.xlsx
10.6084/m9.figshare.21232046

**Data S18. (separate file)**
Data_S18_Supplemental_Table_Gene_Tree_Entanglement.xlsx
10.6084/m9.figshare.21232034

**Data S19. (separate file)**
Data_S19_Supplemental_Table_Mammal_Body_Mass_PanTHERIA.xlsx
10.6084/m9.figshare.21232040

**Data S20. (separate file)**
Data_S20_Supplemental_Table_body-mass&adductor-mass.xlsx
10.6084/m9.figshare.21232031

**Data S21. (separate file)**
Data_S21_Figure_silhouette_files&attributions.zip
10.6084/m9.figshare.21232025
